# CLN8 enables a non-canonical phospholipid synthesis pathway

**DOI:** 10.64898/2025.12.18.693953

**Authors:** Johannes Breithofer, Nermeen Fawzy, Clara Zitta, Martin Tischitz, Dominik Bulfon, Clemens Hofmann, Lennart Hartig, Carina Wagner, Gernot F. Grabner, Anita Pirchheim, Achim Lass, Ulrike Taschler, Keira Turner, Kasparas Petkevicius, Ulrich Stelzl, Dagmar Kratky, Rolf Breinbauer, Robert Zimmermann

**Affiliations:** Institute of Molecular Biosciences, University of Graz, Graz, Austria; Institute of Organic Chemistry, Graz University of Technology, Graz, Austria; Gottfried Schatz Research Center, Molecular Biology and Biochemistry, Medical University of Graz, Graz, Austria; MRC Mitochondrial Biology Unit, University of Cambridge, Cambridge Biomedical Campus, Cambridge, UK; Institute of Pharmaceutical Sciences, Pharmaceutical Chemistry, University of Graz, Graz, Austria; Cell Death and Metabolism Unit, Danish Cancer Institute, Copenhagen, Denmark; BioTechMed-Graz, Graz, Austria; Field of Excellence BioHealth, University of Graz, Graz, Austria

## Abstract

According to text book knowledge, *de novo* glycerophospholipid (GPL) synthesis begins with the acylation of glycerol-3-phosphate to form phosphatidic acid, the precursor of all other GPLs. Here we describe an alternative GPL synthesis pathway that starts with the acyl-CoA-dependent acylation of glycerophosphoglycerol (GPG), resulting in the formation of lysophosphatidylglycerol (LPG). The acyltransferase reaction is catalyzed by the Batten disease-associated protein ceroid lipofuscinosis neuronal 8 (CLN8). Tracer studies revealed that CLN8-derived LPG is selectively converted into bis(monoacylglycero)phosphate (BMP), a GPL essential for lysosomal lipid homeostasis, but not into phosphatidylglycerol or cardiolipin. *CLN8*-knockout cells and mice cannot utilize GPG for BMP synthesis, resulting in BMP-deficiency and excess accumulation of phospholipids in lysosomes. The lipid synthesis pathway described herein is relevant for understanding lysosomal lipid metabolism and the pathogenesis of neurodegenerative diseases. BMP-deficiency may contribute to or even underlie lysosomal cargo accumulation in certain forms of Batten disease and other lysosomal storage disorders.

## Main

Membranes of different organelles are characterized by defined lipid and protein compositions, which determine their function. The lipid composition affects membrane fluidity and curvature, and is essential for folding and activity of membrane-associated proteins. Lumenal membranes of late endosomes and lysosomes contain bis(monoacylglycero)phosphate (BMP), also known as lysobisphosphatidic acid, as lipidomic signature^1^. BMP plays a central role in lysosomal lipid homeostasis. It is highly resistant to lysosomal hydrolases^2^, promotes the formation of lumenal membranes^3^, and acts as a cofactor of lysosomal lipid hydrolases and transport proteins that mediate the breakdown and export of lipids^4^.

BMP is a structural isomer of phosphatidylglycerol (PG), indicating that PG acts as precursor of BMP. However, these lipids differ in their stereo- and acyl chain configuration. PG exhibits *sn-3*-glycerophosphate-*sn-1*-glycerol (*R,S*) backbone configuration and is acylated exclusively at the *R*-bound glycerol. Conversely, BMP has an *S,S*-backbone configuration and contains a single fatty acid (FA) group at each glycerol residue^5–7^, implicating that the conversion of PG into BMP requires both stereoconversion and positional rearrangement of FAs.

Previous studies suggest that PG is converted into BMP within lysosomes. This process requires the activity of lysosomal hydrolases degrading PG to lysophosphatidylglycerol (LPG)^8^, followed by the conversion of LPG into BMP via transacylation and transphosphatidylation reactions^9,10^. Here, we show that the synthesis of BMP begins at the endoplasmic reticulum (ER) via an unusual reaction that is independent of established *de novo* GPL synthesis pathways starting with the acylation of glycerol-3-phosphate^11,12^. The ER-associated protein ceroid lipofuscinosis neuronal 8 (CLN8) catalyzes the acyl-CoA-dependent acylation of glycerophosphoglycerol (GPG), producing LPG, which is selectively converted into BMP. *CLN8*-deficient cells and mice lack GPG acyltransferase activity and BMP. Mutations in the *CLN8* gene are linked to the neurodegenerative disorder Batten disease^13–16^, characterized by the accumulation of autofluorescent lipopigments (ceroid lipofuscin) in lysosomes of neurons and other cells. Our observations suggest that the lack of BMP is causal for the lysosomal storage disorder in *CLN8*-deficient cells.

## Results

### GPG is used for LPG and BMP synthesis

The deacylation of GPLs by (lyso)phospholipases A1/A2 generates glycerophosphodiesters (GPDs), including GPG, which constitutes the backbone of PG and BMP. It is currently thought that GPDs are degraded by glycerophosphodiesterases, thereby providing building blocks for *de novo* lipid synthesis and signaling molecules^17^. To investigate whether mammalian cells can reuse GPG for GPL synthesis, we incubated lysates of human cell lines and mouse tissues with GPG containing ^13^C-labeled glycerol (*sn*-[U-^13^C]glycero-1(3)-phospho-*sn*-3-glycerol (*rac*,*R*); *m/z* +3; [^13^C]GPG^+3^) in the presence and absence of oleoyl-CoA (**Fig. 1a**). These experiments revealed that [^13^C]GPG^+3^ is acylated in an acyl-CoA-dependent manner in the presence of lysates of human cell lines and mouse tissues (**Fig. 1b, c**). Among the tested mouse tissues, the formation of ^13^C-labeled LPG ([^13^C]LPG^+3^) was highest in brain lysates. Thus, mammalian cells and tissues contain acyl-CoA:GPG acyltransferase (GPGAT) activity.

**Figure 1.**
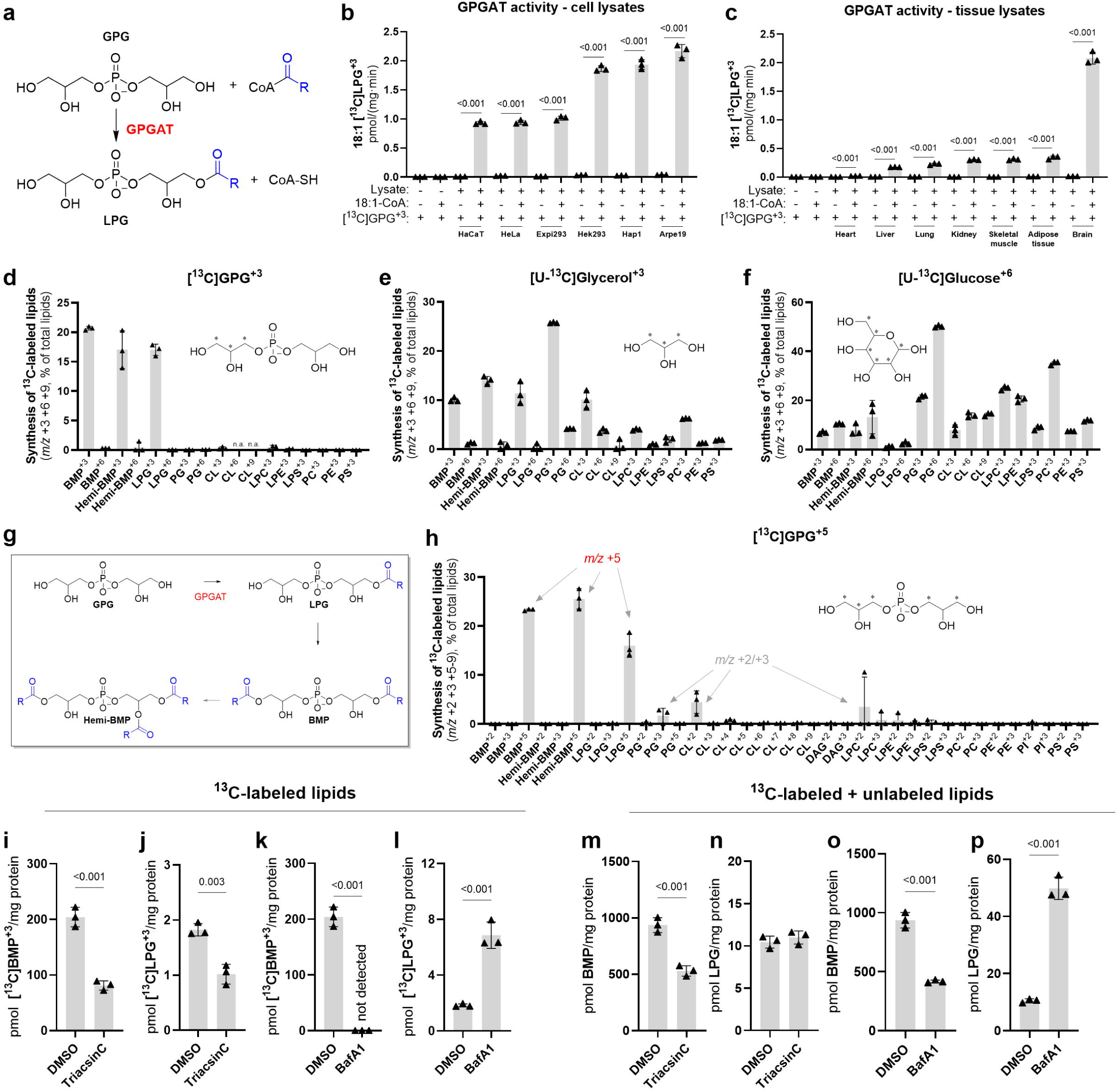
Mammalian cells utilize GPG for phospholipid synthesis. **a,** Schematic representation of the acyl-CoA:GPG acyltransferase (GPGAT) reaction. **b,c,** GPGAT activity in lysates (1.000 x *g* supernatant) of the indicated human cell lines (b) and mouse tissues (c) was detected using 100 µM [^13^C]GPG^+3^ and oleoyl-CoA as substrate. Assays were conducted in 100 mM bis-tris-propane buffer (pH 7) supplemented with 1% FFA free BSA and 1 mM DTT. [^13^C]LPG^+3^ formation was analyzed by LC-MS (n=3). **d-f,** Incorporation of [^13^C]GPG^+3^ (500 µM), [U-^13^C]glycerol^+3^ (500 µM), and [U-^13^C]glucose^+6^ (25 mM) into phospholipids of Hek293 cells (*indicates ^13^C-atoms). In f, glucose of the medium was replaced by [U-^13^C]glucose^+6^. Incubations were performed for 16h under standard cell culture conditions. The results are presented as a percentage of labeled lipids relative to total lipids for each lipid class examined (n=3, data are representative of two independent experiments). **g,** Schematic representation of GPG-dependent BMP and hemi-BMP synthesis. **h,** Incorporation of [^13^C]GPG^+5^ (250 µM) into lipids of Hek293 cells. Experiments were performed as described in d-f (n=3). **i-l,** Effect of Triacsin C (10 µM; i,j) and Bafilomycin A1 (BafA1, 0.1 µM; k,l) on the incorporation of [^13^C]GPG^+3^ (500 µM) into LPG and BMP. Incubations were performed for 18h in the presence of inhibitors or DMSO as vehicle control (n=3). **m-p,** Effect of Triacsin C and BafA1 on endogenous BMP and LPG levels of Hek293 cells (labeled plus unlabeled). Experiments were performed as described in i-l (n=3). Labeled and unlabeled lipid species were analyzed with LC-MS and are presented as sum of analyzed subspecies. Data are presented as mean ± SD. Statistical comparisons in b,c,i-p were performed with unpaired two-tailed Student’s *t* test. *P*-values are indicated.

To track the distribution of GPG within phospholipid species, we incubated Hek293 cells with [^13^C]GPG^+3^ and monitored its incorporation into GPLs. Mass spectrometry analysis revealed that GPG is preferentially incorporated into LPG, hemi-BMP, and BMP. Hemi-BMP, also known as acyl-phosphatidylglycerol, is considered as intermediate in the synthesis of BMP^18^. Of the analyzed BMP, LPG, and hemi-BMP, 21%, 17%, and 17%, showed a shift in the mass-to-charge ratio *(m/z)* of +3, respectively. (**Fig. 1d**). In comparison, no substantial labeling was detected for other polyglycerophospholipids, such as PG and cardiolipin (CL), as well as other GPL classes (**Fig. 1d**). The degradation of GPG by phosphodiesterases generates glycerol-3-phosphate and glycerol, which can be reused for *de novo* synthesis of PA, the precursor of all GPLs, as well as for the head group synthesis of polyglycerophospholipids. Accordingly, we also monitored phospholipids with a *m/z* shift of +6 for LPG, PG, BMP, and hemi-BMP. However, we could not detect species containing more than one labeled glycerol moiety (**Fig. 1d**). In comparison, incubation of cells with [U-^13^C]glycerol^+3^ or [U-^13^C]glucose^+6^, both delivering glycerol-3-phosphate for *de novo* lipid synthesis, produced a different labeling pattern (**Fig. 1e,f**). As expected, these metabolites were incorporated into all detected GPL classes. For polyglycerophospholipids, we also detected species containing more than one labeled glycerol *(m/z* shift of +6 and +9), suggesting that the ^13^C-labeled glycerol was used for PA and head group synthesis. The differences in the GPL labeling patterns between cells incubated with ^13^C-labeled GPG or glycerol/glucose indicate that (i) GPG enters a specific metabolic pathway leading to the formation of LPG, BMP, and hemi-BMP (**Fig. 1g**) and that (ii) GPG is incorporated into GPLs without prior degradation, likely via the GPGAT reaction.

To confirm direct acylation of GPG, we synthesized GPG containing ^13^C-labeled carbon atoms on both glycerol moieties (*sn*-[U-^13^C]glycero-1(3)-phospho-*sn*-1(3)-[1,3-^13^C]-glycerol (*rac*,*rac*); *m/z* +5; [^13^C]GPG^+5^) and again monitored its incorporation into lipids of Hek293 cells. Direct acylation of this tracer results in lipids with a *m/z* value of +5, whereas degradation of GPG and reuse of labeled glycerol for lipid synthesis leads to lipid species with *m/z* values of +2 or +3. We detected exclusively the formation of labeled BMP, hemi-BMP, and LPG with *m/z* +5 (**Fig. 1h**), confirming direct acylation of GPG. Minor labeling of other GPL species with *m/z* of +2 or +3, suggest that [^13^C]GPG^+5^ is partially degraded and reutilized for PA or headgroup synthesis (**Fig. 1h**).

To characterize the cellular pathways that mediate the incorporation of GPG into phospholipids, we treated Hek293 cells with Triacsin C, an inhibitor of acyl-CoA synthetases 1 and 4^19^, and Bafilomycin A1 (BafA1), an inhibitor of lysosomal V-ATPase, which leads to alkalization of lysosomes^20^. Triacsin C reduced the incorporation of [^13^C]GPG^+3^ into BMP and LPG by 60 and 40%, respectively (**Fig. 1i, j**). This observation indicates that the availability of acyl-CoAs affects both GPG-dependent LPG and BMP synthesis. BafA1 completely abolished [^13^C]BMP^+3^ synthesis (**Fig. 1k**), while [^13^C]LPG^+3^ levels increased three-fold (**Fig. 1l**), suggesting that LPG synthesis occurs outside of acidic organelles, whereas BMP synthesis requires functional lysosomes. The alterations observed in tracer studies were partially reflected by changes in total cellular lipid content. Triacsin C reduced BMP levels by 40%, while LPG remained unchanged (**Fig. 1m,n**). In the presence of BafA1, BMP content decreased by 55% and LPG levels increased five-fold (**Fig. 1o,p**).

### CLN8 exhibits GPGAT activity

To gain insights into the molecular mechanisms mediating GPG acylation, we first performed a rough separation of cell lysates (1,000 x *g* supernatant) into a mitochondria-enriched fraction (10,000 x *g* pellet), a microsomal fraction (100,000 x *g* pellet), and a cytosolic fraction (100,000 x *g* supernatant). GPGAT activity was observed exclusively in the membrane-containing fractions, suggesting that this reaction is catalyzed by a membrane-bound enzyme (**Fig. 2a**).

**Figure 2.**
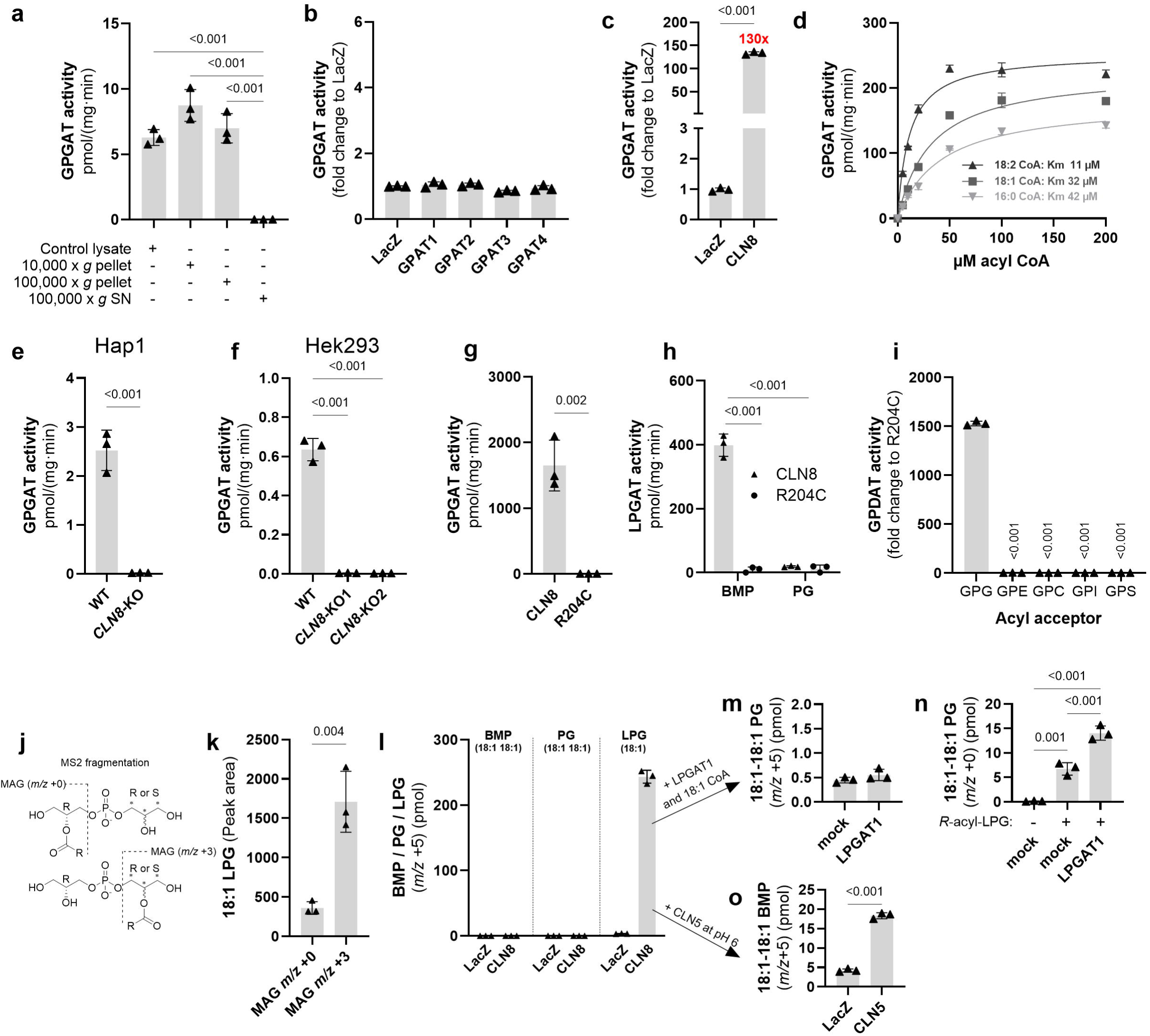
CLN8 exhibits GPGAT activity. **a,** GPGAT activity detected in 1,000 x *g* supernatant (SN), 10,000 x *g* pellet, 100,000 x *g* pellet, and the 100,000 x *g* SN fraction of Expi293 cells (n=3). **b,** Changes in GPGAT activity detected in Expi293 cell lysates (1,000 x *g* SN) overexpressing human GPAT1-4 in comparison to the LacZ control (n=3). **c,** Change in GPGAT activity detected in lysates of Expi293 cells overexpressing human CLN8 in comparison to the LacZ control (n=3, data are representative of three independent experiments). **d,** GPGAT activity in CLN8-overexpressing Expi293 lysates in the presence of [^13^C]GPG^+3^ as acyl acceptor and increasing concentrations of 16:0-, 18:1-, and 18:2-CoAs as acyl donor (n=3). **e,f,** GPGAT activity detected in lysates of WT and *CLN8*-deficient Hap1 or Hek293 cells (*CLN8*-KO). Two *CLN8*-KO Hek293 cell clones were analyzed (n=3). **g,** GPGAT activity of purified His-tagged CLN8 and its mutant variant R204C (1 µg each; n=3, data are representative of two independent experiments). **h,** CLN8-catalyzed formation of BMP and PG by purified CLN8 and R204C (1 µg each) using *sn-*1-oleoyl-*R,rac* LPG as acyl acceptor (n=3). **i,** Head group specificity of purified CLN8 was determined by the formation of lysophospholipids in the presence of oleoyl-CoA and the indicated GPDs (100 µM each). The inactive R204C variant was used as negative control (n=3). **j,** Schematic representation showing MS2 fragmentations of LPG in *R,rac*-[^13^C]-GPG^+3^ backbone configuration. Depending on the position of the acyl-chain, the monoacylglycerol (MAG) fragments show *m/z* values of +0 or +3. **k,** Analysis of MAG fragments, as depicted in j, of oleoyl-[^13^C]LPG^+3^ generated by purified CLN8. [^13^C]GPG^+3/PG^, derived from headgroup labeled PG, and oleoyl-CoA were used as substrate (n=3). **l,** Formation of [^13^C]BMP/PG/LPG^+5^ by sonicated membrane fractions (100,000 x *g* pellet) of LacZ and CLN8 expressing Expi293 cells using 100 µM [^13^C]GPG^+5^ and 100 µM of oleoyl-CoA as substrate (n=3). **m,** Conversion of CLN8-derived [^13^C]LPG^+5^ into [^13^C]PG^+5^: After heat inactivation, CLN8-derived [^13^C]LPG^+5^ was further incubated with membranes of cells overexpressing recombinant LPGAT1 and oleoyl-CoA (6 µM). Membranes of mock-transfected cells were used as a control. (n=3) **n,** Conversion of *R*-acyl LPG into PG. The experiment was performed as described in m using *R*-acyl-LPG (0.6 µM) and oleoyl-CoA (6 µM) as acyl acceptor and donor, respectively (n=3). **o,** Conversion of CLN8-derived [^13^C]LPG^+5^ into [^13^C]BMP^+5^. After heat inactivation, CLN8-derived [^13^C]LPG^+5^ was further incubated with lysates of cells overexpressing recombinant CLN5 at pH 6.0. LacZ-transfected cell lysates were used as a control (n=3). Formation of lipids was analyzed by LC-MS. Data are presented as mean ± SD. Statistical comparisons in a,f,i,n were performed with one-way ANOVA followed by Bonferroni posthoc analysis and in c,e,g,h,k,o with unpaired two-tailed Student’s *t* test. *P*-values are indicated.

Mammalian cells express four acyl-CoA:glycerol-3-phosphate acyltransferases (GPAT1-4). GPAT1 and GPAT2 are localized in the outer mitochondrial membrane, while GPAT3 and GPAT4 are found in the ER^21^. To test whether human GPATs possess GPGAT activity, we expressed these acyltransferases in Expi293 cells (**Extended data fig. 1a)** and measured GPGAT activity in lysates. However, we found no change in comparison to control cells the express β-galactosidase (LacZ) (**Fig. 2b**), implicating that GPG enters a GPAT-independent lipid synthesis pathway.

Recent observations showed that CLN8, a member of the TRAM-LAG1-CLN8 domain-containing protein family, exhibits acyl-CoA:LPG acyltransferase (LPGAT) activity and strongly affects BMP biosynthesis^22^. Strikingly, overexpression of CLN8 in Expi293 cells (**Extended data fig. 1b**) increased GPGAT activity 130-fold (**Fig. 2c**). Saturation kinetics with different acyl-CoAs revealed a dose-dependent increase in GPGAT activity and lower *K_m_* values for linoleoyl- and oleoyl-CoA than for palmitoyl-CoA (**Fig. 2d**). These observations suggest that CLN8 preferentially uses acyl-CoAs with polyunsaturated FAs, similar as previously described for the LPGAT reaction^22^. CRISPR/Cas9-mediated deletion of *CLN8* in Hap1 and Hek293 cells completely abolished GPGAT activity (**Fig. 2e,f**). Mutations in the *CLN8* gene and depletion of CLN8 protein levels were confirmed via Sanger sequencing and proteomics, respectively (**Extended data fig. 2a,b**).

GPGAT activity of CLN8 was confirmed with the purified enzyme (**Fig. 2g, Extended data fig. 1c**). The purified mutant CLN8 variant R204C, which is associated with Batten disease^15,16^, showed no activity (**Fig. 2g**). Wild-type (WT) CLN8, but not the R204C variant, also catalyzed the LPGAT reaction, but at an ∼ four-fold lower rate than GPG acylation (**Fig. 2g,h**). The LPGAT reaction generated exclusively BMP, suggesting that CLN8 mediates head group acylation of LPG (**Fig. 2h**). Purified CLN8 did not acylate other GPDs than GPG, suggesting that the enzyme has pronounced head group specificity (**Fig. 2i**).

To further investigate the stereospecificity of CLN8, we isolated GPG from head group labeled PG and used this probe ([^13^C]GPG^+3/PG^) as acyl acceptor for the CLN8-catalyzed acyltransferase reaction. The generated LPG was analyzed by MS2 fragmentation analysis. Depending on the acylation of the unlabeled *R-*linked glycerol or the ^13^C-labeled head group, fragmentation of LPG can lead to the formation of either unlabeled or ^13^C-labeled monoacylglycerol species, respectively (**Fig. 2j**). We observed mostly labeled monoacylglycerol fragments, suggesting that CLN8 preferentially acylates the *S*-linked glycerol moiety (**Fig 2j,k)**.

Our tracer studies suggested that GPG is not directly incorporated into PG and cardiolipin (**Fig. 1d,h**), which may either be due to spatial separation of LPG and PG synthesis or due to the stereospecific acylation of LPG at the *S*-position. To exclude spatial separation, we monitored lipid synthesis in sonicated membrane preparations of Expi293 cells. CLN8-containing membrane vesicles efficiently produced [^13^C]LPG^+5^ in the presence of [^13^C]GPG^+5^ and oleoyl-CoA, but no substantial amounts of [^13^C]BMP^+5^ or [^13^C]PG^+5^ (**Fig. 2l**). Further incubation of CLN8-derived LPG with membranes containing the recombinantly expressed ER-resident LPG acyltransferase 1^23^ (LPGAT1, **Extended data fig. 1d**) did not increase [^13^C]PG^+5^ formation (**Fig. 2m**). In contrast, supplementation of *R*-acylated LPG (*sn*-1-oleoyl-*R,rac* LPG) enabled the formation of PG by membranes of mock-transfected cells, which was two-fold elevated in the presence of membranes containing recombinant LPGAT1 (**Fig. 2n**). Thus, LPGAT1 exhibits stereoselectivity for *R*-acyl-LPG and cannot convert CLN8-derived *S*-acyl-LPG into PG.

To investigate whether the CLN8 product can be used for BMP synthesis, we monitored the conversion of CLN8-derived LPG into BMP at pH 6.0, the pH optimum of the lysosomal BMP synthase CLN5^9^. This enzyme mediates a transacylation reaction in which LPG acts as both FA donor and acceptor, resulting in the formation of BMP and GPG^9^. Under the applied conditions, the formation of [^13^C]BMP^+5^ by cell lysates containing recombinant CLN5 was increased four-fold compared to LacZ controls (**Fig. 2o**). Together, these observations suggest that the stereospecific acylation of GPG by CLN8 at the *S*-linked glycerol moiety prevents PG formation and directs CLN8-derived LPG into BMP synthesis.

### *CLN8*-deficient cells lack BMP

To gain further insights into the metabolic function of CLN8, we determined the levels of GPG and various lipids in the whole cell and in purified lysosomes. Lysosomes were purified from WT and *CLN8*-KO Hek293 cells stably expressing the 3xHA-tagged lysosomal membrane protein TMEM192^24^ (**Extended data fig. 3a**), and the enrichment of the lysosomal marker lysosomal associated membrane protein 1 (LAMP1) was confirmed by Western blotting (**Fig. 3a**).

**Figure 3.**
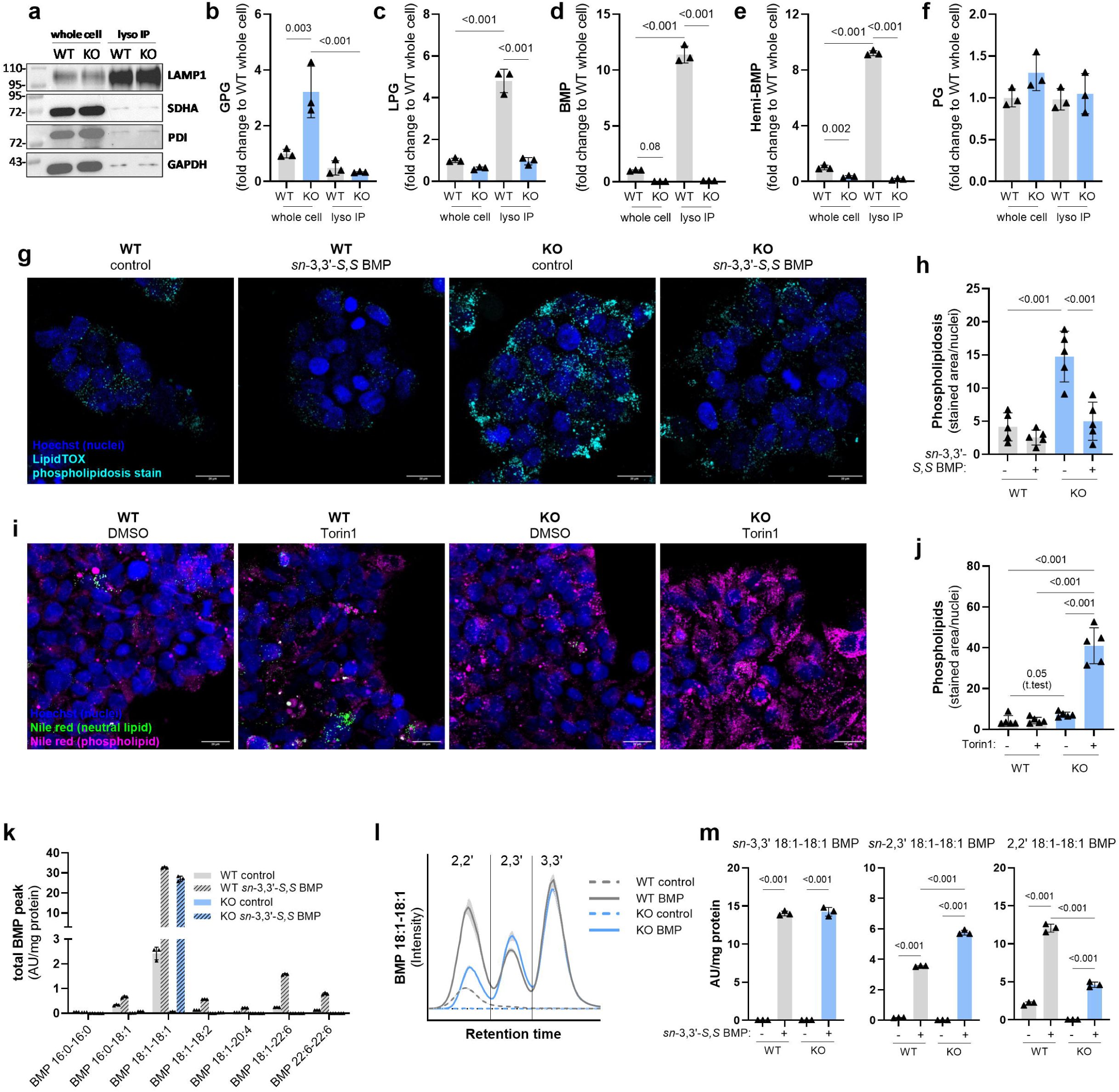
*CLN8*-deletion leads to BMP-deficiency and phospholipidosis. **a,** Expression of the lysosomal marker protein LAMP1 in whole cell fractions and purified lysosomes of WT and *CLN8*-KO Hek293 cells. Lysosomes were isolated via immunopurification^24^ and the purity of lysosomal fractions was confirmed by Western blotting analysis using succinate-dehydrogenase (SDHA), protein disulfide-isomerase (PDI), and glycerinaldehyde-3-phosphate-dehydrogenase (GAPDH) as marker proteins for mitochondria, ER, and cytosol, respectively. Equal volumes of each fraction were used for Western blotting. **b-f,** GPG, LPG, BMP, hemi-BMP and PG levels in whole cell fractions and purified lysosomes of WT and *CLN8*-KO Hek293 cells. Lipid levels in figures c-f are shown as sum of the analyzed subspecies (n=3; whole cell data are representative of two independent experiments). **g,** Fluorescent imaging of formaldehyde-fixed WT and *CLN8*-KO Hek293 cells stained with the HCS LipidTOX™ Green phospholipidosis detection reagent (cyan) and Hoechst (blue). Cells were cultured in the absence and presence of 5 µM *sn*-3,3’-18:1-18:1-*S,S* BMP (*sn*-3,3’*S,S* BMP) for 72h. **h,** Quantification of the area stained by HCS LipidTOX™ phospholipidosis detection reagent (n=5 scans per condition; data are representative of three independent experiments). **i,** Fluorescent imaging of phospholipid deposits (magenta) and neutral lipids (green) in WT and *CLN8*-KO Hek293 cells cultured in the absence and presence of 250 nM Torin1 for 16h. DMSO was used as vehicle control. Cells were fixed and stained with Nile red and Hoechst prior to imaging (n=5 scans per condition; data are representative of two independent experiments). **j,** Quantification of the stained area corresponding to the Nile red phospholipid signal. Scale bar in all images: 20 µm. **k,** BMP subspecies of WT and *CLN8*-KO Hek293 cells cultured in the absence and presence of 5 µM *sn*-3,3’-*S,S* BMP for 96h (n=3). **l,** Representative chromatogram of 18:1-18:1 BMP from experiment i, showing partially separated peaks corresponding to 2,2’, 2,3’ and 3,3’ acyl-chain positions. **m,** Quantification of 2,2’, 2,3’ and 3,3’ 18:1-18:1 BMP peak from experiment i. Data are shown as mean ± SD. Statistical comparisons in all graphs were performed with one-way ANOVA followed by Bonferroni posthoc analysis. *P*-values are indicated.

Whole cell fractions of *CLN8*-KO cells showed a three-fold increase in GPG content compared to WT controls. Independent of the genotype, GPG concentrations were lower in lysosomes than in the whole cell fractions, suggesting that GPG is primarily located outside of acidic organelles (**Fig. 3b**). LPG was enriched five-fold in lysosomes of WT cells compared to whole cells, and decreased by 80% in lysosomal fractions of *CLN8*-KO cells (**Fig. 3c**). This was mostly due to a reduction in the major LPG subspecies 18:1, while specifically LPG 18:0 was increased (**Extended data fig. 3b,e**). As expected from previous observations, BMP was enriched in the lysosomal fraction of WT cells and absent in *CLN8*-KO cells^22^ (**Fig. 3d**). Similarly, hemi-BMP was elevated in lysosomes of WT cells and strongly reduced in *CLN8*-KO cells (**Fig. 3e**). Conversely, PG levels in cells remained unchanged in both fractions irrespective of the genotype (**Fig. 3f**). Changes in subspecies of different lipid classes are shown in **Extended data fig. 3b-i.**

Depleted BMP stores, reduced LPG levels, and unchanged PG levels were also detected in Hap1 cells lacking CLN8 (hemi-BMP was below the detection limit, **Extended data fig. 4a**). In contrast to the Hek293 cell line, Hap1 cells contained substantial amounts of BMP esterified with docosahexaenoic acid (DHA, 22:6), the major species detected in testis and brain^25^ (**Extended data fig. 4b**). It is noteworthy that these cells also contain the presumed precursor LPG 22:6, which was completely absent in *CLN8*-KO cells, while LPG 18:0 was again elevated (**Extended data fig. 4c**).

### *CLN8*-deletion leads to phospholipidosis

BMP promotes lysosomal membrane degradation and lipid export^4^. Therefore, BMP-deficiency is expected to compromise membrane catabolism, leading to phospholipidosis. Using the LipidTOX^TM^ phospholipidosis detection reagent, we observed increased staining of *CLN8*-KO cells, indicating impaired degradation and excessive accumulation of membranes (**Fig. 3g,h**). In line with this observation, LysoTracker^TM^ imaging in live cells revealed increased staining in *CLN8*-KO Hek293 and Hap1 cells (**Extended data fig. 3j,k and 4d,e**). Importantly, the supplementation of cell culture media with commercially available BMP, containing 18:1 FA residues in *sn*-3/*sn*-3’ position (*sn*-3,3’*-S,S*-BMP), strongly reduced phospholipidosis in *CLN8*-KO cells, suggesting that BMP-deficiency is causal for the observed lipid storage disorder (**Fig 3g,h**).

Nile red staining, which can discriminate between polar and neutral lipids^26,27^, further confirmed that *CLN8*-KO cells accumulate phospholipids in lysosomes (**Fig. 3i,j**). The phenotype of these cells was exacerbated in the presence of the mTOR complex inhibitor Torin1, which is known to increase the lysosomal turnover of phospholipids^28^ (**Fig. 3i,j**). Co-staining with LysoSensor^TM^ revealed that Nile red stained phospholipid deposits colocalize with acidic compartments (**Extended data fig. 3l,m**).

The glycerol moieties of endogenous BMP are acylated primarily at the 2,2’ position^29–31^, whereas BMP used in rescue experiments exhibited 3,3’ acyl-configuration. Recent studies have shown that the 2,2’ acyl-configuration of *S,S*-BMP determines its resistance to lysosomal hydrolysis^2^. To investigate cellular FA remodeling, we monitored the FA composition and positional isomers in BMP-supplemented cells. The addition of BMP strongly increased cellular 18:1-18:1 BMP levels in both WT and *CLN8*-KO cells (**Fig. 3k**). BMP subspecies containing other FAs than 18:1 (18:0, 18:2, and 22:6) increased only in WT cells (**Fig. 3k**). Analysis of the positional isomers revealed that the accumulation of 3,3’ BMP was similar in both genotypes. In comparison to WT controls, 2,3′-BMP levels were increased 1.3-fold in *CLN8*-KO cells, while 2,2′-BMP levels were reduced by 60% (**Fig. 3l,m**). These observations suggest that CLN8 affects FA composition and positional remodeling of BMP. Nevertheless, supplementation of *sn*-3,3’*-S,S*-BMP was sufficient to rescue the phospholipidosis phenotype.

Previous studies showed that LPG and PG supplementation promotes BMP formation^10,18,32^. To test whether LPG counteracts BMP-deficiency in *CLN8*-KO cells, we incubated WT and *CLN8*-KO cells with the commercially available *R*-acylated lipid (*sn-*1-oleoyl-*R,rac* LPG). As expected, BMP levels increased 10-fold in WT cells upon LPG supplementation (**Extended data fig. 5a**). Analysis of the positional isomers revealed that 65% and 32% of the generated BMP exhibited 2,2′- and 2,3’ acyl-configuration, respectively (**Extended data fig. 5b**). *CLN8*-KO cells showed a 90% reduction in LPG-induced BMP synthesis. However, we observed a partial rescue by LPG addition reaching basal WT levels (**Extended data fig. 5a**). The BMP formed in *CLN8*-KO cells exhibited exclusively 2,3’ acyl-configuration, suggesting that mature 2,2’-BMP cannot be formed from *R*-acyl LPG in *CLN8*-deficient cells (**Extended data fig. 5b**). Notably, LPG supplementation resulted in ∼80-fold increased cellular GPG content of WT and *CLN8*-KO cells, respectively (**Extended data fig. 5c**), suggesting that the synthesis of 2,2’-BMP in WT cells is driven by the CLN8-dependent utilization of excess GPG.

### CLN8 acylates GPG *in vitro* and *in vivo*

To track the incorporation of GPG into lipids, we loaded cells with [^13^C]GPG^+3^ and analyzed the formation of isotope-labeled LPG and BMP. In contrast to WT controls, *CLN8*-KO Hek293 and Hap1 cells were unable to produce [^13^C]BMP^+3^ (**Fig. 4a, Extended data fig. 6a**) and showed a ∼ 95% reduction in [^13^C]LPG^+3^ formation (**Fig. 4b, Extended data fig. 6b**). Thus, CLN8 is rate-limiting for the incorporation of GPG into phospholipids.

**Figure 4.**
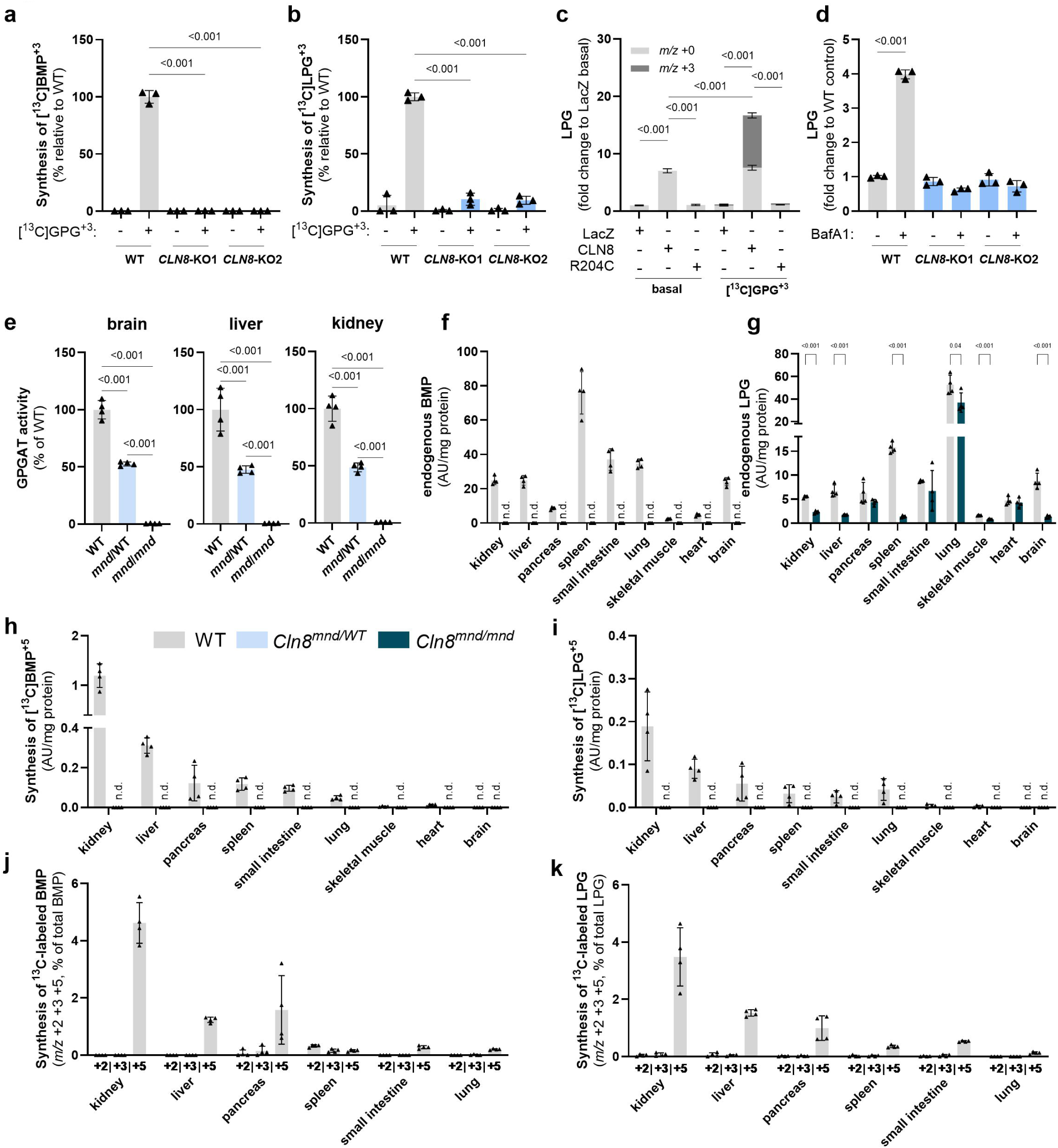
CLN8 enables the utilization of GPG for LPG and BMP synthesis *in vitro* and *in vivo*. **a,b,** Synthesis of [^13^C]BMP^+3^ (a) and [^13^C]LPG^+3^ (b) in WT and *CLN8*-KO Hek293 cells in the presence of 250 µM[^13^C]GPG^+3^. Incubations were performed for 18h under standard cell culture conditions (n=3). **c,** LPG levels in LacZ-, CLN8-, or R204C-overexpressing Hek293 cells under basal conditions and in the presence of exogenously added [^13^C]GPG^+3^ (250 µM; n=3, labeled and unlabeled LPG species are shown). **d,** Effect of 18h Bafilomycin A1 treatment (0.1 µM) on LPG levels of WT and *CLN8*-KO Hek293 cells. DMSO was used as vehicle control (n=3, data are representative of two independent experiments). **e,** GPGAT activity detected in brain, liver, and kidney lysates (1,000 x *g* supernatant) of WT, *Cln8^mnd/^*^WT^, and *Cln8^mnd/mnd^* mice (n=4 mice per genotype). **f,g,** BMP (f) and LPG (g) levels in tissues of WT and *Cln8^mnd/mnd^* mice (n=4 mice per genotype). **h,i,** Levels of [^13^C]BMP^+5^ (h) and [^13^C]LPG^+5^ (i) in tissues of WT and *Cln8^mnd/mnd^* mice 2h post intraperitoneal injection of 1.1 mg [^13^C]GPG^+5^ per animal (55 mg/kg, n=4 mice per genotype). Data classified as n.d. (not detectable) in f,h,i were below the limit of detection, defined as a signal-to-noise ratio of less than 3. **j,k,** Levels of [^13^C]BMP (j) and [^13^C]LPG (k) with *m/z* shifts of +2, +3, or +5, in tissues of WT mice 2h post intraperitoneal injection of 1.1 mg [^13^C]GPG^+5^ (h,i). The results are presented as a percentage of labeled lipids relative to total lipids examined (n=4 mice). Lipid levels are shown as sum of the analyzed subspecies. Data are shown as mean ± SD. Statistical comparison in a-d was performed with one-way ANOVA followed by Bonferroni posthoc analysis and in g with unpaired two-tailed Student’s *t* test. *P*-values are indicated.

Overexpression of CLN8 in Hek293 cells increased cellular LPG levels ∼6- and 16-fold in the absence and presence of exogenously added [^13^C]GPG^+3^, respectively, suggesting that the enzyme acylates GPG from endogenous and exogenous sources. Conversely, expression of the R204C variant had no effect on LPG levels (**Fig. 4c, Extended data fig. 1f**). Re-expression of CLN8 in *CLN8*-KO cells rescued BMP-deficiency and led to a four-fold accumulation of LPG compared to LacZ-expressing cells. Again, no effect was observed in R204C-expressing cells (**Extended data fig. 6c,d**). We next tested whether CLN8 is responsible for the BafA1-induced increase in cellular LPG levels (**Fig. 1p**). *CLN8*-deficiency completely prevented LPG accumulation in BafA1-treated Hek293 and Hap1 cells, thus confirming that CLN8 mediates extra-lysosomal acylation of GPG (**Fig. 4d, Extended data fig. 6e**).

To obtain insights into the *in vivo* function of CLN8, we measured GPGAT activity and BMP synthesis in the motoneuron degeneration mouse (*Cln8^mnd^*) model. *Cln8^mnd^* mice, a model for Batten disease, carry a loss-of-function frameshift mutation in the *Cln8* gene^14,33^.

GPGAT activity in brain, liver, and kidney of WT and mutant mouse models correlated with the *Cln8* gene dosage, showing an ∼50% reduction and no detectable activity in tissue lysates of heterozygous and homozygous mice (*Cln8^mnd/^*^WT^ and *Cln8^mnd^*^/*mnd*^), respectively, compared to WT controls (**Fig. 4e**). Lipidomic analysis revealed that BMP was below the detection limit in all analyzed tissues of *Cln8^mnd^*^/*mnd*^ mice (**Fig. 4f**), while WT controls and *Cln8^mnd/^*^WT^ mice exhibited similar BMP content (**Extended data fig. 7a**). Additionally, we observed strongly decreased LPG levels in brain, kidney, liver, skeletal muscle, and spleen of *Cln8^mnd^*^/*mnd*^ mice (**Fig. 4g**). All analyzed BMP and LPG subspecies of WT, *Cln8^mnd^*^/WT,^ *and Cln8^mnd^*^/*mnd*^ are listed in **Extended data fig. 7a,b**.

To investigate the utilization of GPG for BMP synthesis, we monitored the incorporation of the [^13^C]GPG^+5^ tracer into tissue BMP and LPG. Two hours post intraperitoneal injection, the incorporation of [^13^C]GPG^+5^ into BMP was highest in kidney and liver of WT mice, followed by pancreas, spleen, small intestine, and lung (**Fig. 4j**). Conversely, the formation of [^13^C]BMP^+5^ was not detectable in *Cln8^mnd^*^/*mnd*^ mice. Irrespective of the genotype, no substantial incorporation into BMP was observed in skeletal muscle, heart, and brain of WT mice, suggesting that the tracer is not taken up by myocytes and does not cross the blood-brain-barrier. Formation of [^13^C]LPG^+5^ was detected in kidney, liver, pancreas, spleen, small intestine, and lung of WT, but not *Cln8^mnd^*^/*mnd*^ mice (**Fig. 4i**). Notably, in all WT tissues except for the spleen, [^13^C]BMP were detected with *m/z* +5, but not with *m/z* +3 or +2 (**Fig. 4j**). Similarly, [^13^C]LPG appeared exclusively with *m/z* +5 (**Fig. 4k**), suggesting that GPG is directly acylated for the synthesis of these lipids.

## Discussion

*CLN8* is one of 13 genes that have been associated with the inherited neurodegenerative disorder Batten disease^34^. The genetically distinct forms are grouped together by similar clinical features and the accumulation of autofluorescent storage material in lysosomes of neuronal cells^35^. The age of onset, severity of symptoms, and progression of disease vary between the different types, but most commonly begin during childhood. Typical clinical symptoms include progressive impairment of vision, movement and cognition. Therapy development for Batten disease has been limited because the functions of CLN proteins are only partially understood. Enzyme replacement, gene therapy, and stem cell transplantation are considered the most promising options^34^.

Recent studies provided new insights into the pathogenic mechanisms of Batten disease, revealing that cells lacking CLN3, CLN5, CLN8, and CLN11 develop BMP-deficiency^9,22,36–38^. BMP acts as a cofactor of lysosomal lipid hydrolases and transport proteins that promote the breakdown and export of lipids^39^. It is therefore reasonable to assume that BMP-deficiency impairs lysosomal lipid catabolism in several genetically distinct forms of Batten disease. Our observations suggest that BMP-deficiency is causal for lysosomal lipid accumulation in *CLN8*-KO cells, as BMP supplementation rescued the phospholipidosis phenotype. Thus, the supplementation of BMP may represent a therapeutic option for *CLN8*-deficiency and other lysosomal storage disorders characterized by BMP-deficiency.

Previous work demonstrated that CLN8 is an ER-associated transmembrane protein that regulates lysosomal biogenesis by promoting the ER-to-Golgi transfer of lysosomal enzymes produced at the ER^40,41^. Our study suggests that CLN8 produces LPG at the ER, which is selectively delivered to late endosomes/lysosomes and incorporated into lysosomal BMP. The GPGAT activity is essential for BMP synthesis, as *CLN8*-KO cells and mice lack BMP. CLN8-derived LPG is acylated on *S*-linked glycerol moieties and can therefore not be incorporated into PG and cardiolipin, which contain FA residues exclusively at the *R*-position (**Fig. 5a**). The treatment of cells with BafA1 prevented the conversion of CLN8-derived LPG into BMP and led to the accumulation of LPG, suggesting that BMP but not LPG synthesis requires functional lysosomes. BMP formation in acidic organelles can be catalyzed by CLN5^9^. This transacylase utilizes LPG as both FA donor and acceptor, leading to the formation of BMP and GPG. *CLN5*-deficiency compromises BMP synthesis and is associated with a strong increase in lysosomal LPG levels^9^. Conversely, we found that lysosomal LPG levels are decreased in *CLN8*-deficient cells. Together, these observations indicate that CLN5 uses CLN8-derived LPG as substrate for BMP synthesis.

**Figure 5.**
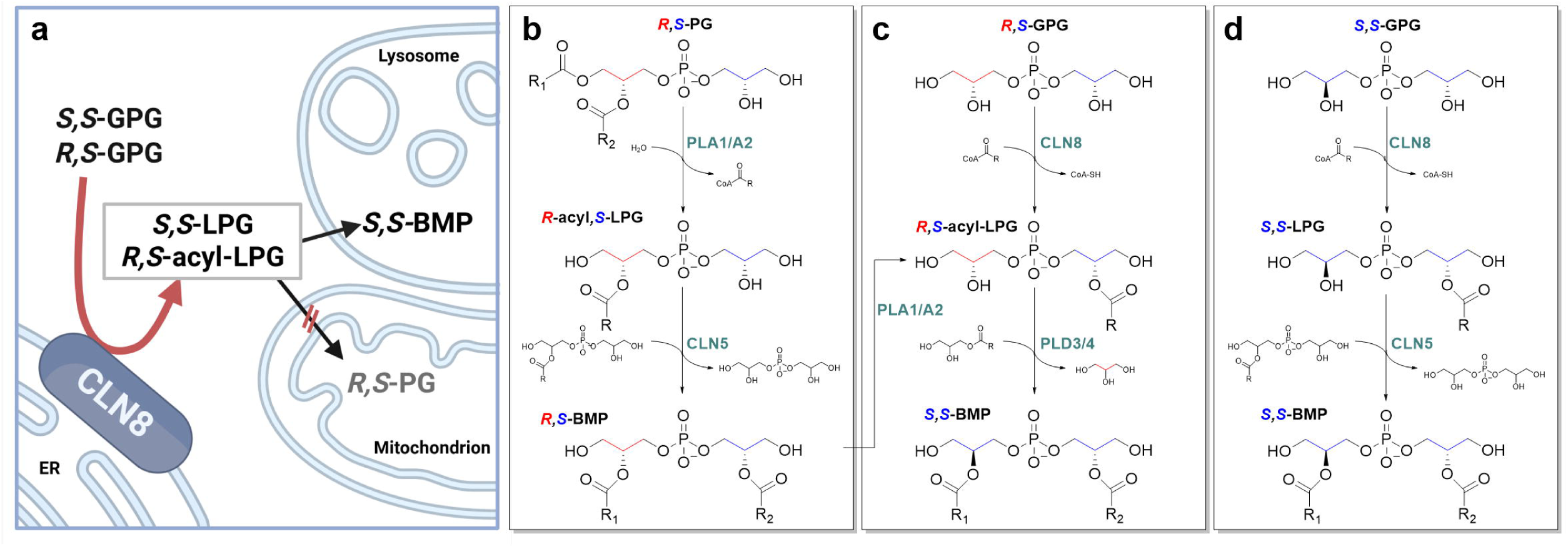
Proposed mechanisms for *S,S*-BMP synthesis. **a,** CLN8-mediated acylation of GPG at *S*-linked glycerol moieties generates *S,S*-^42^, or *R,S*-acyl-LPG at the ER, which enables lysosomal BMP but not PG synthesis (created in *BioRender. Zimmermann, R. (2026)* https://BioRender.com/l57i8gq). **b,** *R,S*-PG is first hydrolyzed by PLA1/A2 enzymes, producing *R*-acyl,*S*-LPG, which can act as acyl acceptor and donor for the CLN5-mediated transacylase reaction, producing *R,S*-BMP, that requires conversion into mature *S,S*-BMP. **c,** The transphosphatidylation reaction, catalyzed by PLD3/4, uses *R,S*-acyl-LPG as phosphatidyl donor and monoacylglycerol as acceptor, generating *S,S*-BMP and glycerol. The precursor for this reaction, *R,S*-acyl-LPG, can either be produced by CLN8-mediated acylation of *R,S*-GPG at the *S*-bound glycerol unit or by PLA1/A2-mediated hydrolysis of *R,S*-BMP at the *R*-bound glycerol. **d,** The acylation of *S,S*-GPG by CLN8 results in *S,S*-LPG, which can be converted into *S,S*-BMP by CLN5^42^.

The transacylation of PG-derived *R,S*-LPG by CLN5 results in BMP with *R,S*-configuration (**Fig. 5b**), which must be converted into mature *S,S*-BMP. A recent study suggests that this stereoconversion can be catalyzed by lysosomal phospholipases D3/D4 (PLD3/4) via transphosphatidylation reactions^10^. The authors proposed that PLD3/4 use *R,S*-LPG, acylated on the *S*-linked glycerol moiety (*R,S-*acyl-LPG), as phosphatidyl donor and monoacylglycerol as acceptor, leading to the formation of *S,S*-BMP and to the release of the *R*-linked glycerol (**Fig. 5c**). The required *R,S-*acyl-LPG precursor can be delivered either by the hydrolysis of *R,S*-BMP by acid phospholipase A1/2 (PLA2G15^2^) or by CLN8 (**Fig. 5c**). Our experiments revealed that the glycerol moieties of GPG are not exchanged during BMP synthesis *in vitro*. Experiments in mice suggested that the GPG tracer is used for BMP synthesis in peripheral tissues, but does not cross the blood-brain barrier. In all tissues except the spleen, GPG was incorporated into BMP without exchange of the glycerol backbones. These observations argue against a PLD3/4-catalyzed stereoconversion in most peripheral tissues. However, it must be considered that experiments were performed with a racemic GPG mixture containing *S,S-*GPG, which can be utilized for the synthesis of *S,S-*BMP without the need for stereoconversion (**Fig. 5d**). In an independent study, Sheokand *et. al.*^42^ identified CLN8 as a stereospecific GPG acyltransferase, which preferentially mediates the acylation of *S,S-*GPG. The generated *S,S-*LPG can be converted into *S,S*-BMP by CLN5.

In summary, our data suggest that CLN8 catalyzes the stereospecific acylation of the *S*-linked glycerol of GPG at the ER, which directs LPG into BMP synthesis and prevents its incorporation into other lipids. *CLN8*-deficiency results in depleted BMP stores *in vitro* and *in vivo*, which compromises lysosomal membrane catabolism and leads to phospholipidosis. We conclude that GPG acylation and BMP synthesis are interrelated metabolic processes that are highly relevant for lysosomal lipid homeostasis and the development of neurodegeneration in patients with CLN8-deficiency. To our knowledge, this is the first report showing that a GPD is utilized for *de novo* lipid synthesis in mammalian cells, although GPD acylation provides a direct synthesis route that is less energy-intensive than *de novo* synthesis and allows FA remodeling of membrane lipids in *sn-1* and *sn-2* positions. Related metabolic lipid synthesis pathways have been described in yeast and plants that recycle GPC for PC synthesis^43,44^.

---

**Extended data figure 1.**
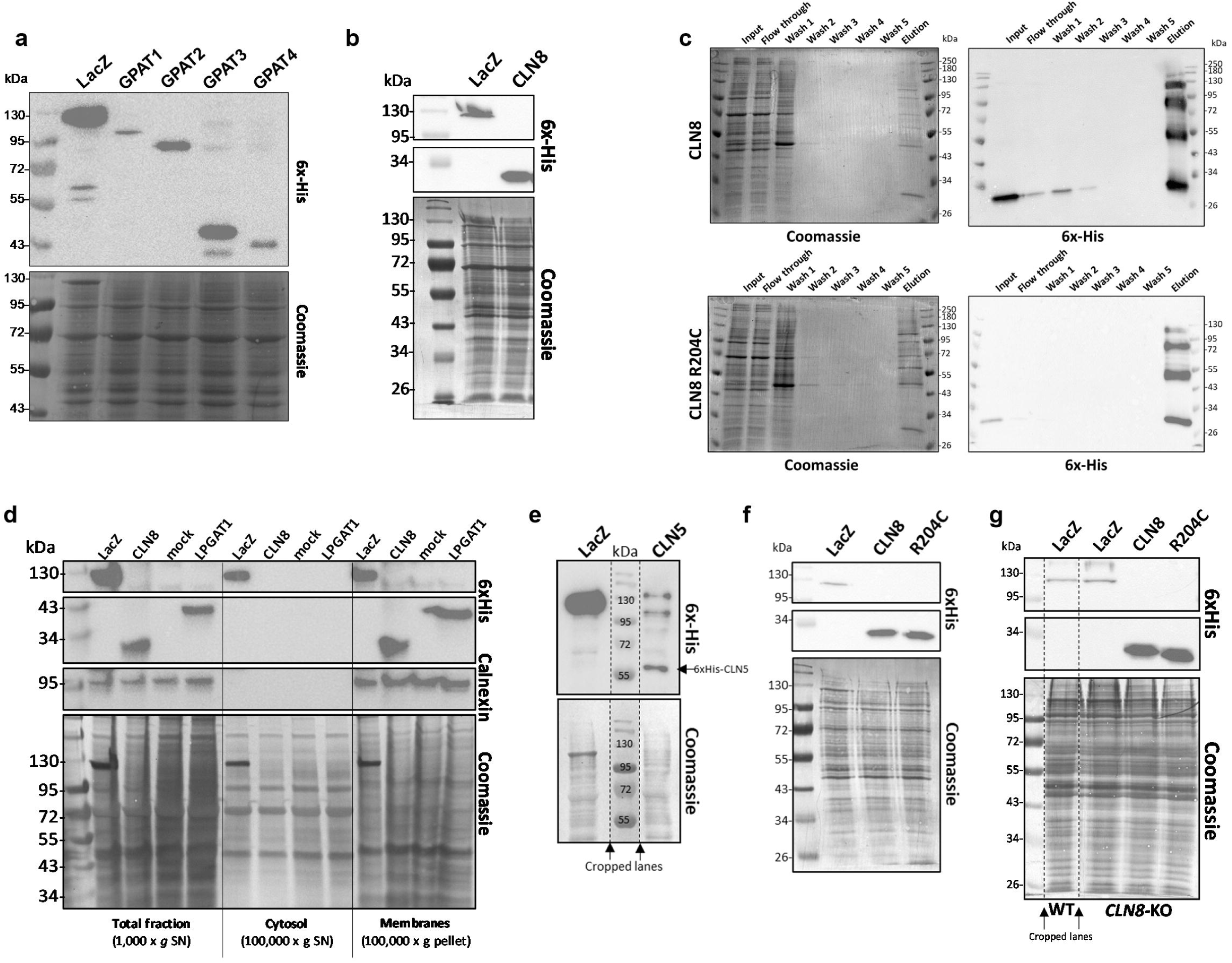
Expression of recombinant proteins and purification of CLN8. **a,b,** Western blot confirming the overexpression of 6xHis-tagged LacZ, GPAT1-4 (a), and CLN8 (b) in Expi293 cells. **c,** Coomassie stained gel and Western blot of fractions obtained during the 6x-His affinity purification of CLN8 and the CLN8 R204C mutant. In line with published data, multiple bands in the purified fraction (elution) represent multimeric aggregates of CLN8^22^. **d,** Western blot of total fraction (1,000 x *g*), cytosolic fraction (100,000 x *g* SN), and membrane fraction (100,000 x *g* pellet) of Expi293 cells overexpressing 6xHis-tagged LacZ, CLN8, LPGAT1, or mock control (transfection of empty vector). Calnexin was used as a marker for membranes (ER). **e,** Western blot confirming the overexpression of 6xHis-tagged LacZ, or CLN5 in Expi293 cells. **f,** Western blot confirming the overexpression of 6xHis-tagged LacZ, CLN8, and CLN8 R204C in Hek293 cells. **g,** Western blot confirming the overexpression of 6xHis-tagged LacZ, CLN8, and R204C in WT and *CLN8*-KO Hek293 cells. In all blots, Coomassie stain was used as loading control. Dashed lines indicate cropped lanes.

**Extended data figure 2.**
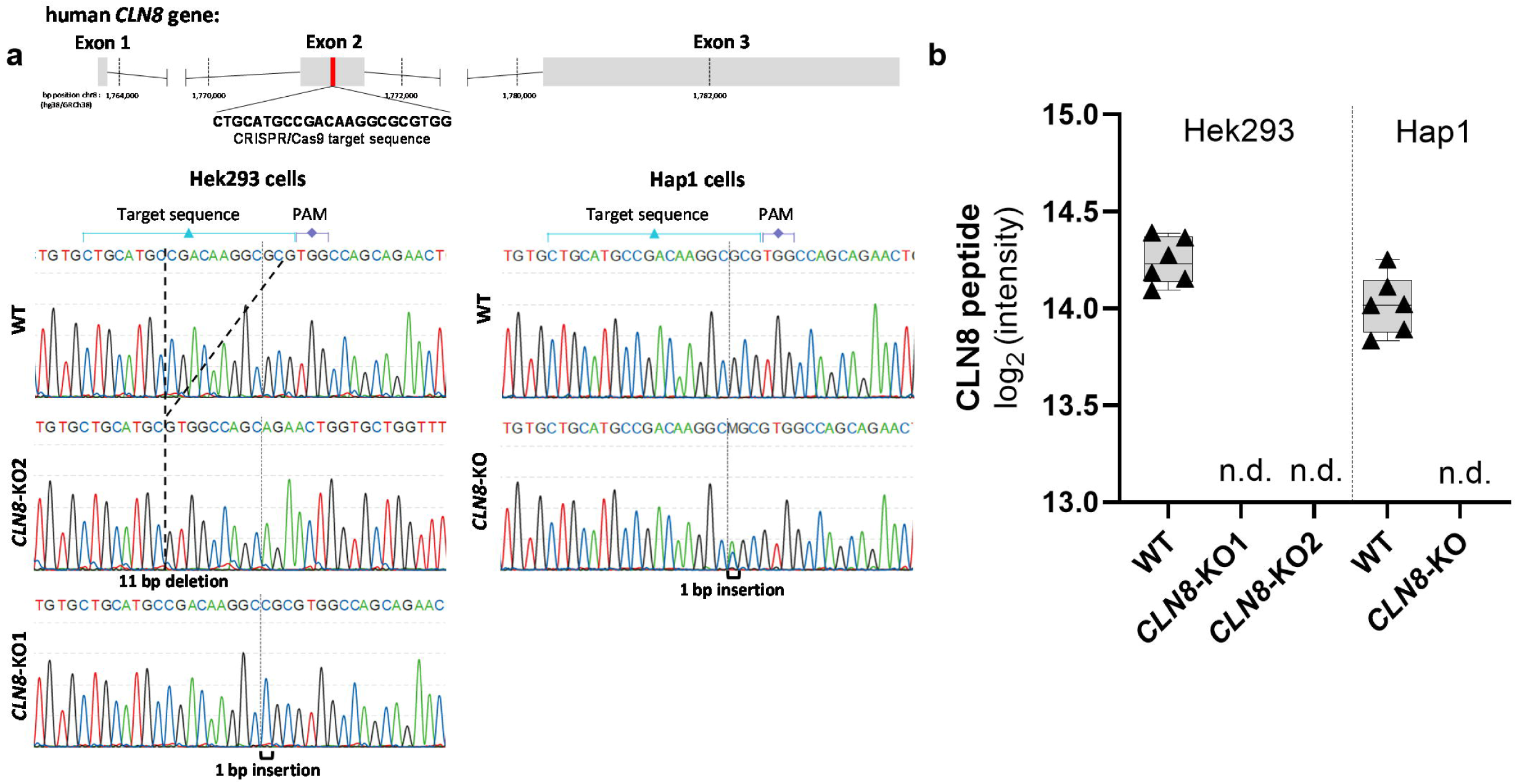
Confirmation of *CLN8*-KO in Hek293 and Hap1 cells. **a,** Sanger sequencing of genomic DNA at the CRISPR-Cas9 targeting locus in the *CLN8* gene (Exon2) of Hek293 and Hap1 WT and *CLN8*-KO clones. CRISPR-Cas9 mediated deletions or insertions are highlighted in the sequencing chromatogram. **b,** Mass Spectrometry based proteome analysis of total Hek293 and Hap1 cell extracts. Relative CLN8 protein levels (log_2_ intensity) of WT and *CLN8*-KO lines are shown (n=6). No CLN8 peptide was detected in KO clones (n.d.).

**Extended data figure 3.**
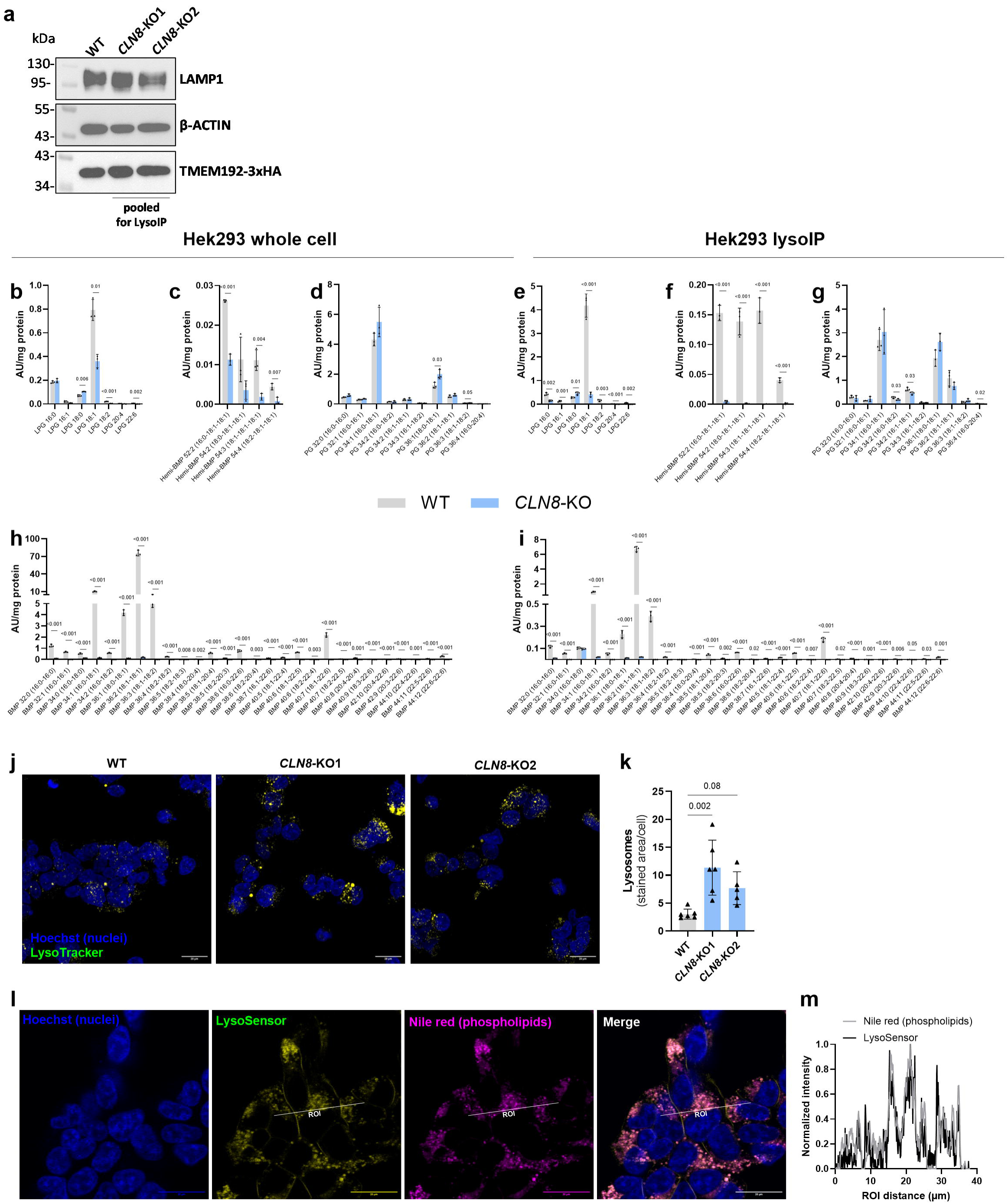
*CLN8*-deficiency disrupts BMP synthesis in Hek293 cells. **a,** Western blot confirming the lentiviral expression of TMEM192-3xHA in Hek293 WT and *CLN8*-KO clones. *CLN8*-KO clones 1 and 2 were pooled for lysosome immunopurification. **b-i,** LPG, BMP, hemi-BMP and PG subspecies in whole cell fractions (b,c,d,h) and purified lysosomes (e,f,g,i) of WT and *CLN8*-KO Hek293 cells (n=3). **i,** Live cell imaging of WT and *CLN8*-KO Hek293 cells. Lysosomes were stained with LysoTracker^TM^ and nuclei were stained with Hoechst. **k,** Quantification of LysoTracker^TM^ signal (n=6 scans per condition). **l,** Representative live cell imaging of *CLN8*-KO Hek293 cells treated with 250 nM Torin1 for 16h. Nuclei, lysosomes, and phospholipids were stained with Hoechst, LysoSensor^TM^, and Nile red respectively. **m,** Normalized phospholipid (Nile red) and LysoSensor^TM^ signal intensity in linear region of interest (ROI), shown as a white line in l. Scale bars in all images 20 µm. Statistical comparisons in b-i were performed with unpaired two-tailed Student’s *t* test and in k with one-way ANOVA followed by Bonferroni posthoc analysis. *P*-values are indicated.

**Extended data figure 4.**
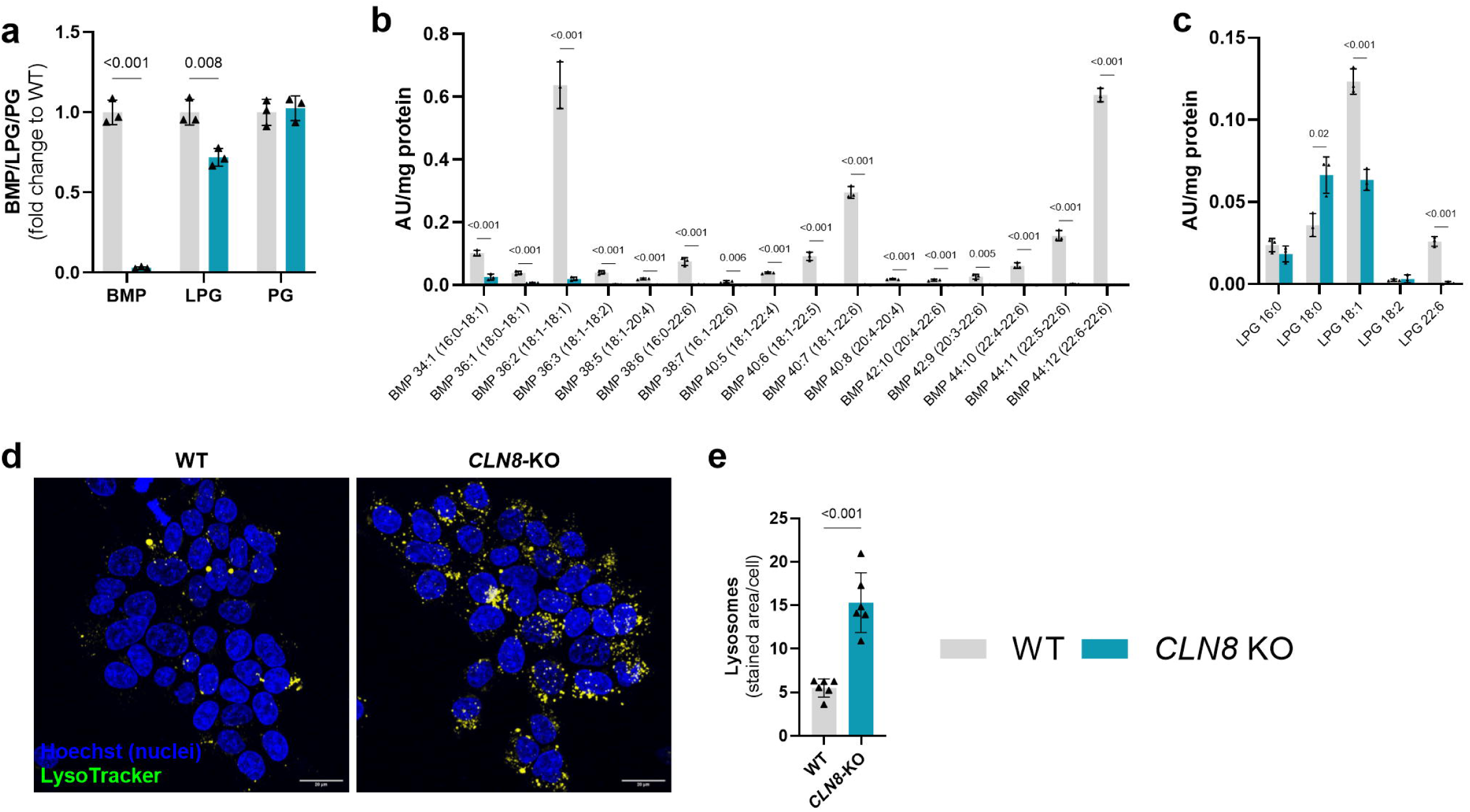
*CLN8*-deficiency disrupts BMP synthesis in Hap1 cells: **a,** BMP, LPG and PG levels in whole cell fractions of WT and *CLN8*-KO Hap1 cells (sum of the analyzed subspecies, n=3). **b,c,** BMP and LPG subspecies in whole cell fractions of WT and *CLN8*-KO Hap1 cells. (n=3; data are representative of 2 independent experiments). **d,** Live cell imaging of WT and *CLN8*-KO Hap1 cells. Lysosomes were stained with LysoTracker^TM^ and nuclei were stained with Hoechst. Scale bars in all images 20 µm. **e,** Quantification of LysoTracker^TM^ signal (n=6 scans per condition). Statistical comparisons were performed with unpaired two-tailed Student’s *t* test. *P*-values are indicated.

**Extended data figure 5.**
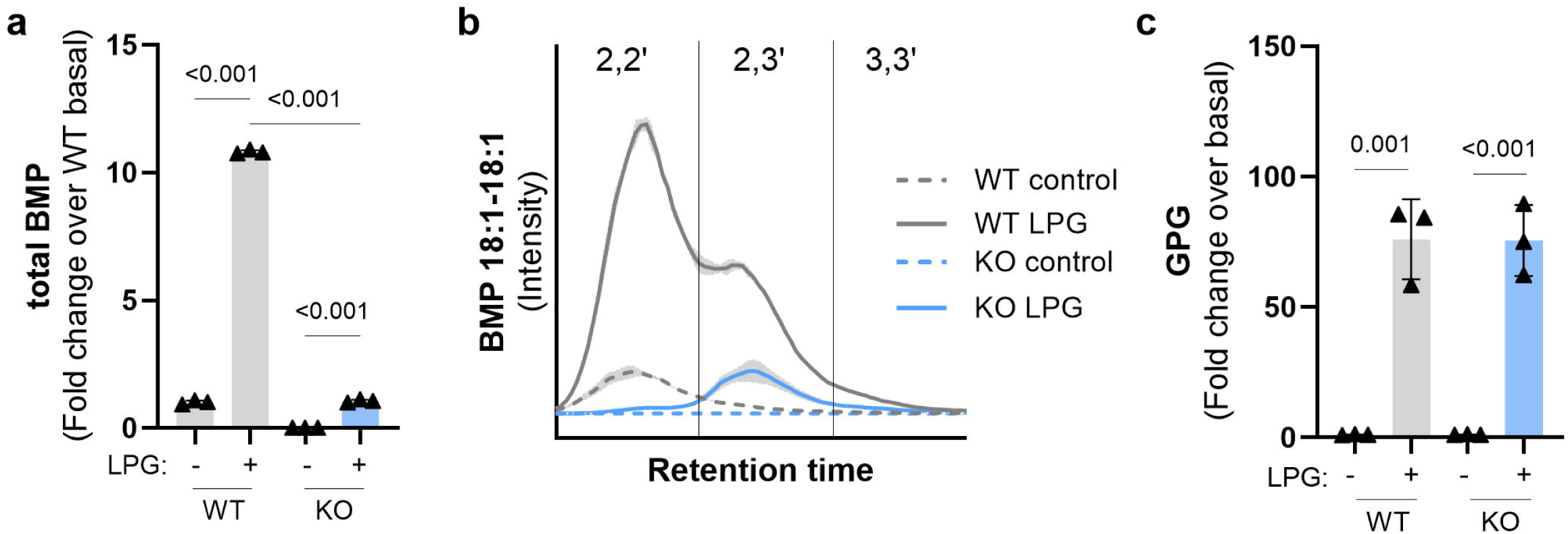
*R*-acyl-LPG supplementation in *CLN8*-deficient cells. **a,** Total BMP levels of WT and *CLN8-*KO Hek293 cultured in the absence and presence of 50 µM *sn-*1-oleoyl LPG (*R,rac*) for 18h (n=3). **b,** Chromatogram of 18:1-18:1 BMP, showing the partially separated peaks corresponding to 2,2’, 2,3’ and 3,3’ acyl-configurations, in control and LPG loaded WT and *CLN8*-KO Hek293 cells shown in a (n=3). **c,** GPG levels of control and LPG loaded WT and *CLN8*-KO Hek293 cells shown in a (n=3). Data are shown as mean ± SD. Statistical comparison was performed with unpaired two-tailed Student’s *t* test. *P*-values are indicated.

**Extended data figure 6.**
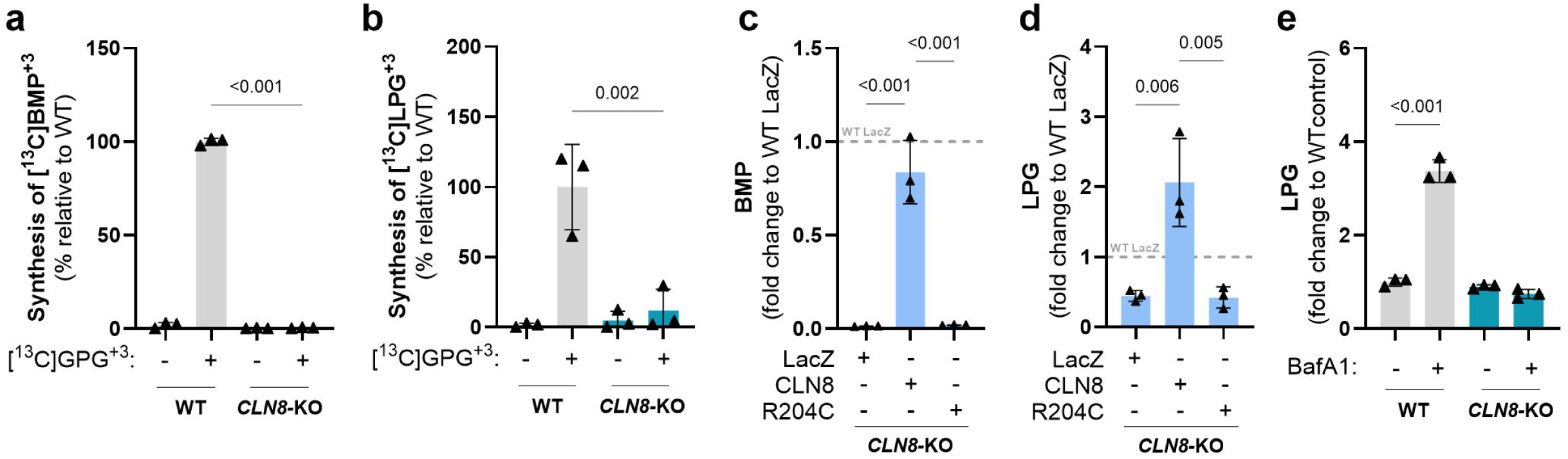
CLN8 enables the utilization of GPG for LPG and BMP synthesis. **a**,**b,** Synthesis of [^13^C]BMP^+3^ (a) and [^13^C]LPG^+3^ (b) in WT and *CLN8*-KO Hap1 cells in the presence of 250 µM [^13^C]GPG^+3^. Incubations were performed for 18h under standard cell culture conditions (n=3). **c,d,** BMP and LPG levels in LacZ-, CLN8-, or R204C-overexpressing *CLN8*-KO Hek293 cells. Data are shown as fold-change to WT Hek293 cells expressing LacZ (dashed line; n=3; data are representative of two independent experiments). **e,** Effect of 18h Bafilomycin A1 treatment (0.1 µM) on LPG levels of WT and *CLN8*-KO Hap1 cells. DMSO was used as vehicle control (n=3, data are representative of two independent experiments). Lipids were analyzed with LC-MS. Data are shown as mean ± SD. Statistical comparison was performed with one-way ANOVA followed by Bonferroni posthoc analysis. *P*-values are indicated

**Extended data figure 7.**
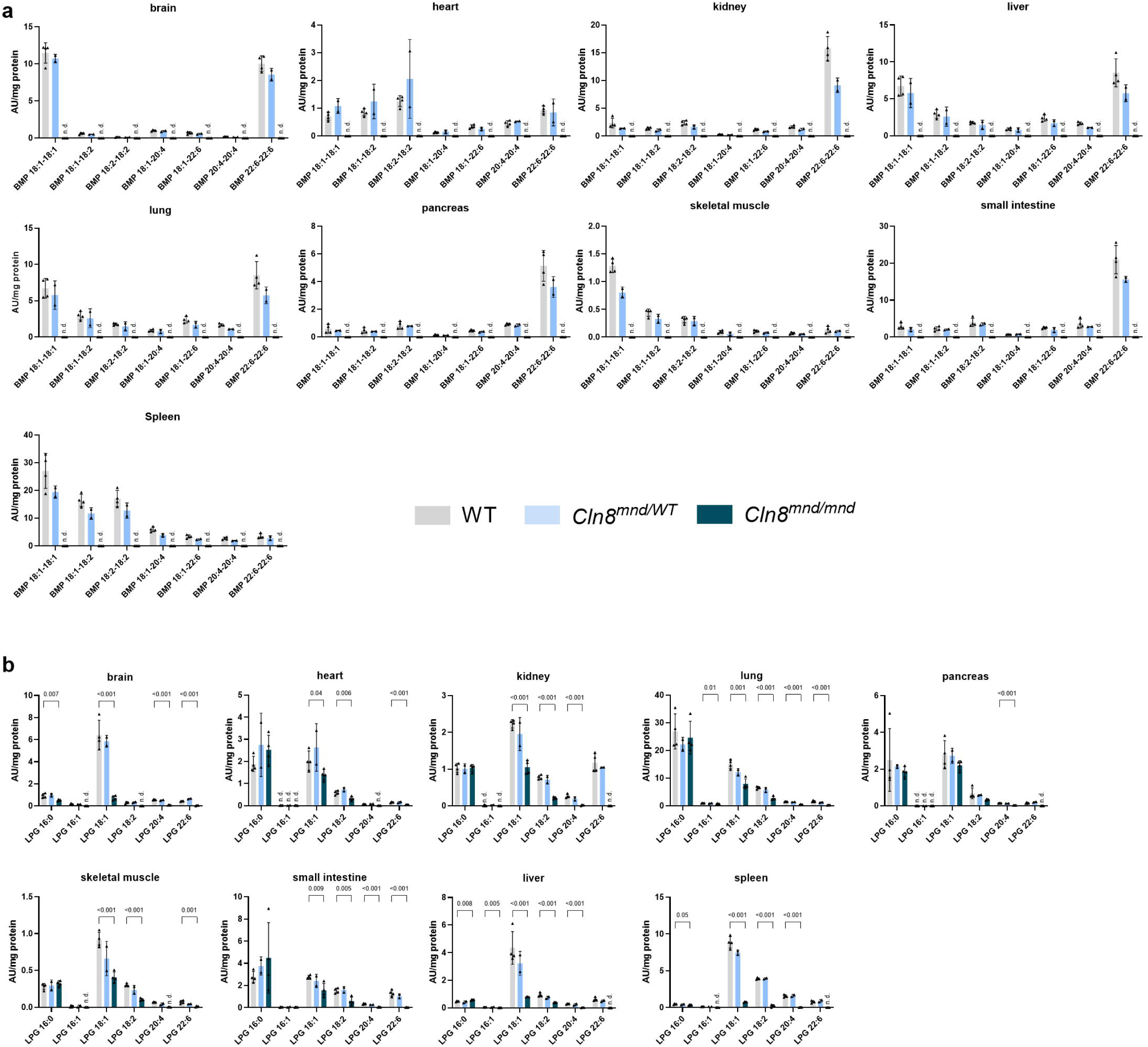
BMP and LPG subspecies in WT, *Cln8^mnd^*^/WT^, and *Cln8^mnd^*^/*mnd*^ mice. **a,** BMP subspecies. **b,** LPG subspecies (n=4 for WT and *Cln8^mnd^*^/*mnd*^, n=2 for *Cln8^mnd/^*^WT^). Data classified as n.d. (not detectable) were below the detection limit, defined as a signal-to-noise ratio of less than 3. Data are shown as mean ± SD. Statistical comparisons between WT and *Cln8^mnd^*^/*mnd*^ were performed with unpaired two-tailed Student’s *t* test. *P*-values are indicated.

## Methods

### Reagents, materials, and antibodies

Detailed information on all reagents, materials, plasmids and antibodies used during this study are listed in **Supplementary table 1-3**.

### Cell culture

For regular maintenance, Hek293 (CRL-1573, ATCC) or Hap1 (C631, Horizon) cells were cultured in DMEM 4.5 g/l glucose or IMDM (both Gibco), respectively, both supplemented with 10% heat-inactivated fetal bovine serum (FBS) as well as 100 IU/ml penicillin and 100 IU/ml streptomycin. For heat-inactivation, FBS was prewarmed to 37°C and subsequently incubated for 1h at 56°C in a water bath.

Expi293F cells (A14527, Gibco) were cultured in Expi293T Expression Medium (A1435101, Gibco) in 25 ml or 10 ml culture volumes using 125 ml vented Erlenmeyer flasks (431143, Corning) or 50 ml CELLSTAR cell reactor tubes (227245, Greiner Bio-One) on an orbital shaker platform at 125 or 220 rpm, respectively. Cell lines were regularly checked for mycoplasma contamination using the Mycoplasmacheck Service from Eurofins Genomics GmbH (Germany). All cells were grown under standard conditions (37°C, 95% humidified atmosphere, 7% CO_2_).

### Animals

Experimental procedures were approved by the University of Cambridge Animal Welfare and Ethics Board and UK Home Office (PPL number PP1740969) and were conducted in accordance with the EU Directive 2010/63. All animal procedures were guided by the 3R’s principles to minimize pain, suffering, and distress. *Cln8^mnd^* mice (AK.B6-*Cln8^mnd^*/J; JAX strain number 003906), were maintained in a SOPF facility on a regular light-dark cycle (12 h light, 12 h dark) at a room temperature (RT 23 ± 1°C) and fed *ad libitum* on a standard laboratory chow diet (SAFE 105; Rosenburg, Germany). Homozygous experimental animals were generated by crossing heterozygous parents. Animals were killed by cervical dislocation. All studies involving animals are reported in accordance with the ARRIVE guidelines.

### Transfection of Expi293, Hek293, and Hap1 cells

To generate cell lysates containing recombinantly expressed GPAT1-4, LacZ, LPGAT1, CLN5, CLN8, or CLN8 R204C for *in vitro* assays or protein purification, Expi293F cells were transfected according to the manufacturer’s instructions and as previously described^45^.

For transient protein expression in Hek293 and Hap1 cells, 6×10^6^ cells were seeded into 100 mm culture dishes and transfected with the Lipofectamine 3000 transfection reagent (L3000001, Invitrogen) the following day. One 100 mm dish was transfected according to the following protocol: 600 μl DMEM without antibiotics and FCS (DMEM-/-) premixed with 30 μl Lipofectamine 3000 were pooled with DMEM-/- containing 15 μg plasmid DNA and 30 μl P3000 reagent. The solution was incubated for 15 min at RT and added dropwise to the medium. For cell culture experiments, the cells were trypsinized 24h post transfection, resuspended in fresh medium, and seeded into 6-well plates at a density of 0.7×10^6^ cells per well. Cells were harvested 48h post transfection. All plasmids used for transfection are listed in **Supplementary table 2**.

### Cell culture experiments

All cell culture experiments were conducted in 6- or 12-well plates with an initial seeding density of 1×10^6^ or 0.5×10^6^ cells per well, respectively.

For isotope tracer experiments, the cells were seeded into 6-well plates and cultured overnight. On the next day, the cells were incubated for 18h with 500 µM [^13^C]GPG^+3^, 250 µM [^13^C]GPG^+5^, 500 µM [U-^13^C]glycerin (489476, Sigma-Aldrich), or DMEM where glucose was replaced by 25 mM [U-^13^C]glucose (389374, Sigma-Aldrich). Cells cultured in standard cultivation medium were used as controls.

For isotope tracer experiments with inhibitors, both Hek293 and Hap1 cells were seeded in 6-well plates. The next day, the cells were preincubated for 1h with 10 µM TriacsinC (T4540, Sigma-Aldrich), 100 nM Bafilomycin A1 (HY-100558, MedChemExpress), or DMSO as vehicle control and further cultured for 18h in the presence of 500 µM [^13^C]GPG^+3^ maintaining constant inhibitor concentrations.

Isotope tracer experiments with *CLN8*-KO Hek293 and Hap1 cells were performed by seeding cells in 12-well plates. The following day cells were cultured in the presence of 250 µM [^13^C]GPG^+3^ for 18h. For BMP supplementation experiments, WT and *CLN8*-KO Hek293 cells were cultured in medium supplemented with and without 5 µM *sn*-3,3’-*S,S*-18:1-18:1 BMP (857135, Avanti Polar Lipids) for 72h with medium change every other day, before being reseeded into 12-well plates and cultured for another 24h in the presence of 5 µM BMP.

LPG supplementation experiments were performed by seeding cells in 12-well plates. The following day cells were cultured in the presence of 50 µM *sn-*1-oleoyl-*R,rac* LPG (858125, Avanti Polar Lipids) for 18h. BMP and LPG were emulsified in media via brief sonication in a water bath.

Before lipidomic analysis, the cells were washed twice with PBS and collected by gentle flushing in PBS. Cellular lipids were extracted as described and endogenous as well as isotope-labeled lipids were analyzed by LC-MS.

### Purification of 6xHis-tagged CLN8 and CLN8 R204C

CLN8 and CLN8 R204C were isolated using 6xHis affinity chromatography. Therefore, half of a cell pellet from a 25 ml culture of Expi293 cells containing recombinantly expressed proteins was resuspended in 6 ml lysis buffer composed of 50 mM HEPES (pH 7.25), 0.1 mM EDTA, 5 mM β-mercaptoethanol, 5% glycerol, 150 mM NaCl, a protease inhibitor cocktail (1 µg/ml pepstatin, 2 µg/ml antipain, 20 µg/ml leupeptin) and 1.5% lauryl maltose neopentyl glycol (LMNG, A50940, Thermo Scientific). The suspension was incubated on ice for 2h, with occasional inversion of the tube to ensure thorough lysis. The lysate was centrifuged at 40,000 x *g* for 30 min at 4°C to remove insoluble material. The resulting supernatant was diluted to 50 ml with lysis buffer lacking LMNG. This solution was mixed with 250 µl of pre-washed (3x in PBS) PureCube 100 Ni-NTA agarose beads (74103, Cube Biotech), and incubated overnight at 4°C on an overhead shaker. The following day, the agarose beads were transferred to a gravity column and washed in five steps. Each washing buffer contained 50 mM HEPES pH 7.25, 0.1 mM EDTA, 5 mM β-mercaptoethanol, 5% glycerol, 1 mM DTT, 0.1% LMNG, protease inhibitor cocktail, imidazole, and NaCl. The concentration of imidazole and NaCl in the washing buffers were as follows: imidazole (mM)/NaCl (mM): Buffer 1: 50/500; Buffer 2: 20/500; Buffer 3: 10/500; Buffer 4: 0/250; Buffer 5: 0/125. The beads were initially washed with washing buffer 1 in 500 µl steps until no more protein was detected in the washing fractions using the Bradford reagent. After that, the beads were washed consecutively with 3 ml of washing buffer 2 to 5. Elution of the purified proteins was performed in 20 x 100 µl steps using an elution buffer containing 50 mM HEPES pH 7.25, 125 mM NaCl, 0.1 mM EDTA, 5 mM β-mercaptoethanol, 5% glycerol, 300 mM imidazole, 1 mM DTT, the protease inhibitor cocktail, and 0.05% LMNG. All protein-containing fractions were pooled, analyzed for protein concentration using the Bradford reagent, aliquoted, snap-frozen in liquid N_2_, and stored at - 80°C. The purification steps and the purity of the isolated proteins (10 µl of each fraction) were analyzed by SDS-PAGE followed by Coomassie staining and Western blotting.

### CRISPR/Cas9-mediated deletion of *CLN8*

*CLN8*-deficient Hek293 and Hap1 cells were generated by CRISPR/Cas9 using the pSpCas9(BB)-2A-Puro (PX459) V2.0 vector (gift from Feng Zhang, Addgene, plasmid #62988) and 5’-CTGCATGCCGACAAGGCGCGTGG-3’ as sgRNA, which was designed using chopchop^46^. The sgRNA was cloned into the *BbsI* site of pSPCas9(BB)-2A-Puro V2.0. Correct insertion was verified by Sanger sequencing. Hek293 and Hap1 cells were transfected with the resulting construct using the Lipofectamine 3000 Reagent (L3000001, Invitrogen) as described above. Cells transfected with empty pSPCas9(BB)-2A-Puro V2.0 served as control. Twenty-four hours after transfection, 2 µg/ml puromycin were added for the selection of successfully transfected cells. After clonal expansion for two-three weeks, the cell lines were screened for mutations in the *CLN8* gene by sequencing a PCR product containing the sgRNA target site (forward primer 5’-GTCGCTGGCTTTGTCTTCTACT-3’; reverse primer 5’-CTTTAAGAGCATCCAGGAAACG-3’). Control cells transfected with an empty vector were subjected to the same procedure, including passaging and validation of the genomic sequence by Sanger sequencing.

### Chemical synthesis of [^13^C]GPG^+3^, [^13^C]GPG^+5^ and generation of [^13^C]GPG^+3/PG^ from headgroup labeled PG

A detailed procedure on the chemical synthesis of [^13^C]GPG^+3^ and [^13^C]GPG^+5^ is provided as **Supplementary Methods.**

[^13^C]GPG^+3/PG^ was generated via saponification of headgroup-labeled di-oleoyl [^13^C]PG. Di-oleoyl [^13^C]PG was synthesized by the conversion of di-oleoyl PC into di-oleoyl PG by cabbage phospholipase D (PLD) (P8398, Sigma Aldrich) in the presence of [U-^13^C]glycerin as described previously^18^.

For saponification, 1 mg of di-oleoyl [^13^C]PG was dissolved in 60 µl of methyl *tert*-butyl ether/methanol (3/1, v/v) and mixed with 30 µl 2 M NaOH for 2h at RT. The solution was neutralized by the addition of 30 µl 2 M HCl. After further adding 340 µl methyl *tert*-butyl ether/methanol (3/1, v/v) and 60 µl ddH_2_O, the solution was vortexed and centrifuged at 20,000 rpm for 5 min. The upper organic phase was quantitatively removed and the aqueous phase containing, GPG was dried at 45°C under a stream of N_2_. The dried extract, including NaCl generated during saponification, was reconstituted in PBS, and the concentration of GPG was determined by comparing the LC-MS signal against the chemically synthesized [^13^C]GPG^+5^.

### Microscopy

All microscopy glass slides used for culturing cells, were precoated with poly-D-lysine (A-003-M, Sigma-Aldrich).

For LysoTracker Red DND-99 (L7528, Invitrogen) staining in live cells, 350,000 WT and *CLN8*-KO Hek293 and Hap1 cells were seeded into 35 mm glass bottom dishes (81218-200, Ibidi). The next day, cells were cultured for 1h in medium containing 62 nM Lysotracker and 1 µg/ml Hoechst 3342 (ab228551, Abcam). The experiment was timed between conditions to ensure uniform exposure of cells to the fluorescent dyes.

For phospholipidosis staining, Hek293 WT and *CLN8*-KO cells were cultured with and without 5 µM *sn*-3,3’-*S,S*-18:1-18:1 BMP (857135, Avanti Polar Lipids) in 100 m culture dishes. BMP was emulsified in DMEM medium via brief sonication. After 24h, the cells were reseeded in 8-well chamber slides (94.6170.802, Sarstedt) at a density of 100,000 cells per well. The following day, the cells were incubated with medium containing HCS LipidTOX™ Green phospholipidosis detection reagent (H34350, Invitrogen) for another 24h. Throughout all steps, BMP supplementation was maintained at 5 µM. For imaging, the cells were fixed with 4% PBS buffered formaldehyde and stained with 1 µg/ml Hoechst 3342.

For Nile red staining, Hek293 WT and *CLN8*-KO cells were seeded in 8-well chamber slides at a density of 100,000 cells per well. On the next day, the cells were cultured with DMEM medium containing 250 nM Torin1 (475991, Sigma-Aldrich) or DMSO vehicle for 16h. For imaging, the cells were fixed with 4% PBS-buffered formaldehyde and subsequently stained with 1 µg/ml Nile red (72485, Sigma-Aldrich) and 1 µg/ml Hoechst 3342. Colocalization experiments were performed in live cells by culturing Torin1-treated *CLN8*-KO cells for 1h in medium containing 1 µg/ml Nile red, 1 µg/ml Hoechst 3342 and 1 µM LysoSensor^TM^ Green DND-189 (L7535, Invitrogen).

Imaging data were acquired with a Leica SP8 confocal microscope with spectral detection (Leica Microsystems, Inc.), using an HC PL APO CS 63×/1.2 NA water immersion objective. Scans were performed in sequential mode to avoid fluorescence bleed-through into different channels. Excitation (ex) of fluorescent dyes and recording of their fluorescent emission (em) were conducted at the following wavelengths: Hoechst 3342 ex: 405 nm, em: 420-452 nm; HCS LipidTOX™ Green ex: 488 nm, em: 507-549 nm; LysoTracker™ Red DND-99: ex: 561 nm, em: 574-666 nm; Nile red - phospholipids: ex: 561 em: 620-660 nm; Nile red - neutral lipids ex: 514 nm, em: 534-574 nm. The imaging data were acquired in z-stacks, with the exception of the colocalization studies. For image quantification, the open-source software Fiji was used. The fluorescent channels were split and z-stacks were converted into a maximum intensity projection. Threshold values were set individually for each channel and kept constant for all images throughout all conditions. Based on the threshold, a mask was created before particles were analyzed using the built-in “Analyze particle” function, with a particle size ranging from 0 to infinity structures. Nuclei in each scan were counted manually. To determine the average fluorescent area/per cell, positive areas were summed and divided by the number of nuclei in each image. To visualize colocalization of the Nile red phospholipid- and LysoSensor signals, the individual- and merged fluorescent channels are shown. Additionally, the fluorescent intensities of both signals along a linear region of interested (ROI) were normalized to respective maximum values and plotted against the ROI distance.

### *In vivo* tracer experiments with [^13^C]GPG^+5^

*In vivo* tracer experiments were performed with female WT and *Cln8^mnd^*^/*mnd*^ mice, aged 10 weeks. [^13^C]GPG^+5^ was dissolved in PBS and administered by intraperitoneal injection at a dosage of 1.1 mg [^13^C]GPG^+5^ per animal (55 mg per kg bodyweight). Tissues were collected 2h post injection and snap-frozen in liquid N_2_. 10-20 mg of pulverized tissues were used for lipid analysis. Incorporation of the tracer into phospholipids was determined as described in the mass spectrometry section.

### Western blotting

Protein samples were mixed with SDS-sample buffer (0.05 M Tris–HCl, 0.1 M DTT, SDS 2% w/v, 1.5 mM bromophenol blue, 1.075 M glycerol). Samples used for the detection of CLN8 were not subjected to heat denaturation at 95°C since this leads to aggregation of CLN8 and prevents the protein from entering the gel. SDS-gel electrophoresis was performed in Tris-glycine buffer (20 mM Tris, 160 mM glycine, 0.083% SDS w/v) using a 10% polyacrylamide gel. The Color Prestained Protein Standard Broad Range (P7719S, New England Biolabs) was used as molecular weight reference. Proteins were transferred onto a PVDF membrane via the wet blotting method in CAPS buffer (10 mM CAPS, 10% v/v, methanol, pH 11). The membrane was blocked with 10% blotting grade milk powder diluted in Tris-buffered saline with Tween20 (20 mM Tris, 150 mM NaCl, 0.1% Tween 20, pH 7.4). Primary antibodies and antibody-host targeting secondary antibodies used for immunological detection and are listed in **Supplementary table 3**.

### Site directed mutagenesis

Mutagenesis was performed using the Q5 site-directed mutagenesis kit (E0552S, New England Biolabs, Frankfurt am Main, Germany) according to the manufacturer’s instructions using the following primers: forward: 5′-GTTTCACTGCtgcATGGTTCTAA-3′; reverse: 5′- ATGTGAATCATCAGCCAC -3′. This mutation exchanges the conserved arginine 204 with a cysteine (R204C). Annealing temperatures were set according to the NEBaseChanger tool. Elongation time was set to 310 sec (∼20–30 s/kb). Constructs were transformed into chemically competent *Escherichia coli*. The base-exchange was verified by Sanger sequencing.

### Lysosome immunopurification

To enable lysosome immunopurification (lyso IP), we first generated Hek293 WT and *CLN8*-KO cell lines stably expressing the lysosomal transmembrane protein TMEM192-HA using lentiviral transduction as previously described^45^. To generate lentiviral particles, Hek293T cells (CRL-3216, ATCC) were transfected with the lentiviral packaging vectors psPAX2 and pMD2.G (both gifts from Didier Trono, Addgene plasmids #12260 and #12259) as well as the pLJC5-Tmem192-3xHA (gift from David Sabatini, Addgene plasmid #102930). Hek293 WT and *CLN8*-KO cell lines expressing TMEM192-3xHA, were maintained in culture medium additionally supplemented with 2 µg/ml puromycin to ensure stable protein expression.

Lyso IP was performed according to published protocols with minor adaptions^24^. First, Hek293 WT and *CLN8*-KO cells (pool of two *CLN8*-KO clones) were seeded in 145 mm dishes at a density of 1×10^7^ cells per dish. After reaching confluency, cells were washed twice with 5 ml ice-cold PBS and harvested in 950 µl of ice cold KPBS (136 mM KCl, 10 mM KH_2_PO_4_, pH 7.25) using a cell-scraper. The cells were centrifuged at 1,000 x *g* for 2 min at 4°C and resuspended in 950 µl fresh KPBS. Thereafter, 25 µl of this suspension were snap frozen in liquid N_2_ as the whole-cell fraction. With the remaining solution, cell lysis was performed by aspirating it 25 x up and 25 x down through a 26G needle. The 1,000 x *g* supernatant (centrifugation for 2 min at 4°C) was transferred into a low protein binding tube (90410, Thermo Scientific) containing 50 µl of prewashed Pierce^TM^ Anti-HA-magnetic beads (88837, Thermo Scientific). After a 3-5 min incubation on an orbital shaker, the tubes were placed on an EasyEights^TM^ EasySep^TM^ magnet (18103, STEMCELL Technologies) and the supernatant was removed. The beads were washed twice with 950 µl KPBS and transferred into a fresh low protein binding tube. Beads were washed one more time and subsequently transferred into a 2 ml safe-lock tube. The supernatant was discarded and purified lysosomes were snap frozen in liquid N_2_ for lipid extraction or Western blot analysis. All procedures except the incubation on the orbital shaker were performed on ice. The lyso IP protocol was performed with 4 samples in parallel (2x WT, 2x *CLN8*-KO). For the Western blot analysis, whole-cell fractions and purified lysosomes were resuspended in 100 µl RIPA buffer (150 mM NaCl, 10 mM Tris, 0.5 mM EDTA, 0.5% NP-40, and protease inhibitor cocktail (1 µg/ml pepstatin, 2 µg/ml antipain, 20 µg/ml leupeptin) and incubated on ice for 1h. The samples were centrifuged at 15,000 x *g* for 10 min at 4°C, and the supernatant was transferred into a fresh low protein binding tube. Finally, 15 µl of each fraction (whole cell and purified lysosomes) were used for Western blot analysis.

### Preparation of cell lysates

Pulverized tissues samples (∼20 mg) were homogenized using an T10 basic Ultra-Turrax (IKA) or with ceramic beads (1.4 mm) in a bead mill homogenizer for 30 sec in 0.5 ml lysis solution (250 mM sucrose, 1 mM EDTA, 1 mM DTT) supplemented with protease inhibitor (1 µg/ml pepstatin, 2 µg/ml antipain, 20 µg/ml leupeptin). The cells were lysed in 0.2-0.5 ml lysis solution using the SONOPULS ultrasonic homogenizer (Bandelin) for 3 x 10 sec with an amplitude of 20%. The cell and tissue homogenates were centrifuged at 1,000 x *g* for 10 min (4°C) to remove nuclei and debris. The supernatants (infranatant for adipose tissue lysates) were collected and adjusted for equal protein content by determining the protein concentration using the Bradford reagent.

### *In vitro* activity assays

GPGAT assays with Expi293 cell lysates containing recombinantly expressed GPAT1-4 were performed under conditions optimized for GPAT enzymes^47^. To this end, 20 µl of the cell lysates (2 mg/ml protein concentration) were incubated for 30 min at 37°C with 180 µl substrate solution containing 75 mM tris-HCl buffer (pH 7.5) supplemented with 800 µM [^13^C]GPG^+3^ as acyl-acceptor, 80 µM 16:0-, or 18:1-CoA as acyl donor, 1 mM DTT, 1 mg/ml FA free BSA, and 4 mM MgCl_2_.

All other GPGAT activity assays were performed with a substrate solution containing 100 mM bis-tris-propane buffer (pH 7) supplemented with 1% FA-free BSA, 1 mM DTT, 100 µM oleoyl-CoA as acyl donor and 100 µM [^13^C]GPG^+3^ as acyl-acceptor, unless otherwise stated.

As source of enzymatic activity, we used 20 µl of tissue/cell lysates (1 mg/ml protein concentration) or 6 µl of purified CLN8 or CLN8 R204G (1µg each), which we incubated with 80 µl substrate solution for 1h at 37°C in a water bath under mild agitation. Assays with purified proteins and Expi293 cell lysates containing recombinantly expressed CLN8 were incubated for 20 min at 37°C to ensure linearity of the reaction. For kinetic measurements, 16:0-, 18:1-, or 18:2-CoA were used at increasing concentrations, while the concentration of [^13^C]GPG^+3^ remained at 100 µM. Comparison of GPGAT and LPGAT activity of purified CLN8 was performed using either 100 µM [^13^C]GPG^+3^ or 100 µM *sn*-1-oleoyl-*R,rac*-LPG (858125, Avanti Polar Lipids) as acyl-acceptor. To measure the stereospecificity of CLN8, 4 µg of purified CLN8 protein (24 µl) were incubated for 1h at 37°C with a substrate containing 200 µM [^13^C]GPG^+3/PG^ as acyl acceptor, which was generated via saponification of head group labeled PG as described above. The determination of the head group specificity of purified CLN8 was performed by using 100 µM of commercially available GPE (Cay34460, Cayman Chemical), GPC (G5291, Sigma-Aldrich), or custom generated [^13^C]GPG^+3^, GPI, and GPS as acyl acceptor. GPI and GPS were prepared via saponification of PI and PS as described above for [^13^C]GPG^+3/PG^.

To investigate whether LPGAT1 or CLN5 can utilize the CLN8 reaction product as a substrate for PG or BMP synthesis, we first prepared Expi293 lysates containing recombinantly expressed CLN5 and sonicated membrane preparations of Expi293 cells containing recombinantly expressed LPGAT1 or CLN8, and used these as a source of enzymatic activity. To isolate membranes, cell lysates (2 mg/ml protein concentration) were centrifuged at 100,000 x *g* for 1h. The membrane pellet was subsequently emulsified in equal amount of lysis solution via sonication. For the assay, 20 µl of membranes containing recombinantly expressed CLN8 were first incubated with 80 µl substrate containing 100 µM oleoyl-CoA and [^13^C]GPG^+5^ as described above. After 20 min, the reaction was heat inactivated for 10 min at 85°C and centrifuged at 10,000 x *g* for 3 min to pellet the protein precipitate. Next, the reaction, including the precipitate was further combined with either 20 µl of recombinant LPGAT1-containing membranes, or 20 µl Expi293 cell lysates containing recombinant CLN5. The reactions with LPGAT1 were additionally supplemented with 10 µl substrate containing 18:1-CoA (80 µM) as acyl donor with and without *sn*-1-oleoyl-*R,rac*-LPG (8 µM) as acyl acceptor. For reactions with CLN5, the assay pH was adjusted to ∼6 by adding 80 µl 100 µM MES buffer (pH 5.5). Both, LPGAT1 and CLN5 reactions were incubated for another 20 min at 37°C.

All activity assays were stopped either by snap freezing in liquid N_2_ or by the addition of lipid extraction solution (see lipid extraction). The reaction products were analyzed by LC-MS.

### Lipid and GPD extraction

#### Lipid extraction

Samples of *in vitro* assays, cell pellets, and pulverized tissues (10-20 mg) were extracted in 2 ml safe lock tubes using a modified protocol of Matyash *et al.*^48^ with 1 ml extraction solution containing methyl *tert*-butyl ether/methanol (3/1, v/v), 0.01% (w/v) butylated hydroxytoluene, 0.01% (v/v) acetic acid, and the following internal standards: 150 pmol 17:1 LPG (858127), 150 pmol 14:0-14:0 BMP (857131), 150 pmol 14:0-14:0-14:0 hemi-BMP (857132), 150 pmol 14:0-14:0 PG (840445), 133 pmol 17:0-17:0 PE (830756), 30 pmol 17:0-17:0 PS (840028), 8 pmol 17:1 LPC (855677), 30 pmol 17:1 LPE (856707), 50 pmol 17:1 LPS (858141, all Avanti Polar Lipids), 50 pmol 17:0-17:0 PC (37-1700, Larodan). Total lipid extraction was performed under constant shaking for 30 min at RT. After the addition of 200 μl ddH_2_O (100 µl for *in vitro* assays) and further shaking for 10 min at RT, samples were centrifuged at 14,000 rpm for 5 min at RT to enable phase separation. For the isolation of lipids, 800 µl of the upper organic phase were collected and dried under a stream of N_2_. For LC-MS analysis, the lipids were resolved in 100 µl 2-propanol/methanol/ddH_2_O (7/2.5/1, v/v/v).

#### GPD extraction

To extract cellular GPDs, the extraction solution (see above) was supplemented with 50 pmol (per ml) custom synthesized *sn*-[U-^13^C]glycero-3-phosphocholine ([^13^C]GPC^+3^) as internal standard. The extraction was performed as described above. After phase separation, the upper organic phase was quantitatively removed, and 200 µl of MeOH were added to the aqueous phase followed by a brief vortex. The mixture was centrifuged at 20,000 rpm for 5 min, and 350 µl of the supernatant were transferred into a fresh tube. The solution was dried under a stream of N_2_ at 45°C and reconstituted in 50 µl MeOH for LC-MS analysis.

The extracted cell proteins from lipid and GPD isolations were dried and solubilized in NaOH/SDS (0.3 M/0.1% w/v) at 65°C, and the protein content was determined with the Pierce BCA reagent (23225, Thermo Fisher Scientific) and used for normalizing the of LC-MS measurements.

### Targeted LC-MS analysis

Liquid chromatography was performed using an Agilent 1290 Infinity II UHPLC equipped with an ACQUITY UPLC BEH C18 Column 2.1 × 150 mm, 1.7 µm (186002353, Waters Corporation) for separation of lipids, or with a Kinetex HILIC column 100 × 2.1 mm, 1.7 µm, 100 Å (00D-4474-AN, Phenomenex) for GPD analysis. The column temperature was set to 50°C, and 2 µl sample volume were injected.

For the analysis of polyglycerophospholipids – including LPG, BMP, PG, hemi-BMP, and CL – the mobile phases consisted of MeOH/H₂O (8:2, v/v) as phase A and 2-propanol/MeOH (8:2, v/v) as phase B, both containing 10 mM ammonium acetate, 0.1% formic acid, and 8 µM phosphoric acid. Chromatographic separation was carried out at a flow rate of 0.2 ml/min over a 30 min gradient. The gradient started with a linear increase from 50% to 60% phase B over 13 min, followed by a linear ramp to 100% phase B over 7 min. This was maintained isocratically for 5 min before re-equilibration at 50% phase B for 5 min.

Shorter chromatography gradients, using the same mobile phases, were applied depending on the specific experimental setup and targeted lipid species. For samples targeting LPG, BMP, and PG, an 18 min gradient was used. For *in vitro* assays where only LPG was analyzed, a 10 min gradient was employed. In both shortened protocols, the runs began at 50% phase B, followed by a linear increase to a higher B percentage, an isocratic hold, and a final re-equilibration step to restore initial conditions.

For the analysis of glycerophospholipids – including LPC, LPE, LPS, PC, PE, and PS – separation was performed using water (H₂O, mobile phase A) and 2-propanol (mobile phase B), both supplemented with 10 mM ammonium acetate, 0.1% formic acid, and 8 µM phosphoric acid. The flow rate was set to

0.15 ml/min. The 30 min gradient began at 50% B for 0.5 min, increased to 80% B over 8.5 min, then ramped to 100% B over the following 13 min. The column was held at 100% B for 2.5 min before returning to 50% B for re-equilibration.

For GPD measurements mobile phase A consisted of 10 mM ammonium acetate and 0.05% formic acid in LC–MS grade water, while mobile phase B was 100% LC–MS grade acetonitrile. Elution was carried out isocratically at 50% B for 6 min at a flow rate of 0.22 mL/min.

All analytes were detected using an Agilent 6470 triple quadrupole mass spectrometer equipped with an Agilent Jet Stream electrospray ionization (ESI) source. Data acquisition was performed using MassHunter Data Acquisition software (version B.10, Agilent Technologies).

Lipid species including LPG, BMP, PG, hemi-BMP, and cardiolipin were analyzed in negative ion mode, whereas LPC, LPE, LPS, PC, PE, PS and monoacylglycerol fragments of LPG were measured in positive ion mode.

GPDs were analyzed using fast polarity switching, allowing simultaneous acquisition of positive and negative ions in a single run. Unit resolution was used in both MS1 and MS2 stages. All precursor-to-product ion transitions and collision energies are detailed in **Supplementary table 4**.

Lipid and GPD identification were performed using MassHunter Qualitative Analysis software (version 10.0, Agilent Technologies), and quantified using Skyline (version 24.1.0.414). Data were normalized to analyte-to-internal standard ratios and reported as arbitrary units (AU) per mg of protein. Where applicable, absolute quantification was achieved using external calibration with 18:1-18:1 BMP or 18:1 LPG (Avanti Polar Lipids), and results were expressed as pmol per mg protein.

For lipidomic measurements of tissues from WT, *Cln8*^WT/*mnd*^, *Cln8^mnd^*^/*mnd*^ mice, the signal-to-noise (S/N) ratios were calculated for each lipid species using Agilent MassHunter Quantitative Analysis 10.1 (Agilent Technologies), with the *Peak-to-Peak from Drift* signal-to-noise algorithm. Noise was determined as the peak-to-peak difference in the baseline region near each species, corrected for slow baseline drift. S/N ratios were calculated as the ratio of peak apex height to the baseline noise. Manual adjustment of integration boundaries was performed where necessary, and S/N values were recalculated after such adjustments. S/N data is provided in the source data. Peaks with S/N < 3 were considered below the limit of detection.

To calculate isotope-tracer incorporation, the endogenous peak (m/z +0) of each lipid species was multiplied by the corresponding natural isotope distribution factor, determined by the Agilent MassHunter Isotope distribution calculator (Version 10.0.10305.0). The resulting value was subtracted from the measured isotope peak to correct for natural isotope distribution. For cell culture experiments, the isotope factor-corrected signals from control samples were subtracted from those of isotope-tracer-treated samples. The relative amount of isotope labeled lipids was determined by calculating the ratio of labeled lipids to total lipids (sum of endogenous and labeled lipids).

### Mass spectrometry proteomics

Total cell lysates (5% SDS, 50mM AMBIC) were processed using S- Trap™ micro columns (C02-micro-80, Protifi) following the high recovery protocol recommended by the manufacturer and using trypsin (90059, Pierce) as the carrier protein. Mass spectrometry Samples were analysed on timsTOF ion mobility mass spec- trometer (Bruker) in-line a reversed-phase C 18 Aurora column (25 cm × 75 μm) on a UltiMate 3000 UHPLC system (Thermo) as described previously^49^. The spectra were recorded in DIA mode and processed with DIA-NN (DIA-NN 2.3.0 Academia) using a synthetic library created from UniProt protein database (human reviewed, 2024_07_29).

### Data visualization and statistics

Data visualization and statistical analysis were performed using *Prism* version 10.5.1. Data are presented as mean ± standard deviation (SD) unless stated otherwise. The exact number of replicates for each experiment (either biological replicates or independent samples) and the statistical test applied are reported in the figure legends. Significance levels are shown as *P*-values in the figures. *ChemDraw* version 23.1.1 was used to visualize chemical structures in Fig. 1a,d-h and 5b-d. *BioRender* was used to create the figure schematics in the illustrations for Figure 5a.

## Data availability statement

Source data of all figures are available as supplementary information. Further requests to.

## Supplementary information

Supplementary information is available for this paper. The source data file contains all data and uncropped blots reported in the main and extended data figures. The supplementary table file contains tables with information on chemicals, plasmids, antibodies, and targeted MS precursor-to product transitions. The supplementary methods file contains a detailed description of the synthesis of isotope-labeled compounds.

## Acknowledgments

We thank Davod Mahmudi for help with the mass spec proteomics sample preparation, Thomas Züllig and Gerald Rechberger for their support with the mass spectrometry investigations, and Heimo Wolinski for his help with the microscopy experiments. This work was supported by the Austrian Science Fund (FWF), Field of Excellence BioHealth – University of Graz, Province of Styria, City of Graz, BioTechMed-Graz, NAWI Graz, EMBO, Batten Disease Global Research Initiative (BDGRI) and NCL Foundation (NCL-Stiftung). Grant numbers: SFB Lipid Hydrolysis 10.55776/F73 (D.K., R.Z.); FWF 10.55776/P35532 (R.Z.); doc-fund “Molecular Metabolism” 10.55776/DOC50; FWF 10.55776/ESP2877024 (C.W.), FWF 10.55776/PAT3403323 (U.T.); EMBO Scientific exchange grant 11929 (J.B.). For open access purposes, the authors have applied a CC BY public copyright license to any author accepted manuscript version arising from this submission.

## Author contributions

J.B., D.K., R.B., and R.Z., conceptualized the study. J.B., C.Z., D.B., M.T., and N.F. developed the methodology. J.B., C.Z., D.B., M.T., A.P., C.W., U.T., G.G, and L.H. conducted the investigation. J.B. and R.Z performed formal analysis. C.H. and R.B. synthesized isotope-labeled compounds. N.F. developed the methodology for mass spectrometry analysis. J.B. and N.F. performed mass spectrometry measurements and data analysis. C.Z. generated *CLN8*-deficient cell lines. M.T. established microscopy experiments for the analysis of phospholipidosis in cells. U.S. performed proteomics measurements.

K.P. provided the *Cln8*-deficient *mnd* mice. K.T. conducted the animal experiments. J.B. visualized the data. J.B. and R.Z. wrote the original draft. A.L. and R.Z. supervised the study. J.B., C.W., U.T., K.P., D.K., and R.Z acquired funding.

## Materials and correspondence

Correspondence and material requests should be addressed to Robert Zimmermann, phone: +433163801914; Institute of Molecular Biosciences, University of Graz, Heinrichstraße 31, 8010 Graz, Austria.

