## Supplementary Methods for "CLN8 enables a non-canonical phospholipid synthesis pathway"

### Synthesis of (*sn, rac*-$^{13}C_3$)-GPG and (*rac*-$^{13}C_2$, *rac*-$^{13}C_3$)-GPG

#### Synthesis Pathway A:

The initial pathway targeting GPG-13C3 ([$^{13}$C]GPG$^{+3}$) required protection of the glycerol, which was performed by acetalization with benzaldehyde. The product was synthesized by sequential phosphoramidite coupling of the protected glycerol building blocks as shown in Scheme 1. The product was synthesized successfully, the tendency of the 1,2-benzylidene-glycerol intermediates to isomerize under the coupling conditions and in storage resulted in contamination of the final product with varying amounts of glycerol phosphate. (12% glycerol-$^{13}C_3$ phosphate in the final product GPG-$^{13}C_3$)

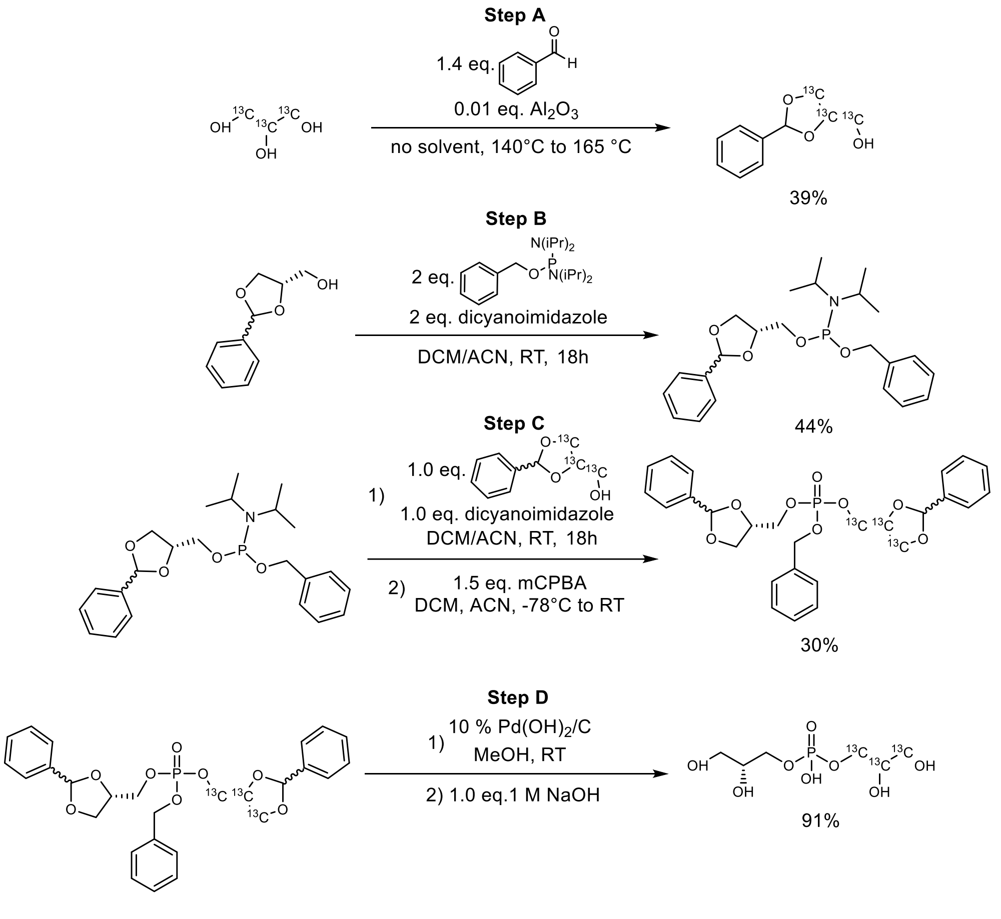


*Scheme 1: Original synthesis pathway to GPG-$^{13}C_3$*

The unlabelled chiral building block was synthesized as described in Scheme 2.

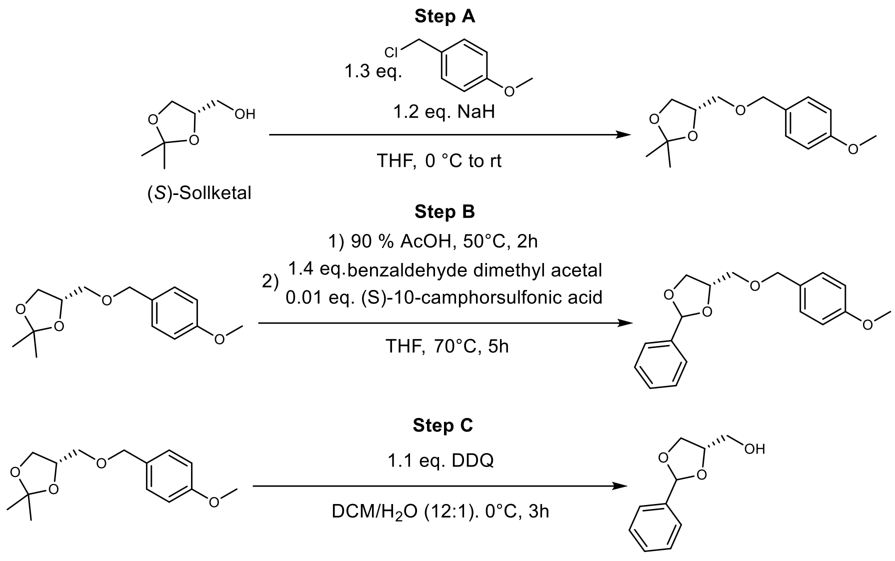


*Scheme 2: Synthesis of chiral (S)-1,2-benzylidene glycerol*

### Synthesis Pathway B:

Due to the lability of the 1,2-benzylidene acetal protecting group on glycerol, which complicates storage and handling of the building blocks and leads to partial product isomerization during the phosphoramidite couplings, a new benzyl protecting group strategy was devised for the synthesis of GPG-$^{13}C_5$. Although the selective synthesis of the intermediary 1,2-bis(benzyloxy)glycerol building blocks adds two additional steps (Scheme 3) to the reaction sequence, this synthetic pathway resolved the instability problems of the synthesis pathway A.

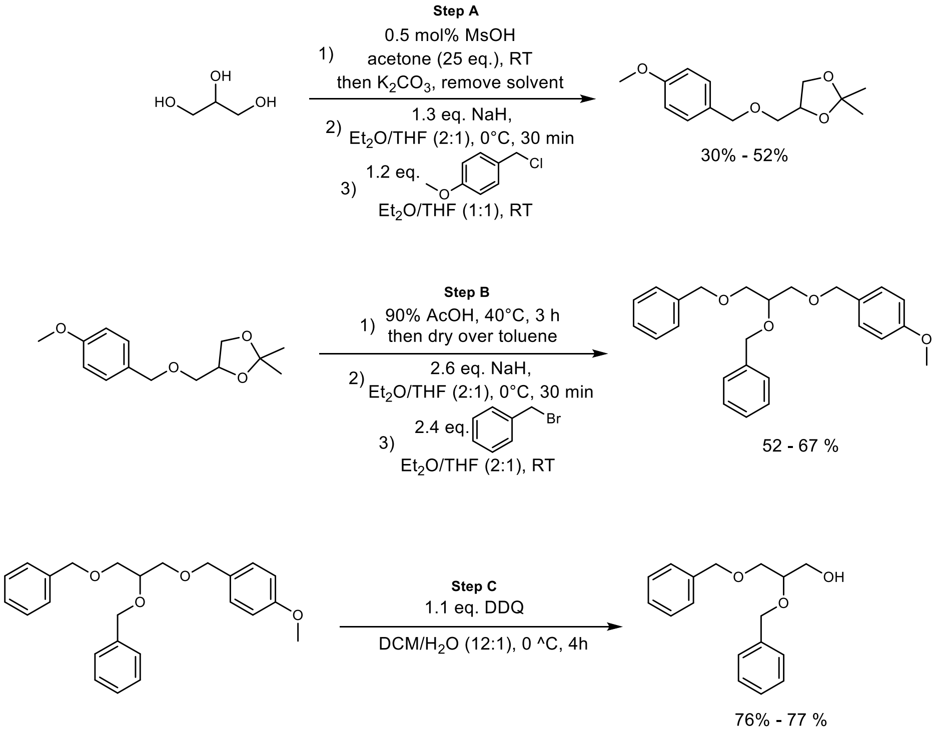


*Scheme 3: Synthesis of glycerol building blocks for path B*

The synthesis of (*rac,rac*)-GPG-$^{13}C_5$ ([$^{13}$C]GPG$^{+5}$) was performed using the same phosphoramidite coupling procedures as in (Scheme 4).

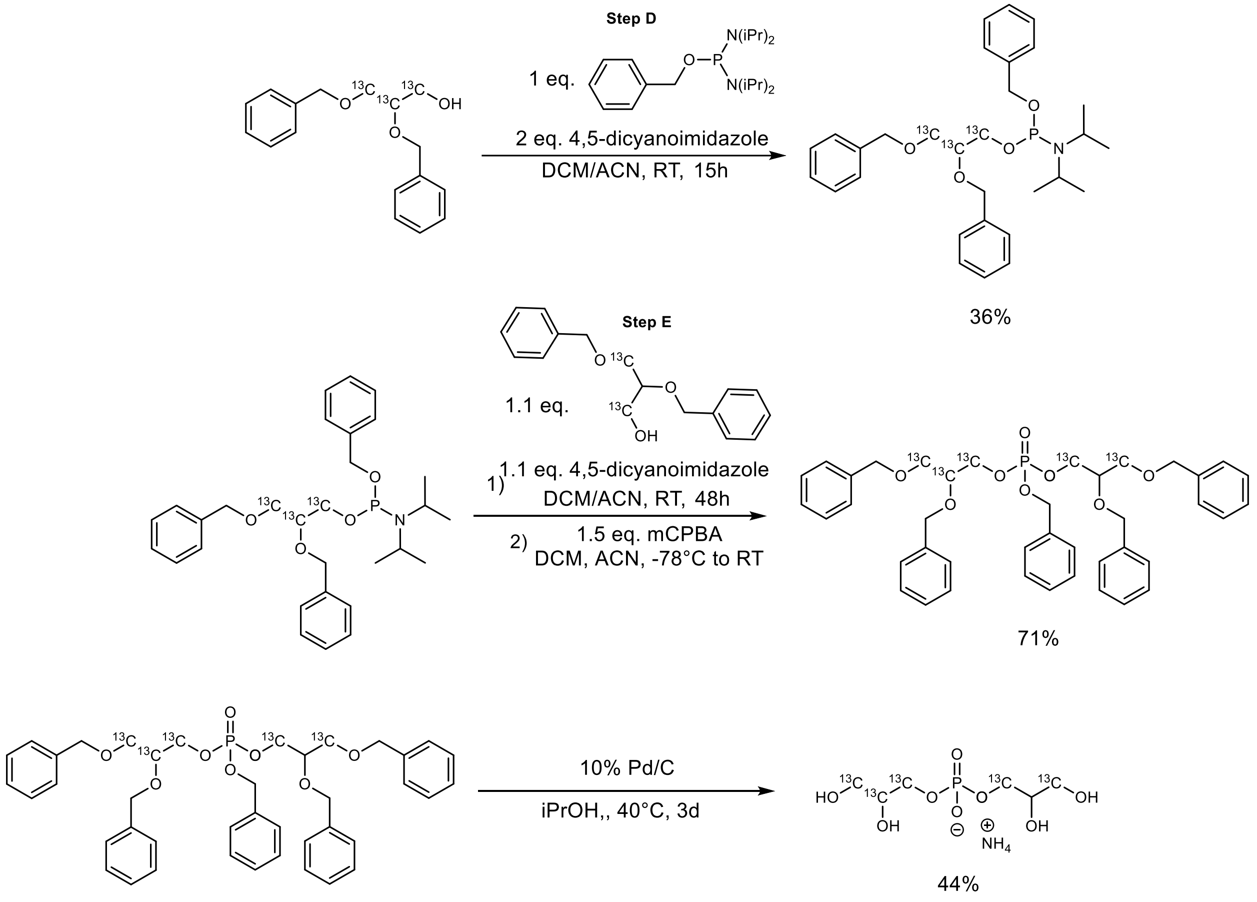


*Scheme 4: Phosphoramidite coupling of pathway B*

### General Information

Experiments were carried out under air with non-dry solvents unless otherwise mentioned. When applying Schlenk techniques the glass apparatus was dried under oil pump vacuum by heating with a heat gun, cooled to RT, and flushed with inert gas. In general, when high vacuum (*in vacuo*) was stated in experimental procedures, typically a vacuum of $10^{-2}$-$10^{-3}$ mbar was applied. Dry solvents were prepared by the below-mentioned procedures and afterwards stored under inert gas atmosphere (argon) over molecular sieves. In some cases, when explicitly mentioned, dry solvents were received from the listed suppliers. All reagents were added in a counter stream of inert gas to keep the inert atmosphere. All reactions were stirred with Teflon-coated magnetic stirring bars.

The stated temperatures generally refer to the oil bath or the cooling bath temperature. Temperatures were measured externally if not otherwise stated. When working at a temperature of 0 °C, an ice-water bath served as the cooling medium. Lower temperatures were achieved by using an acetone/dry ice cooling bath. Reactions, which were carried out at higher temperatures than rt, were heated in a silicon oil bath on a heating plate (RCT basic IKAMAG® safety control, 0-1500 rpm) equipped with an external temperature controller. The water bath temperature of the rotary evaporator was usually set to 40 °C unless otherwise noted.

Molecular sieves (Sigma-Aldrich, beads with 8-12 mesh) were activated in a round-bottom flask with a gas outlet adapter by heating them carefully in a heating mantle at level 1 at least for 24 h under high vacuum until complete dryness was obtained. These activated molecular sieves were stored at rt under argon atmosphere.

### Chemicals

All commercially available chemicals and solvents were purchased from abcr, Acros Organics, Alfa Aesar, Fluka, Honeywell, Merck, Roth, Sigma Aldrich, TCI, Thermo Fisher Scientific, VWR and used without further purification, unless otherwise stated.

*Water ($H_2O$):* If water was used as a solvent in a reaction or for workup, deionized water from an "ELGA Purelab Prima 7/15/30" ion exchanger was used.

*Triethylamine ($Et_3N$, anhydrous):* $Et_3N$ was dried over Na. It was distilled into an amber 1 L Schlenk bottle and stored over activated 4 Å molecular sieves under argon atmosphere.

### Dry Solvents

*Dichloromethane ($CH_2Cl_2$):* Dichloromethane (stabilized with EtOH) was purchased from Fisher Scientific, dried over phosphorus pentoxide, distilled and heated under reflux over $CaH_2$ for 24 h. It was distilled into an amber 1 L Schlenk bottle and stored over activated 4 Å molecular sieves under argon atmosphere.

*Ethanol (EtOH):* Ethanol was purchased in 25 L plastic cans from Merck and dried by treating with sodium and diethyl phthalate in an inert distillation apparatus and was then slowly heated until formation of hydrogen was observed as intensive evolution of gas. The reaction mixture was heated under reflux for 2 h, distilled and stored over activated 3 Å molecular sieves in a brown 1 L Schlenk bottle under argon atmosphere.

*Tetrahydrofuran (THF):* Tetrahydrofuran was purchased from VWR and heated under reflux over Na until benzophenone indicated dryness (intense blue color). It was distilled into an amber 1 L Schlenk bottle and stored over 4 Å molecular sieves under argon atmosphere.

### Analytical Methods

#### High Performance Liquid Chromatography with Mass Spectrometry (HPLC-MS)

Analytical HPLC-MS measurements were performed on an Agilent Technologies 1200 Series system (G1379 Degasser, G1312 Binary Pump, G1367C HiP ALS SL Autosampler, G1330B FC/ALS Thermostat, G1316B TCC SL column compartment, G1365C MWD SL multiple wavelength detector (deuterium lamp, 190-400 nm)) equipped with a single quadrupole LCMS detector "6120 LC/MS" using electrospray ionization source (ESI in positive and negative mode) or on a Shimadzu LCMS-2020 HPLC system with SCL-40 system controller, DGU-405 degassing unit, LC-40D XR solvent delivery module, SIL-40C XR auto sampler, SPD-40 UV-VIS detector, CTO-40C column oven, FCV-20AH2 valve unit and subsequent connected mass detector (Shimadzu LCMS-2020) with an electrospray ionization (ESI) source.

Separations on the Agilent Technologies 1200 Series system were carried out on a C18-Reversed-Phase column of the type „Poroshell® 120 SB-C18, 3.0 x 100 mm, 2.7 µm“ by Agilent Technologies. Flow: Constant flow rate 0.7 mL/min, T = 35 °C. The following method was used:

*MeCN_2_100_A*: 0.0 – 0.1 min, isocratic, 2% MeCN (98% $H_2O$ + 0.05% TFA); 0.1 – 8.0 min, linear, 2% to 100% MeCN (98% to 0% $H_2O$ + 0.05% TFA); 8.0 – 11.1 min, isocratic, 100% MeCN; 11.1 – 11.3 min, linear, 100% to 2% MeCN (0% to 98% $H_2O$ + 0.05 % TFA); 11.3 – 12.0 min, isocratic, 2% MeCN (98% $H_2O$ + 0.05% TFA).

Separations on the Shimadzu LCMS-2020 HPLC system were carried out on a Waters ACQUITY UPLC CSH C18 column (130 Å, 1.7 µm, 2.1 mm x 50 mm, 1/pk). Signals were detected at 288 nm. As mobile phase acetonitrile (VWR HiPerSolv, HPLC-MS grade) and water (Barnstead NANOpure®, ultrapure water system) with 0.05 % formic acid (HCOOH) were used. Flow: Constant flow rate 0.5 mL/min, T = 40 °C. The following method was used:

*MeCN_2_100_B*: 0.0-0.2 min, isocratic, 2% MeCN (98% $H_2O$ + 0.05% HCOOH); 0.2-6.5 min, linear, 2% to 100% MeCN (98% to 0% $H_2O$ + 0.05% HCOOH); 6.5-7.9 min, isocratic, 100% MeCN; 7.9-8.5 min, linear, 100% to 2% MeCN (0% to 98% $H_2O$ + 0.05 % HCOOH); 8.5-9.0 min, isocratic, 2% MeCN (98% $H_2O$ + 0.05% HCOOH).

#### Gas Chromatography with Mass Spectrometry (GC-MS)

GC-MS analyses were performed on an Agilent Technologies 7890A GC system equipped with a 5975C mass selective detector (inert MSD with Triple Axis Detector system) by electron-impact ionization (EI) with a potential of E = 70 eV. Herein, the samples were separated depending on their boiling point and polarity. The desired crude materials or pure compounds were dissolved, and the solutions were injected by employing the autosampler 7683B in a split mode 1/20 (inlet temperature: 280 °C; injection volume: 0.2 µL). Separations were carried out on an Agilent Technologies J&W GC HP-5MS capillary column ((5%-phenyl)methylpolysiloxane, 30 m x 0.2 mm x 0.25 µm) with a constant helium flow rate (He 5.0 (Air Liquide), 1.085 mL·min$^{-1}$, average velocity: 41.6 cm·s$^{-1}$). The following method was used:

*MT_50_S*: initial temperature: 50 °C for 1 min; linear increase to 300 °C (40 °C·min$^{-1}$); hold for 5 min; 1 min post-run at 300 °C; detecting range: 50.0-550.0 amu; solvent delay: 2.60 min.

#### Nuclear Magnetic Resonance Spectroscopy

NMR spectra were recorded on a Bruker Avance III 300 spectrometer ($^{1}$H: 300.36 MHz; $^{13}$C: 75.53 MHz) with autosampler, or a Jeol JNM-ECZL 400 MHz NMR Spectrometer ($^{1}$H: 399.78 MHz; $^{13}$C: 100.53 MHz, $^{19}$F: 376.17 MHz, $^{31}$P: 161.83 MHz).

Chemical shifts δ are referenced to the residual proton and carbon signal of the deuterated solvent ($CDCl_3$: δ = 7.26 ppm ($^{1}$H), 77.16 ppm ($^{13}$C); $CD_3OD$: δ = 3.31 ppm ($^{1}$H), 49.00 ppm ($^{13}$C); DMSO-$d_6$: δ = 2.50 ppm ($^{1}$H), 39.52 ppm ($^{13}$C); $D_2O$: δ = 4.79 ppm ($^{1}$H)). Chemical shifts δ are given in ppm (parts per million) and coupling constants *J* in Hz (Hertz). If necessary, 1D spectra (APT) as well as 2D spectra (H,H-COSY, HSQC, HMBC) were recorded for the identification and confirmation of the structure. Signal multiplicities are abbreviated as s (singlet), bs (broad singlet), d (doublet), t (triplet), q (quartet), quint (quintet), m (multiplet), dd (doublet of doublets), td (triplet of doublets), dt (doublet of triplets), and qd (quartet of doublets). Deuterated solvents for nuclear resonance spectroscopy were purchased from euriso-top®.

#### High Resolution Mass Spectrometry (HRMS)

High-resolution mass spectra (LC-ESI-MS/MS) were acquired by data-dependent high-resolution tandem mass spectrometry on a QExactive Focus (Thermo Fisher Scientific, Germany). The electrospray ionization potential was set to +3.5 or -3.0 kV, the sheath gas flow was set to 20, and an auxiliary gas flow of 5 was used. Samples were diluted with an appropriate solvent (methanol or chloroform) and 1 µL was injected on a SeQuant® ZIC®-pHILIC HPLC column (Merck, 100 x 2.1 mm; 5 µm; 100 Å; peek coated; equipped with a guard column) or on a RP-column (Waters, ACQUITY UPLC HSS T3 150 x 2.1 mm; 1.8 µm with VanGuard column). The separation solvent (pHILIC: A: $CH_3CN$, B: 25 mM $NH_4HCO_3$; RP: A: 0.1% HCOOH, B: 0.1% HCOOH in $CH_3CN$) was delivered through an Ultimate 3000 HPLC system (Thermo Fisher Scientific, Germany) with a flow rate of 100 µL·min$^{-1}$ and appropriate gradients were used for proper sample elution.

#### Determination of Melting Points

Melting points were determined on a Mel-Temp® melting point apparatus from Electrothermal with an integrated microscopical support. They were measured in open capillary tubes with a mercury-in-glass thermometer and were not corrected.

#### Thin Layer Chromatography

Analytical thin layer chromatography (TLC) was carried out on Merck TLC silica gel aluminum sheets (silica gel 60, F254, 20 x 20 cm). All separated compounds were visualized by UV light (λ = 254 nm and/or λ = 366 nm) and by the listed staining reagents followed by development in heat.

KMnO$_4$: 3.0 g $KMnO_4$ and 20 g $K_2CO_3$ were dissolved in 300 mL $H_2O$ and afterwards 5.0 mL 5% aq. NaOH were added.

CAM: 50 g $(NH_4)_6Mo_7O_{24}$, 2.0 g $Ce(SO_4)_2$ and 50 mL $H_2SO_4$ conc. were dissolved in 400 mL water.

#### Flash Column Chromatography

Flash column chromatography was performed on silica gel 60 from Acros Organics with particle sizes between 35 µm and 70 µm. Depending on the problem of separation, a 30 to 100-fold excess of silica gel was used with respect to the dry amount of crude material, if not otherwise stated. The dimension of the column was adjusted to the required amount of silica gel and formed a pad between 10 cm and 30 cm. In general, the silica gel was mixed with the eluent and the column was equilibrated. Subsequently, the crude material was dissolved in the eluent and loaded onto the top of the silica gel and the mobile phase was forced through the column using a rubber bulb pump. The volume of each collected fraction was adjusted between 20% and 30% of the silica gel volume.

#### Ion Exchange Chromatography for GPG-$^{13}C_5$

Ion-exchange chromatography was performed with Sephadex DEAE A-25 chloride form from Sigma Aldrich. 3.0g were conditioned to 0.1M $NH_4OAc$ buffer at pH 5 to a total volume of 20 mL. 32 mg of the contaminated product was applied as a solution in distilled $H_2O$, then washed with 100 mL $H_2O$. Elution of the compound was performed with 100 mL 0.5 M $NH_4OH$ to recover 22 mg of GPG-$^{13}C_5$ (69% recovery) and removal of the glycerol-$^{13}C_3$ impurity.

### Synthesis pathway A:

#### (*S*)-4-(((4-Methoxybenzyl)oxy)methyl)-2,2-dimethyl-1,3-dioxolane (1)

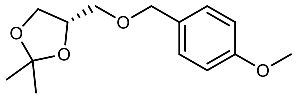

A Schlenk flask containing 4.60 g (34.8 mmol, 1.0 eq.) (s)-sollketal was evacuated and then backfilled with Ar (3x) followed by the addition of 40 mL dry THF. The reaction mixture was cooled to 0 °C in an ice-bath, followed by addition of 1.67 g (60% in mineral oil, 41.8 mmol, 1.2 eq.) NaH. A bubbler was connected to the Schlenk adapter and the suspension was stirred at 0 °C for 25 min until $H_2$-evolution stopped, which was accompanied by strong thickening of the solution. Subsequently, a solution of 498 µL (572 mg, 4.05 mmol, 1.2 eq.) PMB-Cl in 20 mL $Et_2O$ was added over the course of 10 min and the ice-bath was removed. The reaction mixture was stirred for 16h and excess NaH was quenched by slow addition of 30 mL $H_2O$, then 100 mL brine. The mixture was extracted with 2 x 80 mL EtOAc and the combined organic phases were dried over $Na_2SO_4$. The volatiles were removed under reduced pressure to yield a yellow oil, which was further purified via column chromatography (400 mL $SiO_2$, CH/EE = 10:1, CAM, Rf = 0.20) yielding 3.36 g (38% yield) of a colorless oil.

| | |
|---|---|
| **$C_{12}H_{20}O_4$** | [252.3 g/mol] |
| **Yield** | 3.36 g (13.3 mmol, 38%), colorless oil |
| **TLC** | $R_f$ = 0.20 (CH/EtOAc = 10:1, UV and CAM) |
| **GC-MS** | $t_R$ = 6.483 min; m/z (%): 252 (2), 193 (13), 163 (12), 121 (100) |

**$^1$H-NMR** (300 MHz, $CDCl_3$) δ = 7.26 (d, *J* = 8.2, 2H), 6.88 (d, *J* = 8.3, 2H), 4.39 - 4.62 (m, 2H), 4.17 - 4.39 (m, 1H), 3.95 - 4.14 (m, 1H), 3.67 - 3.86 (m, 4H), 3.30 - 3.67 (m, 2H), 1.39 (d, *J* = 17.8, 6H).
**$^{13}$C-NMR** (76 MHz, $CDCl_3$) δ = 159.4 (s), 130.2 (s), 129.5 (s), 113.9 (s), 109.4 (s), 74.9 (s), 73.3 (s), 70.9 (s), 67.1 (s), 55.4 (s), 26.9 (s), 25.5 (s).

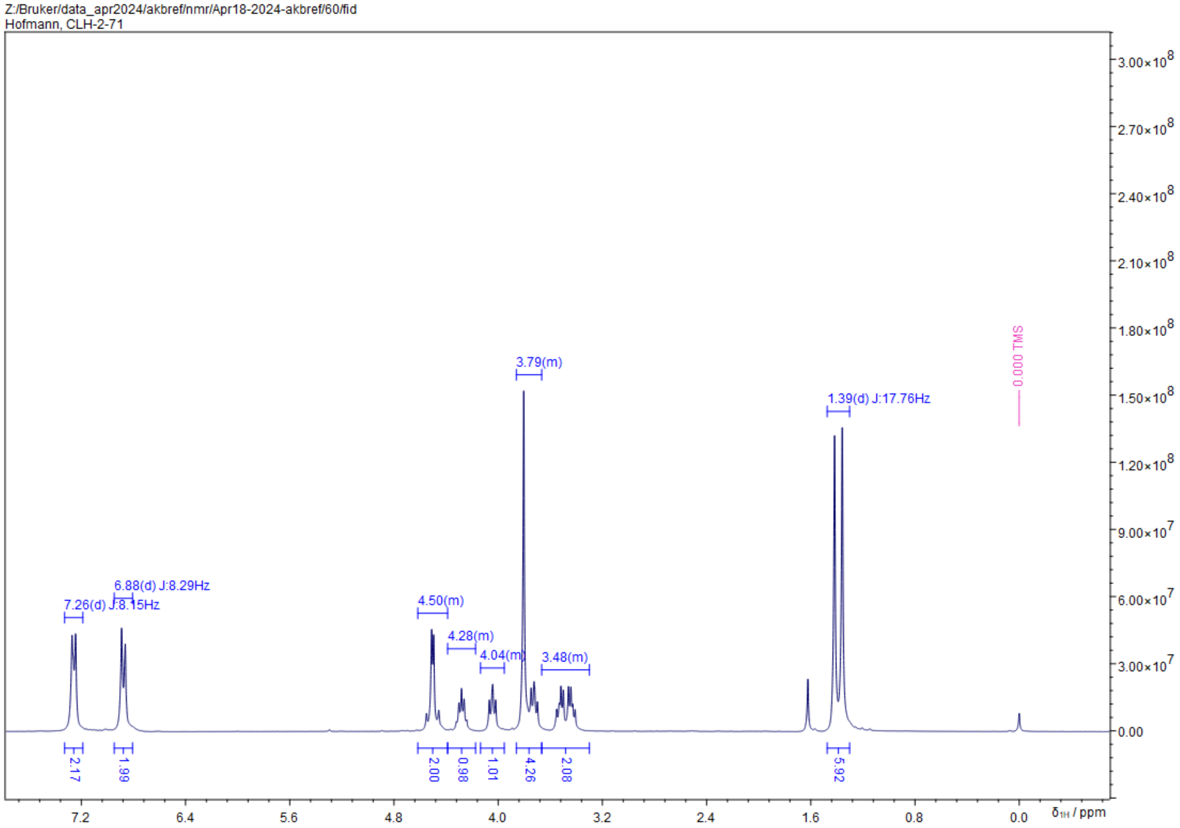
Z:/Bruker/data_apr2024/akbref/nmr/Apr18-2024-akbref/60/fid
Hofmann, CLH-2-71
7.26(d) J:8.15Hz
6.88(d) J:8.29Hz
4.50(m)
4.28(m)
4.04(m)
3.79(m)
3.48(m)
1.39(d) J:17.76Hz
0.000 TMS
2.17
1.99
2.00
0.98
1.01
4.26
2.08
5.92
δ1H / ppm

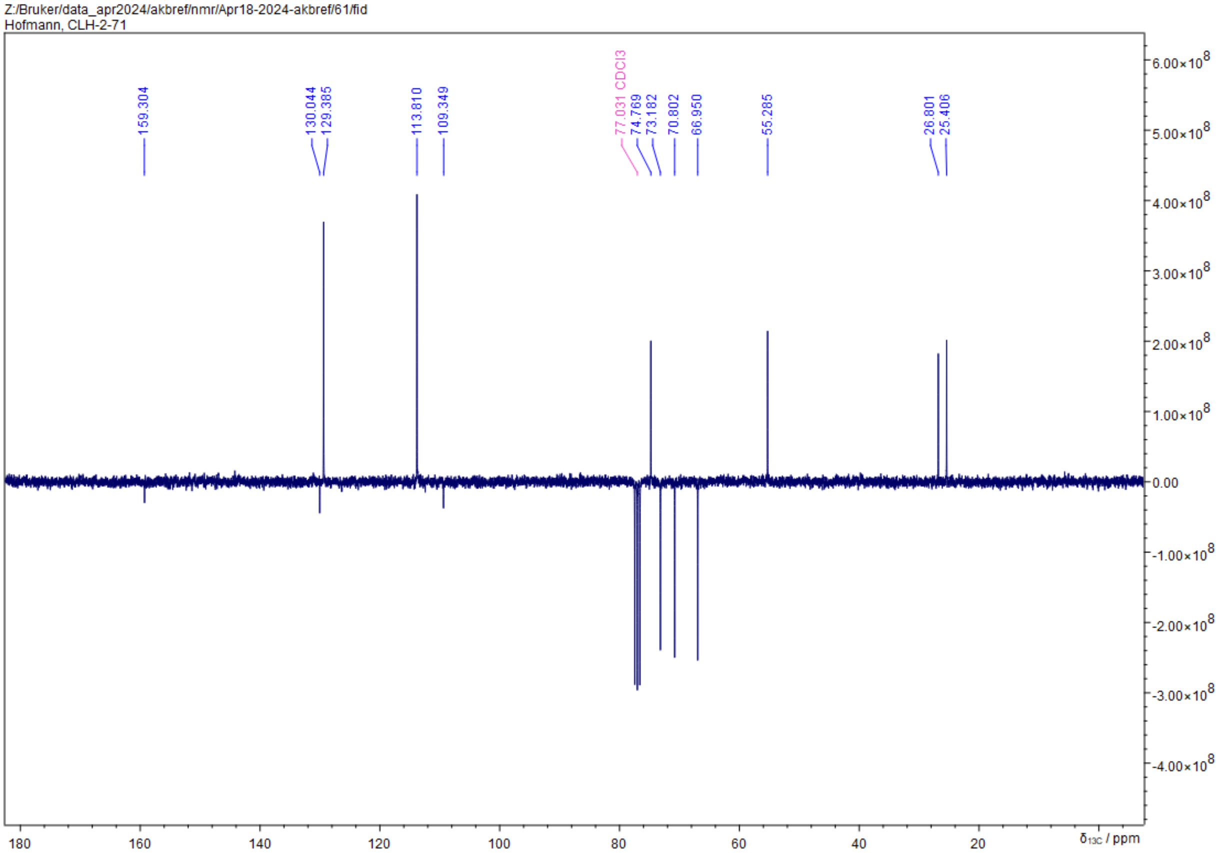
Z:/Bruker/data_apr2024/akbref/nmr/Apr18-2024-akbref/61/fid
Hofmann, CLH-2-71
159.304
130.044
129.385
113.810
109.349
77.031 CDCl3
74.769
73.182
70.802
66.950
55.285
26.801
25.406
δ13C / ppm

***cis/trans*-(4S)4-(((4-methoxybenzyl)oxy)methyl)-2-phenyl-1,3-dioxolane (2)**

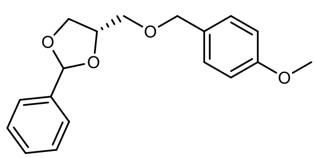

In a 50 mL round-bottom flask 3.31 g (13.1 mmol, 1.0 eq.) **1** were dissolved in 10 mL 90% acetic acid. The reaction was stirred at 50 °C until full conversion was indicated by HPLC (2 h). The volatiles were removed under reduced pressure to yield 2.77 g (13.1 mmol, quant.) pale yellow oil, which was used immediately in the next step without further purification. The intermediate was dissolved in 20 mL THF, followed by addition of 2.71 g (17.8 mmol, 1.4 eq.) benzaldehyde dimethylacetal and 31 mg (0.127 mmol, 0.01 eq.) (S)-10-camphorsulfonic acid. The mixture was stirred at 70°C for 5h, then quenched by addition of 20 mL sat $NaHCO_3$. After stirring for 5 min, the reaction was diluted with 5 mL $H_2O$ and extracted with EtOAc (2 x 20 mL). The combined organic phases were dried over $Na_2SO_4$. The volatiles were removed under reduced pressure and the resulting yellow oil was purified via column chromatography (50 mL $SiO_2$, CH/EE = 10:1) to yield 2.36 g (62%) of a colorless oil.

NMR spectra of cis/trans-isomers could not be resolved.

| | |
|---|---|
| **$C_{10}H_{12}O_3$** | [300.35 g/mol] |
| **Yield** | 2.36 g (7.85 mmol, 62%) |
| **TLC** | $R_f$ = 0.21 (CH/EtOAc = 10:1, UV and CAM) |
| **HPLC-MS 1** | $t_R$ = 6.736 min. |
| **HPLC-MS 2** | $t_R$ = 6.788 min. |

**$^1$H-NMR** (300 MHz, $CDCl_3$) δ = 7.15 - 7.50 (m, 6H), 6.86 (d, *J* = 6.6, 2H), 5.83 (d, *J* = 39.9, 1H), 4.50 (d, *J* = 7.6, 2H), 4.32 - 4.46 (m, 2H), 3.34 - 4.25 (m, 8H).

**$^{13}$C-NMR** (76 MHz, $CDCl_3$) δ = 159.5 (s, 1C), 138.0 (s), 137.4 (s), 134.5 (s) 130.1 (s), 129.9 (s), 129.5 (s), 129.5 (s), 129.5 (m), 129.3 (s), 129.1 (s), 128.4 (s), 126.8 (s), 126.6 (s), 114.0 (s), 104.5 (s,), 103.9 (s), 75.6 (s), 75.2 (s), 73.4 (s), 71.8 (s), 70.8 (s), 70.3 (s), 68.2 (s), 67.9 (s), 64.3 (s), 55.4 (s).

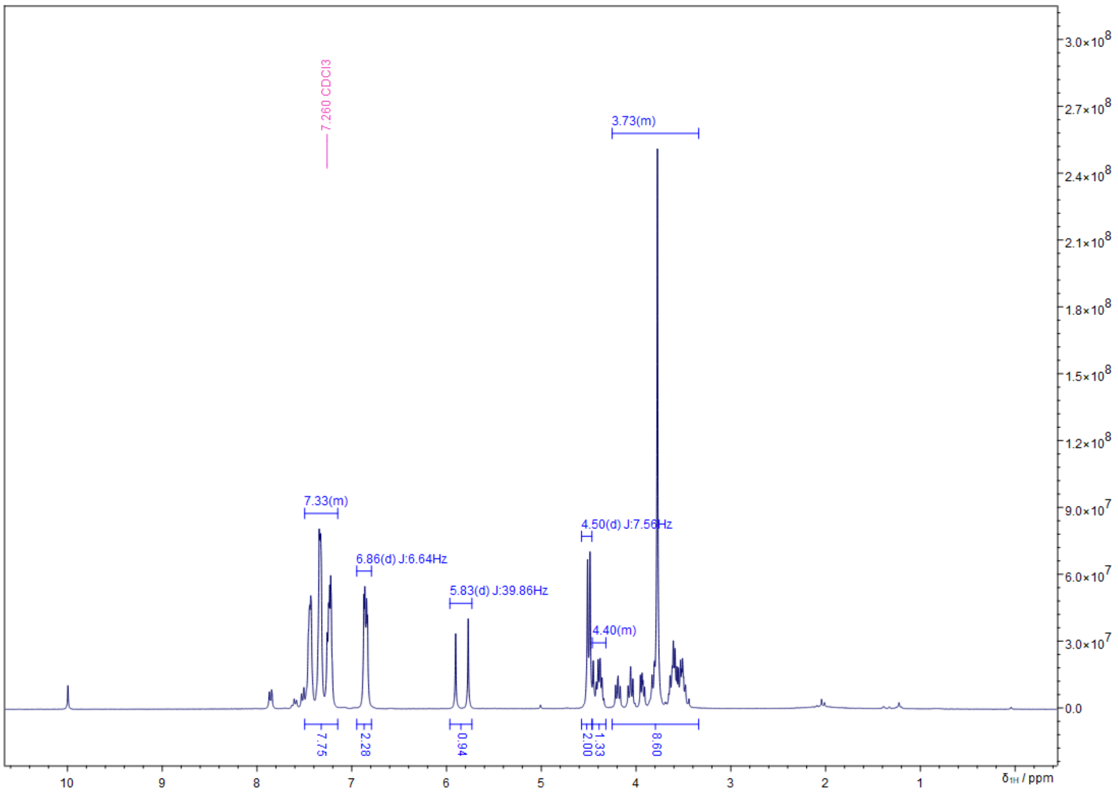
7.260 CDCl3
3.73(m)
7.33(m)
6.86(d) J:6.64Hz
5.83(d) J:39.86Hz
4.50(d) J:7.56Hz
4.40(m)
7.75
2.28
0.94
2.00
1.33
8.60
δ1H / ppm

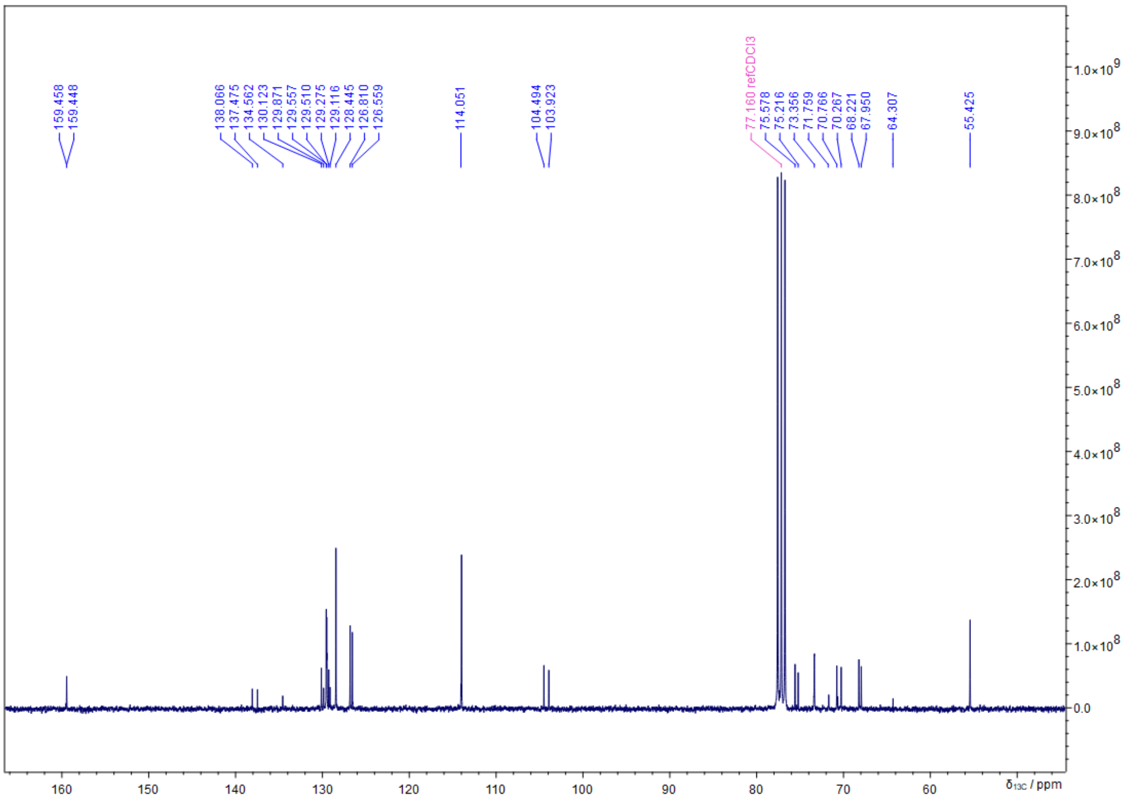
159.458
159.448
138.066
137.475
134.562
130.123
129.871
129.557
129.510
129.275
129.116
128.445
126.810
126.559
114.051
104.494
103.923
77.160 refCDCl3
75.578
75.216
73.356
71.759
70.766
70.267
68.221
67.950
64.307
55.425
δ13C / ppm

**(S)-(2-phenyl-1,3-dioxolan-4-yl)methanol (3)**

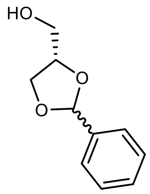

In a 50 mL round-bottom flask 260 mg (0.86 mmol, 1.0 eq.) **2** were dissolved in 4 mL degassed DCM and 200 µL $H_2O$. The reaction mixture was cooled to 0 °C in an ice bath, followed by the addition of 216 mg (0.95 mmol, 1.10 eq.) DDQ. The dark green emulsion was stirred at 0 °C and the progress of the reaction was accompanied by the formation of a light brown precipitate. After 3 h full conversion was indicated via HPLC-MS and the reaction was quenched by the addition of 5 mL sat. $NaHCO_3$ and extracted with DCM (3 x 10 mL). The combined organic phases were dried over $Na_2SO_4$. The volatiles were removed under reduced pressure and the resulting orange oil was purified via column chromatography (20 mL $SiO_2$, $Et_2O$/pentane = 1:1 to 2:1) to yield 137 mg (85%) of a light yellow oil.

| | |
|---|---|
| **$C_{10}H_{12}O_3$** | [180.20 g/mol] |
| **Yield** | 137 mg (0.821 mmol, 6%, *cis*/*trans* = *2:3*, 21%) |
| **TLC** | $R_f$ = 0.36 ($Et_2O$/pentane = 2:3, UV and CAM) |
| **GC-MS** (*cis*) | $t_R$ = 5.625 179 (76), 149 (35), 123 (12), 105 (100) |
| **GC-MS** (*trans*) | $t_R$ = 5.684 179 (83), 149 (21), 123 (100), 105 (62) |

**$^1$H-NMR** (400 MHz, $CDCl_3$) δ = δ = δ = 7.48 (s, 2H), 7.39 (s, 3H), 5.97 (s, *cis*-1,2-acetal, 1H, 39%), 5.83 (s, s, *trans*-1,2-acetal, 1H, 61%), 4.37 (s, 1H), 4.22 (t, *J* = 7.2, *cis*-1,2-acetal, 1H, 39%), 4.10 (t, *J* = 7.4, *trans*-1,2-acetal, 1H, 61%), 3.52 - 3.95 (m, 3H), 2.02 (d, *J* = 19.7, 1H).
**$^{13}$C-NMR** (101 MHz, $CDCl_3$) δ = 129.6 (s, *trans*), 129.4 (s, *cis*), 128.6 (s, *trans*), 128.5 (s, *cis*), 126.7 (s, *trans*), 126.5 (s, *cis*), 104.5 (s, *trans*), 103.9 (s, *cis*), 77.1 (s, *cis*), 76.7 (s, *trans*), 67.0 (s, *trans*), 66.9 (s, *trans*), 63.4 (s, *cis*).

Z:/Bruker/data_may2024/akbref/nmr/May23-2024-akbref/10/fid
Hofmann, CLH-2-78-2

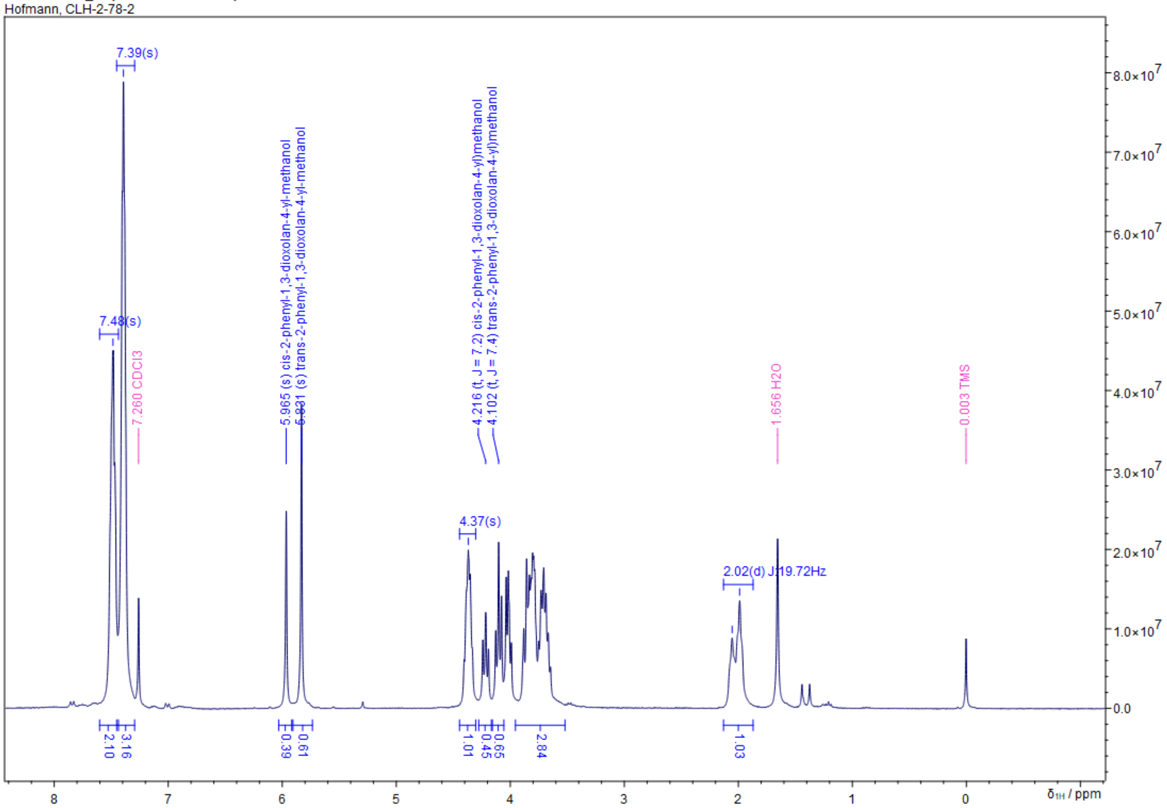


Z:/Bruker/data_may2024/akbref/nmr/May23-2024-akbref/11/fid
Hofmann, CLH-2-78-2

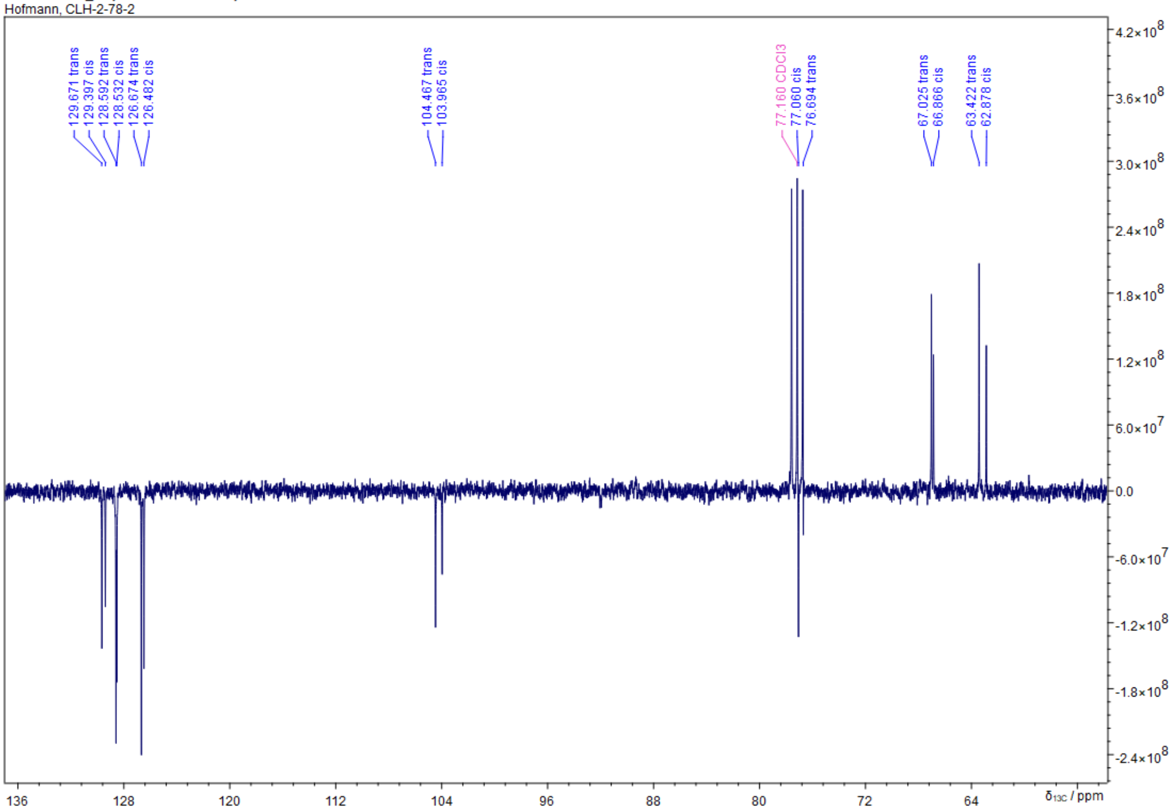

**rac-2-Phenyl-1,3-dioxolan-4-yl)methanol-$^{13}C_3$ (4)**

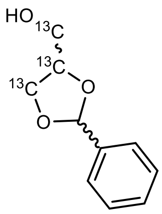

Into a 50 mL round-bottom were weighed 502 mg (5.45 mmol, 1.0 eq.) $^{13}C_3$-glycerol, 821 mg (7.74 mmol, 1.4 eq.) benzaldehyde and 5 mg acidic $Al_2O_3$ (0.05 mmol, 0.01 eq.). The flask was placed in an oil bath and was attached to a short-path distillation setup with the collecting flask being placed in an acetone/dry-ice bath, thus continuously removing $H_2O$ from the reaction and shifting the equilibrium towards the product. The reaction was subsequently heated to 145 °C and stirred for 2 h. As high reaction temperatures favor the reaction equilibrium towards the five-membered cyclic acetal, the mixture of condensation products was then heated to 165 °C for 30 min followed by immediate cooling in an ice-bath. The reaction was then diluted with 15 mL $Et_2O$, filtered and washed with sat. $NaHCO_3$ (5 mL), sat. $Na_2SO_3$ (10 mL) and $H_2O$ (5 mL). The aqueous phases were every time back-extracted with 5 mL $Et_2O$. The combined organic phases were dried over a 1:1 mixture of $MgSO_4$ and $K_2CO_3$ and the solvent removed under reduced pressure to obtain a crude yellow oil, which was purified via column chromatography (100 mL $SiO_2$, $Et_2O$/pentane = 1:1 to 3:1) to obtain 383 mg (39 % yield) of the 5-membered acetal products (*cis*/*trans*=2/3), as well as 59.2 mg (6 % yield) *trans*-isomer and 57.2 mg (6 % yield) *cis*-isomer of the six-membered acetal side-products. The product was immediately used as it was unstable. After 2 h in $CDCl_3$ at RT already 2% of product were isomerized in the resulting NMR spectrum as the authors were unaware of the DCl contamination of $CDCl_3$ at the time.

| | |
|---|---|
| **$^{13}C_3C_7H_{12}O_3$** | [183.20 g/mol] |
| **Yield** | 383 mg (2.38 mmol, 39%, *cis*/*trans* = *2:3*,), colorless oil |
| **TLC** | $R_f$ = 0.36 ($Et_2O$/pentane = 2:3, UV and CAM |
| **GC-MS** (*cis*-isomer) | $t_R$ = 5.623 min, 182 (76), 151 (36), 124 (13), 105 (100) |
| **GC-MS** (*trans*-isomer) | $t_R$ = 5.681 min, 182 (81), 151 (22), 124 (100), 105 (60) |

**$^1$H-NMR** (400 MHz, $CDCl_3$) δ = 7.44 - 7.53 (m, 2H), 7.34 - 7.43 (m, 3H), 5.97 (s, *cis*-1,2-acetal, 1H, **40%**), 5.835 (s, *trans*-1,2-acetal, 1H, **50%**); 5.56 (s, *cis*-1,3-acetal, 1H, **5%**), 5.43 (s, *trans*-1,3-acetal, 1H, **5%**), 4.08 - 4.61 (m, 2H), 3.73 - 4.08 (m, 2H), 3.37 - 3.72 (m, 1H), 1.88 (bs, 1H).

**$^{13}$C-NMR** (101 MHz, $CDCl_3$) δ = 129.7 (s, *trans*), 129.4 (s, *cis*), 128.6 (s, *trans*), 128.6 (s, *cis*), 126.7 (s, *trans*), 126.5 (s, *cis*), 104.5 (s, *trans*), 104.0 (s, *cis*), 77.1 (t, *J* = 34.5, *cis*, **$^{13}$C**), 76.7 (t, *J* = 34.8, *trans*, **$^{13}$C**), 67.0 (d, *J* = 33.7, *trans*, **$^{13}$C**), 66.9 (dd, *J* = 1.7, 34.0, *cis*, **$^{13}$C**), 63.4 (d, *J* = 41.6, *trans*, **$^{13}$C**), 62.9 (dd, *J* = 1.8, 41.1, *cis*, **$^{13}$C**)

Z:/Jeol-400/Backup_data/akbref/CLH-2-90_Säule2_PROTON-1-1.jdf
CLH-2-90_Säule2

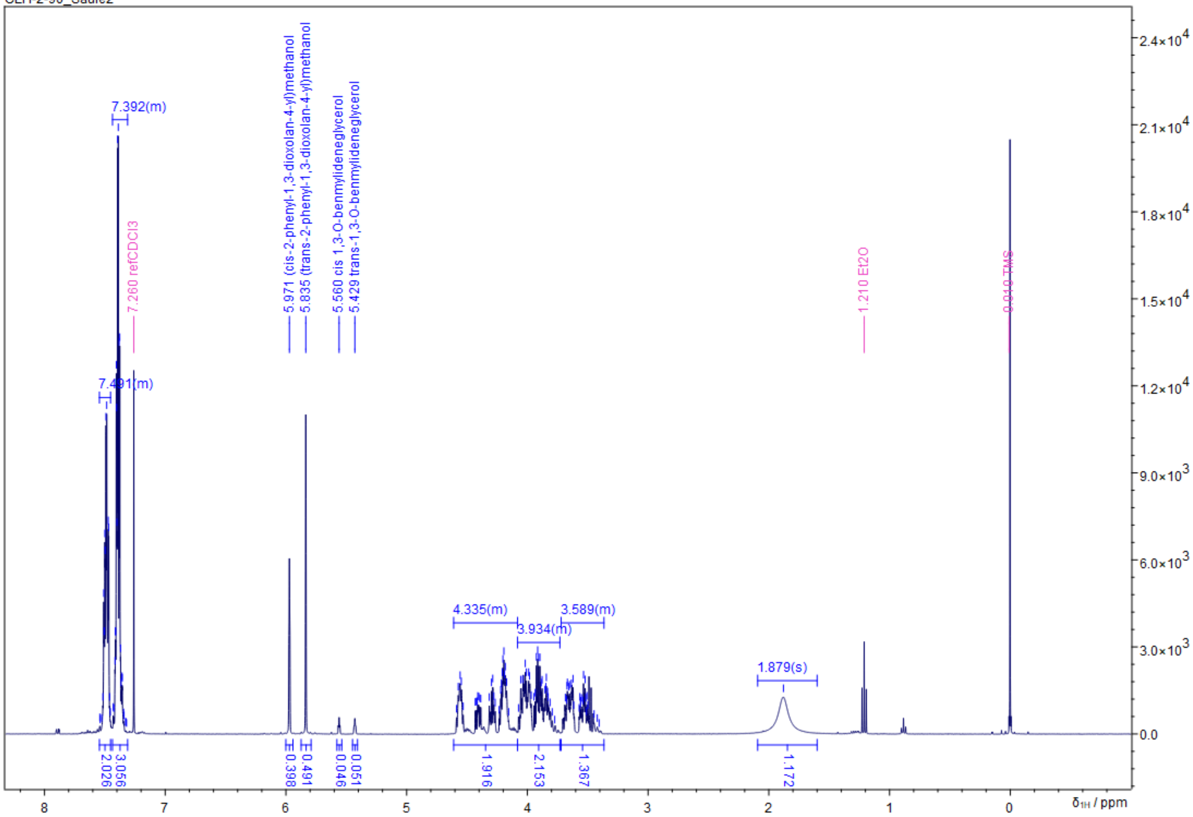


Z:/Jeol-400/Backup_data/akbref/CLH-2-90_Säule2_CARBON-1-1.jdf
CLH-2-90_Säule2

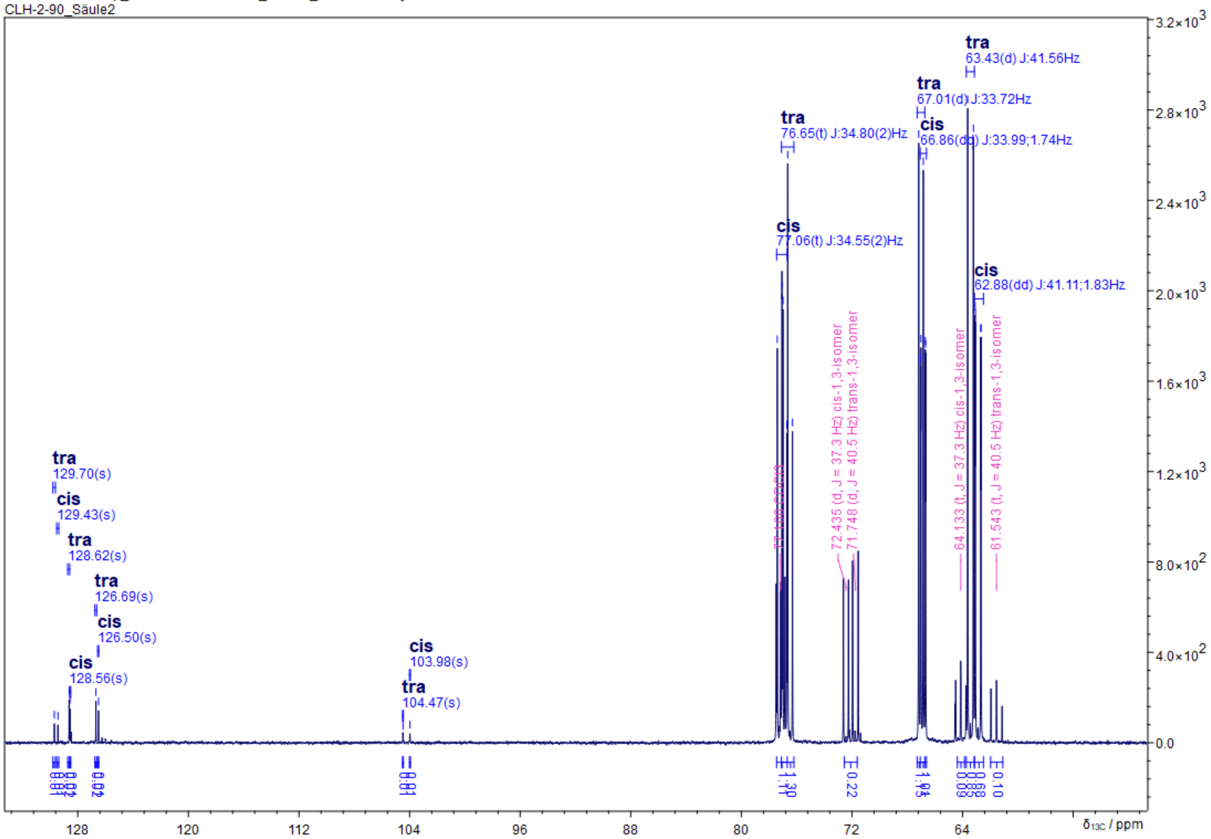

**Benzyl (*cis*/*trans*-((4R)-2-phenyl-1,3-dioxolan-4-yl)methyl) diisopropylphosphoramidite (5)**

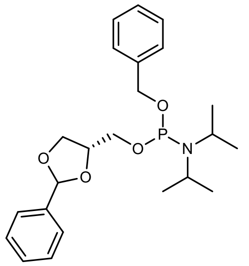


In an argon flushed Schlenk flask 55 mg (0.308 mmol, 1.0 eq.) **3** and 401 mg (0.771 mmol, 2.00 eq.) 1-(benzyloxy)-*N*,*N*,*N*',*N*'-tetraisopropylphosphanediamine were dissolved in 5 mL DCM, followed by the addition of 4 Å molecular sieves. After stirring at RT for 25 min, a solution of 73 mg (0.617 mmol, 2.00 eq.) 4,5-dicyanoimidazole in 2 mL ACN was added to the pre-dried reagents. After stirring at RT for 3 h, the reaction was diluted with 5 mL DCM and quenched by pouring the reaction onto 8 mL sat. $NaHCO_3$. After separation the aqueous phase was further extracted with 2 x 5 mL DCM. The combined organic phases were dried over $MgSO_4$ and the crude yellow oil was purified via column chromatography (20 mL $SiO_2$, CH/EE = 2:1 + 5% $Et_3N$, CAM, Rf = 0.85) to obtain 174 mg (81% theoretical yield) of the desired product as a yellow oil, however NMR analysis indicated contamination with 27 w% diisopropylamine, which was carried into the next synthesis.

| | |
|---|---|
| **$C_{23}H_{32}O_4P$** | [417.49 g/mol] |
| **Yield** | 127 mg + 47 mg diisopropylamine (0.28 mmol, 59%), yellow oil |
| **TLC** | $R_f$ = 0.82 (CH/EE = 2:1 + 5% $Et_3N$, UV and CAM) |

**$^1$H-NMR** (400 MHz, $CDCl_3$) δ = 7.28 - 7.44 (m, 10H), 6.5 (s, *cis*-1,3-benzylidene, 1H, 12%), 6.2 (s, *trans*-1,3-benzylidene, 1H, 87%), 4.91 - 4.7 (m, 4H), 3.37 - 3.61 (m, 5H), 1.96 (s, 1H), 1.11 - 1.32 (m, 12H).
**$^{13}$C-NMR** (101 MHz, $CDCl_3$) δ = 128.6 (d, *J* = 7.6), 128.2 (s), 127.7 (s), 127.0 (s), 65.1 (s), 45.3 (s), 44.6 (s), 23.7 (d, *J* = 2.7), 22.7 (s).
**$^{31}$P-NMR** (162 MHz, $CDCl_3$) = 149.0 (d, *J* = 4.4), 148.8 (d, *J* = 6.3).

Z:/Jeol-400/Backup_data/akbref/CLH-2-80-2_PROTON-1-1.jdf
CLH-2-80-2

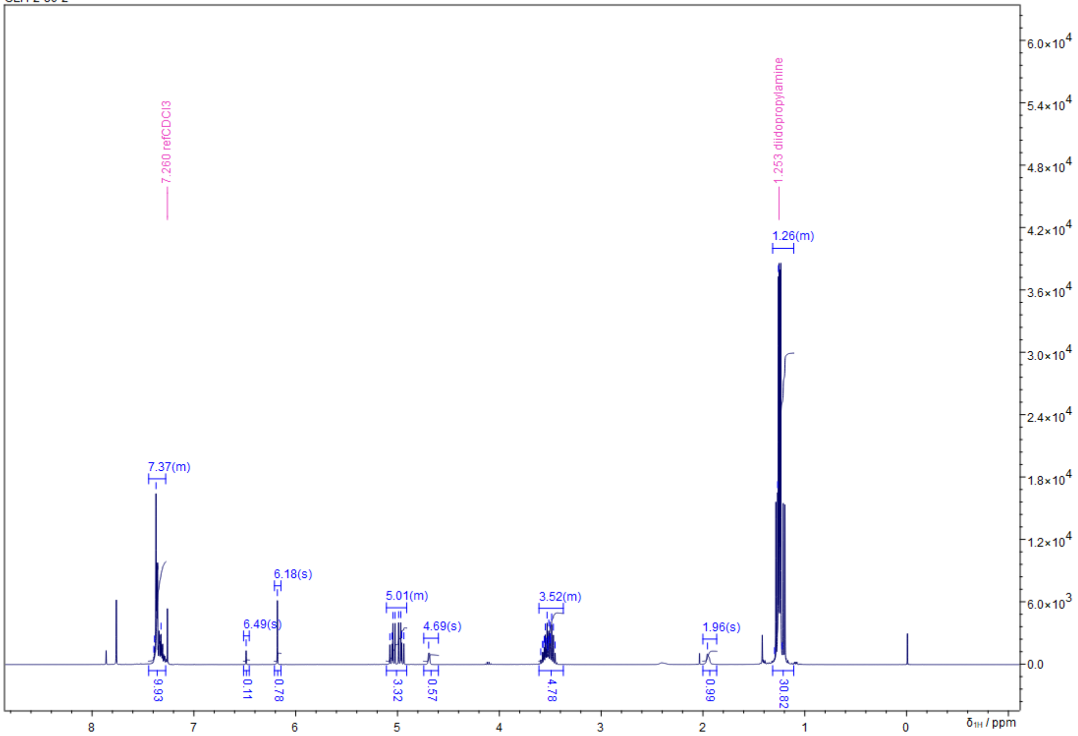


Z:/Jeol-400/Backup_data/akbref/CLH-2-80-2_CARBON-1-1.jdf
CLH-2-80-2

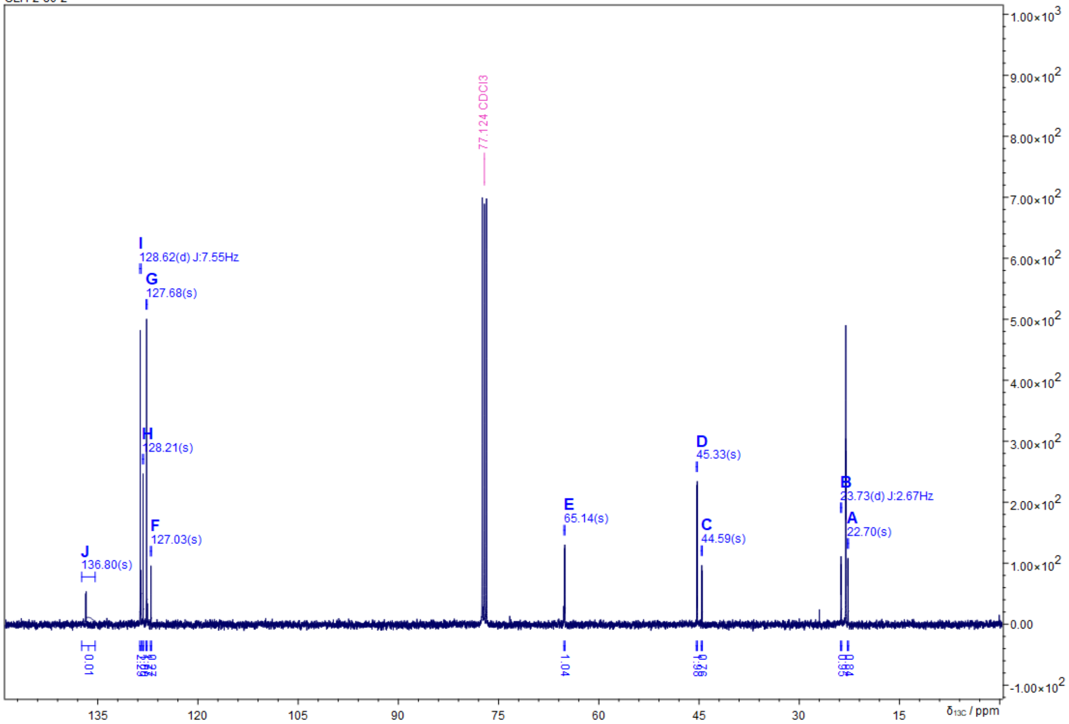

Z:/Jeol-400/Backup_data/akbref/CLH-2-80-4_Fr1-5_PHOSPHORUS-1-1.jdf
CLH-2-80-4_Fr1-5

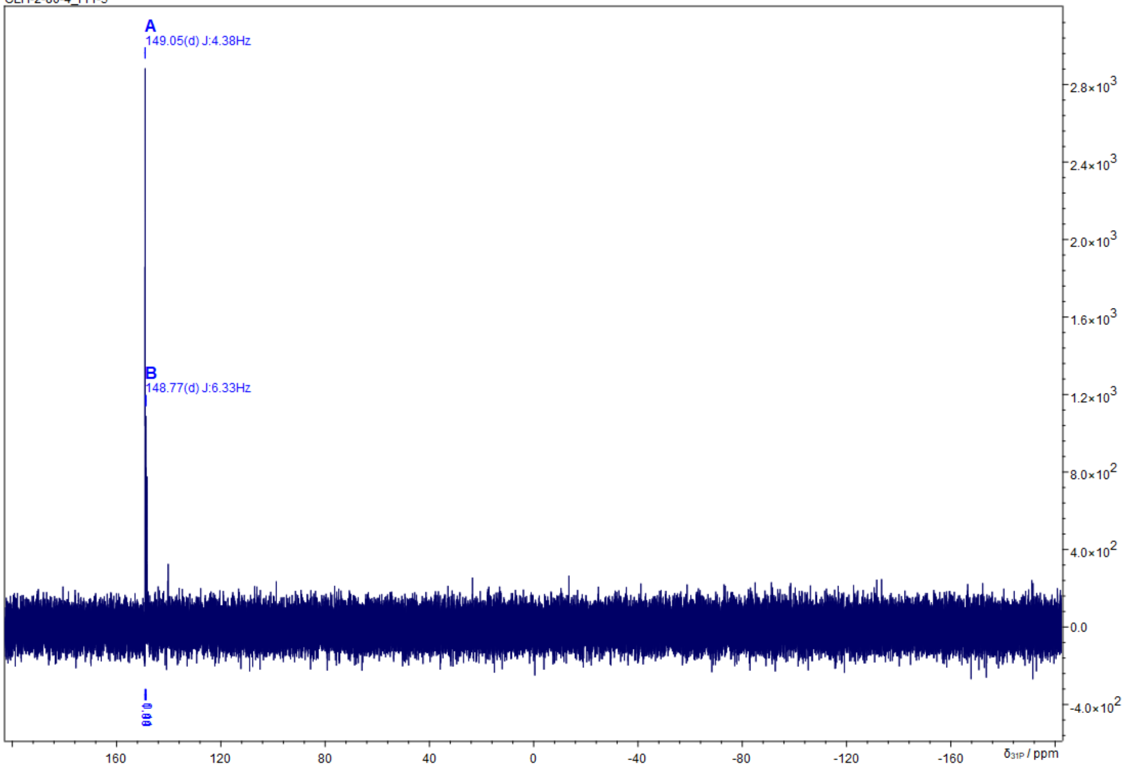

**Benzyl (((4R)-2-phenyl-1,3-dioxolan-4-yl)methyl) ((2-phenyl-1,3-dioxolan-4-yl-4,5-$^{13}C_2$)-methyl-$^{13}C$) phosphate (6)**

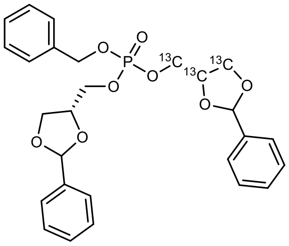

In an argon flushed Schlenk flask 185 mg (0.443 mmol, 1.0 eq.) **5** and 81 mg (0.452 mmol, 1.02 eq.) **4** were azeotropically dried by addition and evaporation using dry toluene, followed by the addition of 4 Å molecular sieves. The dried reagents were dissolved in 8 mL dry DCM, followed by the addition of a solution of 52 mg (0.452 mmol, 1.02 eq.) 4,5-dicyanoimidazole in 2 mL dry ACN. After stirring at RT for 5 h, the reaction was cooled to -78 °C using an acetone/dry-ice bath and 114 mg (0.664 mmol, 1.5 eq.) m-CPBA were added. After 5 h of stirring at -78 °C, the reaction was quenched by the addition of 50 mL sat. $NaHCO_3$. After extraction with DCM (3 x 15 mL), the combined organic phases were washed with 30 mL brine, then dried over $MgSO_4$ and solvent evaporated under reduced pressure to obtain an orange emulsion, which was purified via column chromatography (20 mL $SiO_2$, CH/EE = 1:1 to 2.1, CAM, Rf = 0.22) to obtain 67 mg (30% yield) of the desired product.

| | |
|---|---|
| **$^{13}C_3C_{24}H_{29}O_8P$** | [512.49 g/mol] |
| **Yield** | 67 mg, (0.189 mmol, 30%), yellow oil |
| **TLC** | $R_f$ = 0.22 (CH/EE = 2:1, UV and CAM) |

**$^1$H-NMR** (400 MHz, $CDCl_3$) δ = 7.31 - 7.50 (m, 15H), 5.74 - 5.98 (m, 2H), 5.02 - 5.17 (m, 2H), 3.44 - 4.69 (m, 10H)
**$^{31}$P-NMR** (162 MHz, $CDCl_3$) δ = -0.31 (m)

Z:/Jeol-400/Backup_data/akbref/CLH_2_92_PROTON-1-1.jdf
CLH_2_92

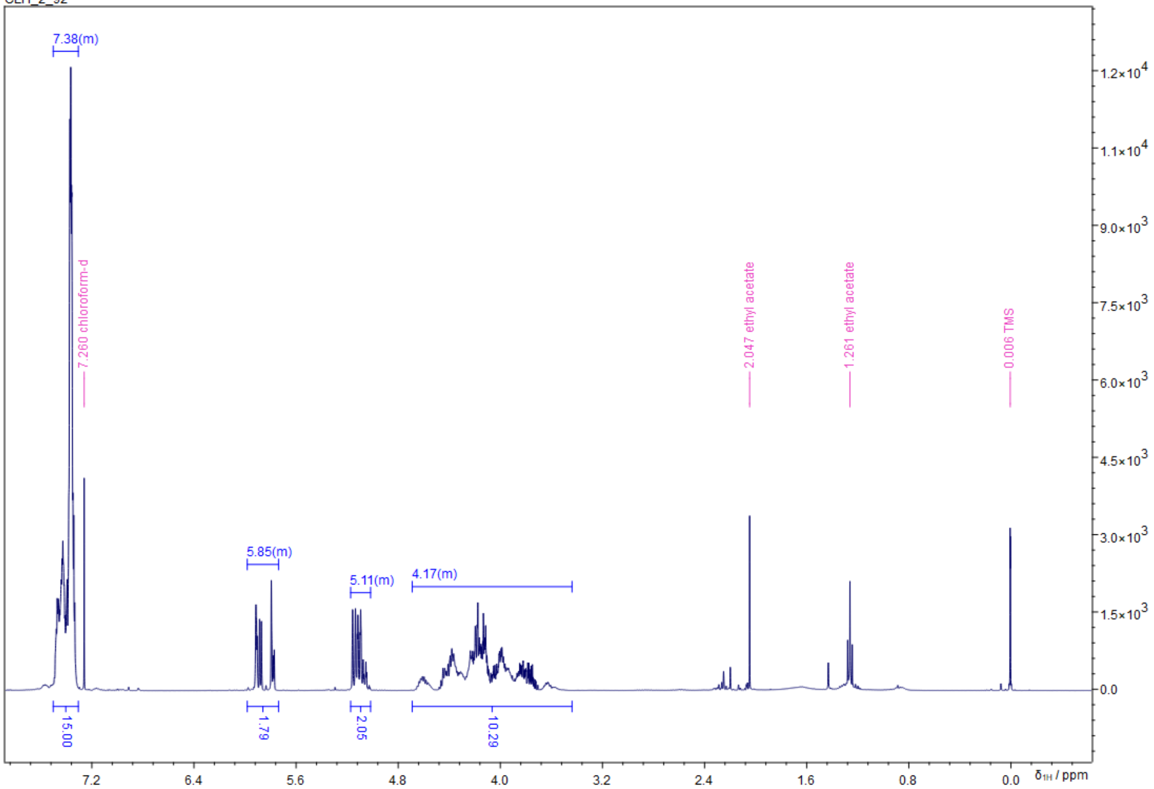


Z:/Jeol-400/Backup_data/akbref/CLH_2_92_PHOSPHORUS-1-1.jdf
CLH_2_92

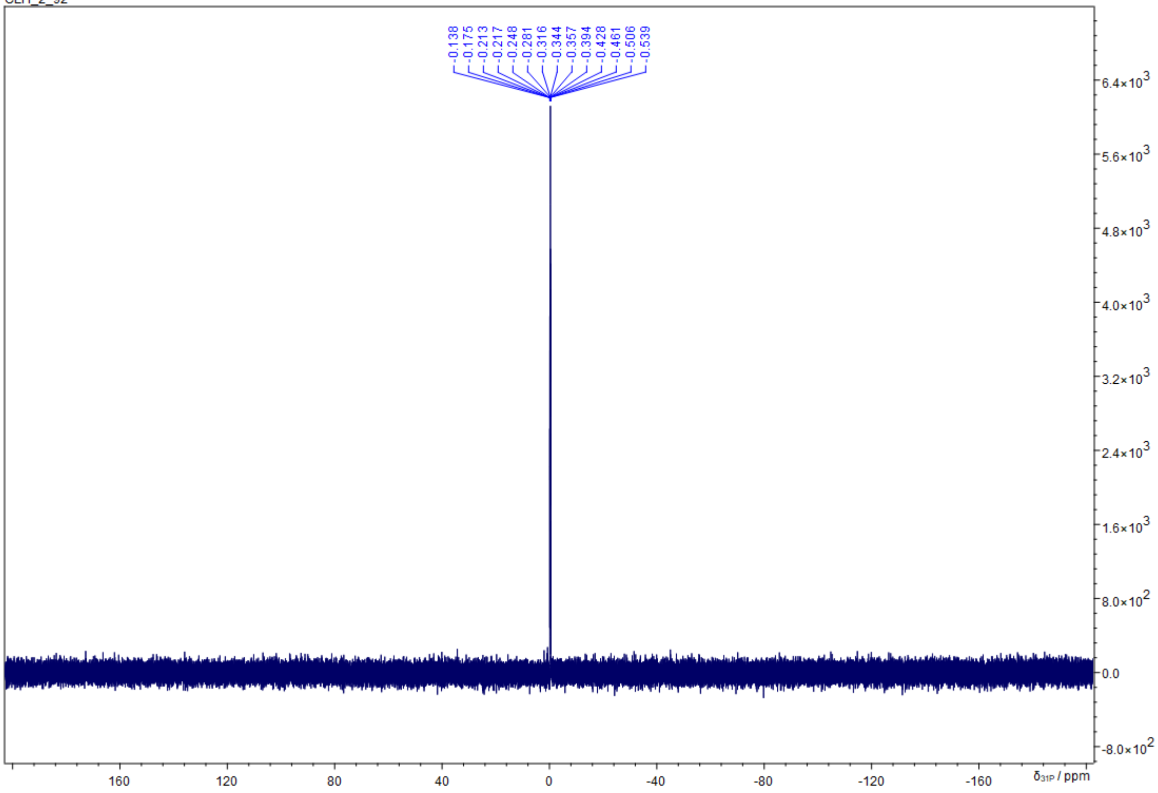

**Sodium ((R)-2,3-dihydroxypropyl) (rac-2,3-dihydroxy-propyl-1,2,3-13$C_3$) phosphate**

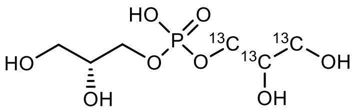

In an argon flushed 10 mL round-bottom flask with a Schlenk adapter 64.5 mg (0.125 mmol, 1.0 eq.) **7** were dissolved in 1.5 mL dry MeOH, followed by the addition of 9 mg (20% $Pd(OH)_2$, 0.013 mmol, 0.1 eq.) $Pd(OH)_2/C$. The flask was carefully evacuated and flushed with $H_2$-gas (3 x), then stirred at RT for 17 h until HPLC-MS indicated full conversion of starting material. The reaction was subsequently diluted with 5 mL MeOH and the Pd-catalyst removed by filtration through a pad of celite (1 cm) under argon. The pad was washed using 5 mL MeOH. The solvent was removed under reduced pressure and the crude product was purified via reverse-phase column chromatography (18C-$SiO_2$, 20 mL, $H_2O$, CAM), obtaining 28 mg of free acid $^{13}C_3$-GPG which was subsequently converted to the sodium salt by addition of 111 µL (0.111 mmol, 1 eq.) 1 M NaOH in 5 mL $H_2O$. After lyophilization, 30.9 mg of the desired product was obtained as an amorphous solid. The final product was contaminated with 12% $^{13}C_3$-glycerol phosphate.

| | |
|---|---|
| **$^{13}C_5C_1H_{15}O_8P$** | [289.05 g/mol] |
| **Yield** | 30.9 mg, (0.111 mmol, 91%), amorphous solid |
| **HPLC-MS** | $t_R$ = 0.560 min (m/z (%): [M+H$^+$] = 250 (58), [M+Na$^+$] = 272 (37), [$M_2$+Na$^+$] = 521 (100) |

**$^1$H-NMR** (400 MHz, $CDCl_3$) δ = 3.97 - 4.16 (m, 1.5H), 3.82 - 3.97 (m, 3H), 3.64 - 3.82 (m, 3H), 3.53 - 3.64 (m, 1.5H), 3.33 - 3.53 (m, 1H)
**$^{13}$C-NMR** (101 MHz, $CDCl_3$) δ = 70.7 (ddd, *J* = 7.8, 41.2, 42.3, **$^{13}$C**), 66.3 (ddd, *J* = 3.1, 5.8, 42.2, **$^{13}$C**), 62.1 (dd, *J* = 3.1, 41.1, **$^{13}$C**)
**$^{31}$P-NMR** (162 MHz, $CDCl_3$) δ = 7.79 (dd, *J* = 7.8, 5.4 Hz)

Z:/Jeol-400/Backup_data/akbref/CLH-2-96_CARBON-1-1.jdf
CLH-2-96

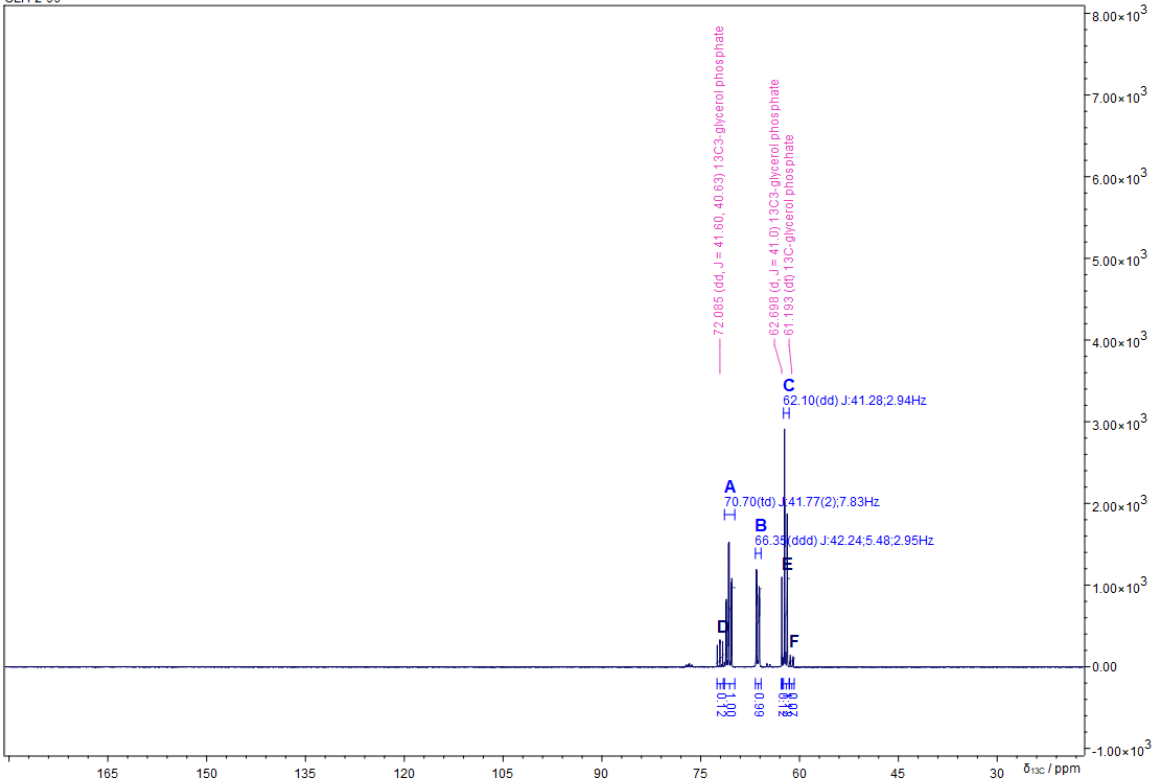


Z:/Jeol-400/Backup_data/akbref/CLH-2-96_PROTON-1-1.jdf
CLH-2-96

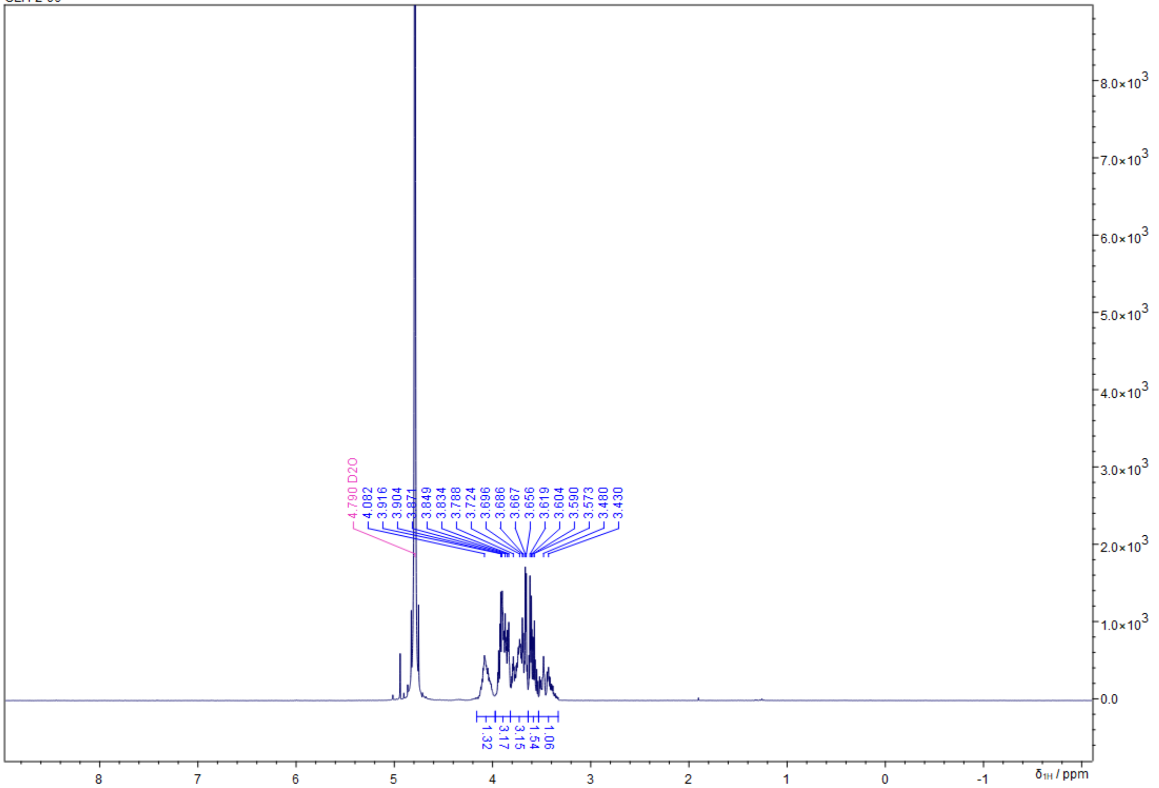

Z:/Jeol-400/Backup_data/akbref/CLH-2-96_PHOSPHORUS-1-1.jdf
CLH-2-96

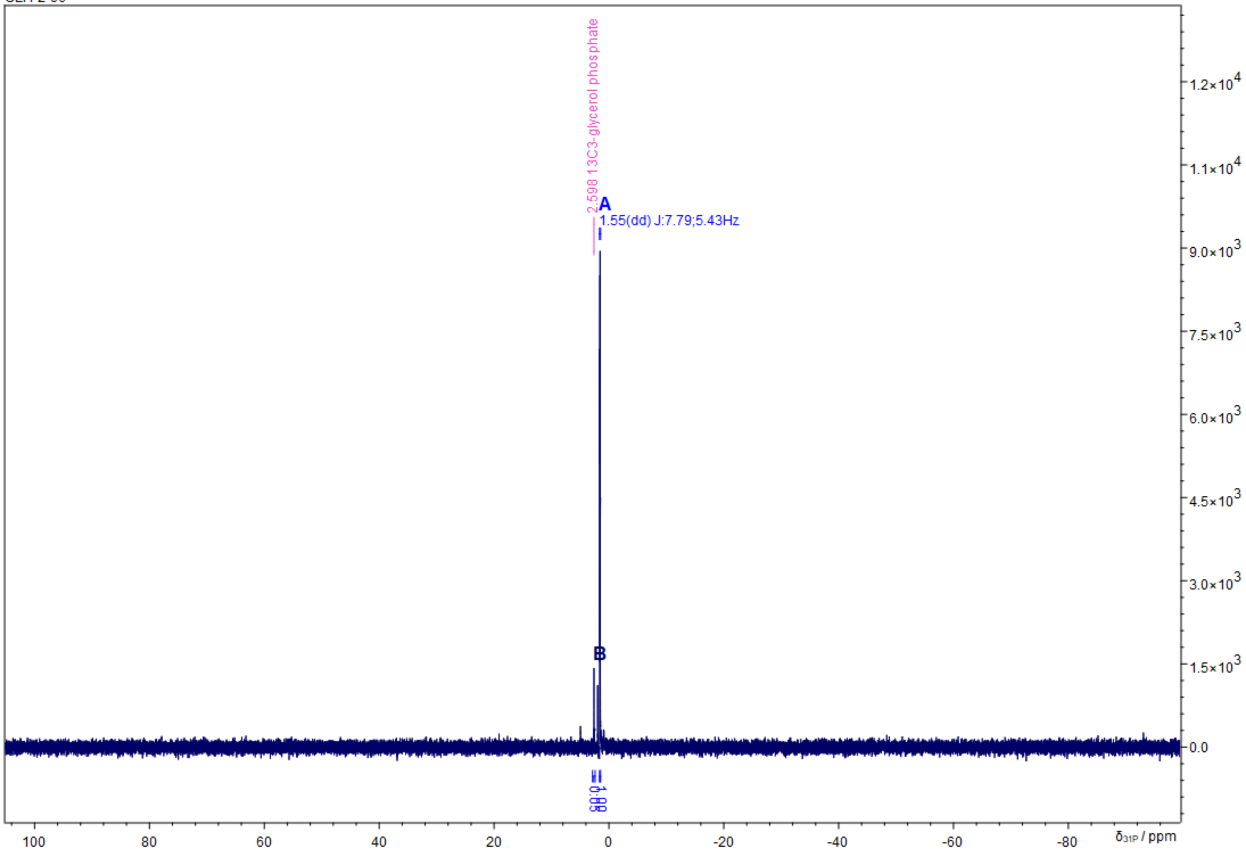

### Synthesis Pathway B:

#### 4-(((4-Methoxybenzyl)oxy)methyl)-2,2-dimethyl-1,3-dioxolane (8)

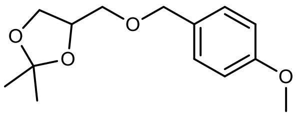

In a 100 mL round-bottom flask 2.51 g (27.1 mmol, 1.0 eq.) glycerol were suspended in 50 mL dry acetone (39.4 g, 678 mmol, 25 eq.), followed by the addition of 10 µL (14.8 mg, 0.19 mmol, 0.01 eq.) methanesulfonic acid. The cloudy mixture was stirred at RT for 40 min yielding a clear colorless solution, which was subsequently neutralized by the addition of solid $K_2CO_3$. After stirring for further 30 min at RT solids were filtered off and solvent was removed under reduced pressure to obtain the crude solketal as a clear colorless oil, which was immediately used in the next step without further purification. The 100 mL round bottom flask containing the intermediate was equipped with a Schlenk adapter and evacuated and then backfilled with Ar (3x). Then the intermediate was dissolved in a mixture of 25 mL THF and 50 mL $Et_2O$. The reaction mixture was cooled to 0 °C in an ice-bath, followed by batchwise addition of 1.41 g (60% in mineral oil, 35.2 mmol, 1.3 eq.) NaH. A bubbler was connected to the Schlenk adapter and the suspension was stirred at 0 °C for 1 h until $H_2$-evolution stopped. Subsequently 5.1 mL (5.86 g, 37.6 mmol, 1.2 eq.) PMB-Cl were added and the ice-bath was removed. The reaction mixture was stirred for 4 d at RT until full conversion of solketal was indicated via GC-MS and excess NaH was quenched by slow addition of 50 mL $H_2O$. The mixture was extracted with 2 x 30 mL EtOAc and the combined organic phases were washed with 25 mL brine and dried over $Na_2SO_4$. The volatiles were removed under reduced pressure to yield the crude product, which was purified via column chromatography (250 mL $SiO_2$, CH/EE = 9:1, CAM, Rf = 0.20) yielding 2.68 g (39% yield) of a colorless oil.

| | |
|---|---|
| **$C_{14}H_{20}O_4$** | [252.3 g/mol] |
| **Yield** | 2.68 g (10.6 mmol, 39%), colorless oil |
| **TLC** | $R_f$ = 0.20 (CH/EtOAc = 9:1, UV and CAM) |
| **GC-MS** | $t_R$ = 6.503 min; m/z (%): 252 (2), 194 (13), 163 (12), 121 (100) |

**$^1$H-NMR** (400 MHz, $CDCl_3$) δ = 7.26 (d, *J* = 8.7, 2H, H12, H16), 6.88 (d, *J* = 8.7, 2H, H13, H15), 4.45 - 4.56 (m, 2H, H10), 4.28 (p, *J* = 6.0, 1H), 4.04 (dd, *J* = 6.4, 8.3, 1H), 3.72 (dd, *J* = 6.3, 8.3, 1H), 3.52 (dd, *J* = 5.7, 9.8, 1H), 3.49 (ddd, *J* = 5.6, 9.8, 35.4, 2H, H2), 1.42 (s, 3H, H8), 1.36 (s, 3H, H9)
**$^{13}$C-NMR** (101 MHz, $CDCl_3$) δ = 159.4 (s), 130.1 (s), 129.5 (s), 113.9 (s), 109.5 (s), 74.8 (s), 73.3 (s), 70.9 (s), 67.0 (s), 55.4 (s), 27.0 (s), 26.9 (d, *J* = 12.8), 25. 5 (s).

Z:/Jeol-400/Backup_data/akbref/CLH-3-144_PROTON-1-1.jdf
CLH-3-144

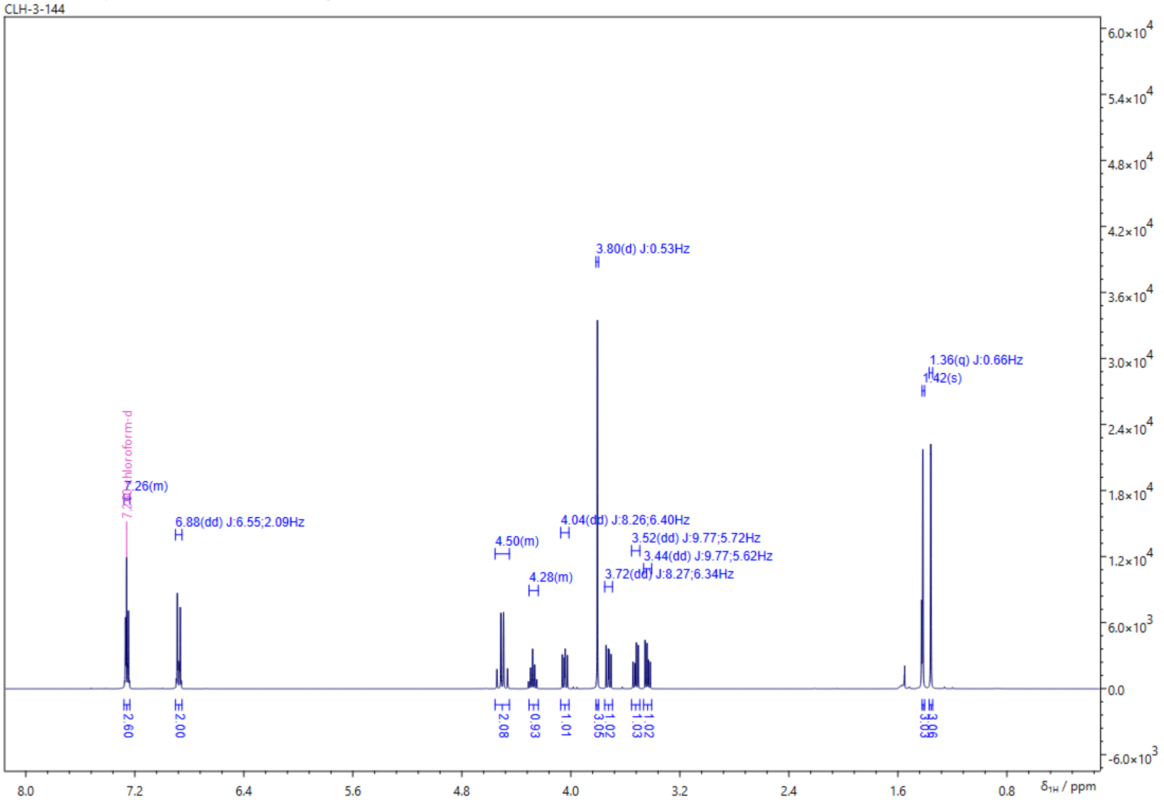


Z:/Jeol-400/Backup_data/akbref/CLH-3-144_CARBON-1-1.jdf
CLH-3-144

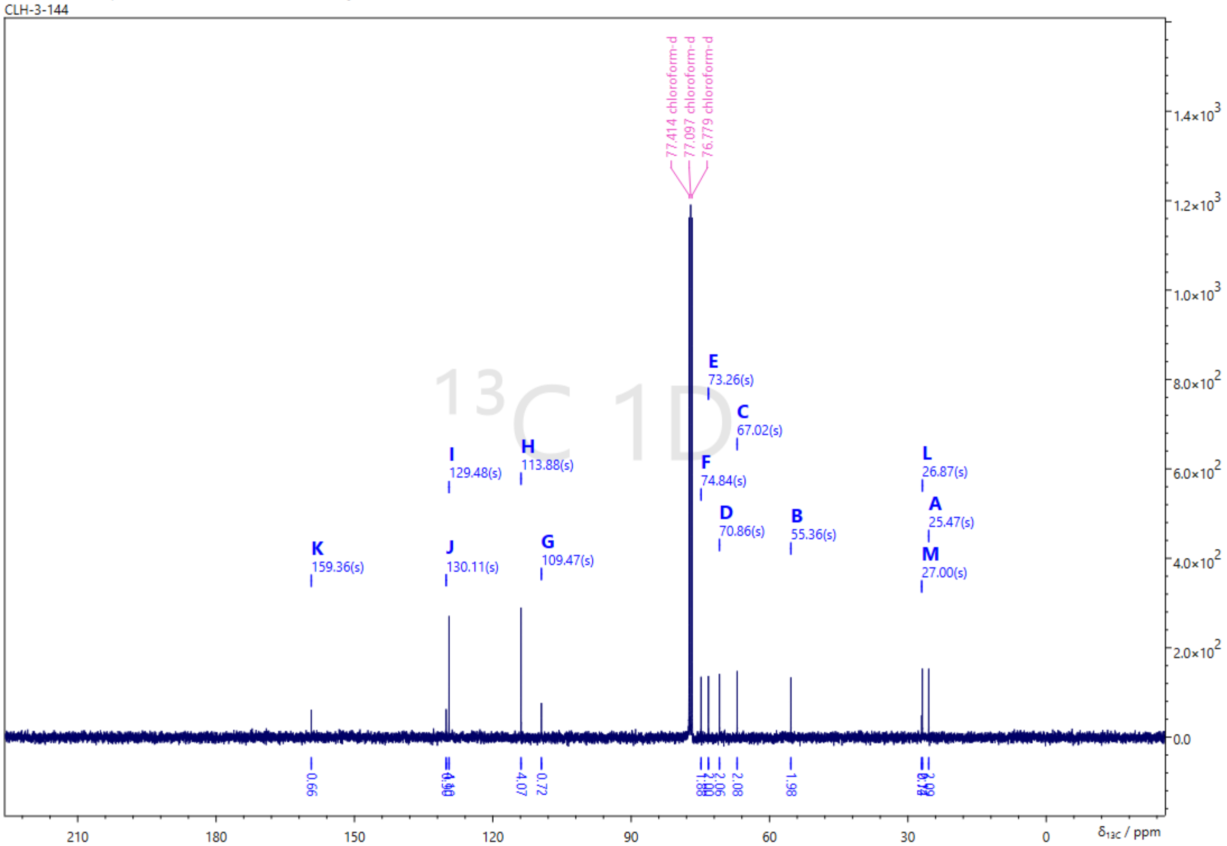

**4-(((4-Methoxybenzyl)oxy)methyl-$^{13}$C)-2,2-dimethyl-1,3-dioxolane-5-$^{13}$C (9)**

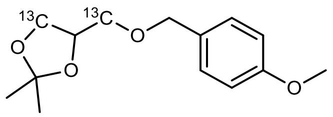

In a 50 mL round-bottom flask 255.9 mg (2.72 mmol, 1.0 eq.) $^{13}C_2$-glycerol were suspended in 6 mL dry acetone (4.74 g, 81.6 mmol, 30 eq.), followed by the addition of 1.3 µL (1.92 mg, 0.02 mmol, 0.01 eq.) methanesulfonic acid. The cloudy mixture was stirred at RT for 18 h yielding a clear colorless solution, which was subsequently neutralized by the addition of solid $K_2CO_3$. After stirring for further 30 min at RT solids were filtered off and the solvent was evaporated under reduced pressure to obtain crude $^{13}C_2$-solketal as a clear colorless oil, which was immediately used in the next step without further purification. The 50 mL round bottom flask containing the intermediate was equipped with a Schlenk adapter and evacuated and then backfilled with Ar (3x). Then the intermediate was dissolved in a mixture of 25 mL THF. The reaction mixture was cooled to 0 °C in an ice-bath, followed by addition of 141 mg (60% in mineral oil, 3.52 mmol, 1.3 eq.) NaH. A bubbler was connected to the Schlenk adapter and the suspension was stirred at 0 °C for 25 min until $H_2$-evolution stopped, which was accompanied by strong thickening of the solution. Subsequently, a solution of 498 µL (572 mg, 4.05 mmol, 1.2 eq.) PMB-Cl in 12.5 mL $Et_2O$ was added over the course of 10 min and the ice-bath was removed. The reaction mixture was stirred for 4 d at RT until full conversion of $^{13}C_2$-solketal was indicated via GC-MS and excess NaH was quenched by slow addition of 150 mL $H_2O$. The mixture was extracted with 2 x 100 mL EtOAc and the combined organic phases were washed with 50 mL brine and dried over $Na_2SO_4$. The volatiles were removed under reduced pressure to yield a yellow oil, which was further purified via column chromatography (50 mL $SiO_2$, CH/EE = 10:1, CAM) yielding 209 mg (30% yield) of a colorless oil.

| | |
|---|---|
| **$^{13}C_2C_{10}H_{20}O_4$** | [254.3 g/mol] |
| **Yield** | 209 mg (0.82 mmol, 30%), colorless oil |
| **TLC** | $R_f$ = 0.19 (CH/EtOAc = 10:1, UV and CAM) |
| **GC-MS** | $t_R$ = 6.487 min; m/z (%): 254 (2), 196 (13), 165 (12), 121 (100) |

**$^1$H-NMR** (400 MHz, $CDCl_3$) δ = 7.26 (d, *J* = 8.1, 2H), 6.88 (d, *J* = 8.1, 2H), 4.43 - 4.59 (m, 2H), 4.18 - 4.33 (m, 1H), 3.75 - 3.95 (m, 4H), 3.21 - 3.75 (m, 2H), 1.42 (s, 2H), 1.36 (s, 2H).
**$^{13}$C-NMR** (101 MHz, $CDCl_3$) δ = 159.4 (s), 130.2 (s), 129.5 (s), 113.9 (s), 109.5 (s), 74.9 (dd, *J* = 33.8, 44.9, 1C), 73.3 (s), 70.9 (s), 67.1 (s), 55.4 (s), 26.9 (s), 25.5 (s).

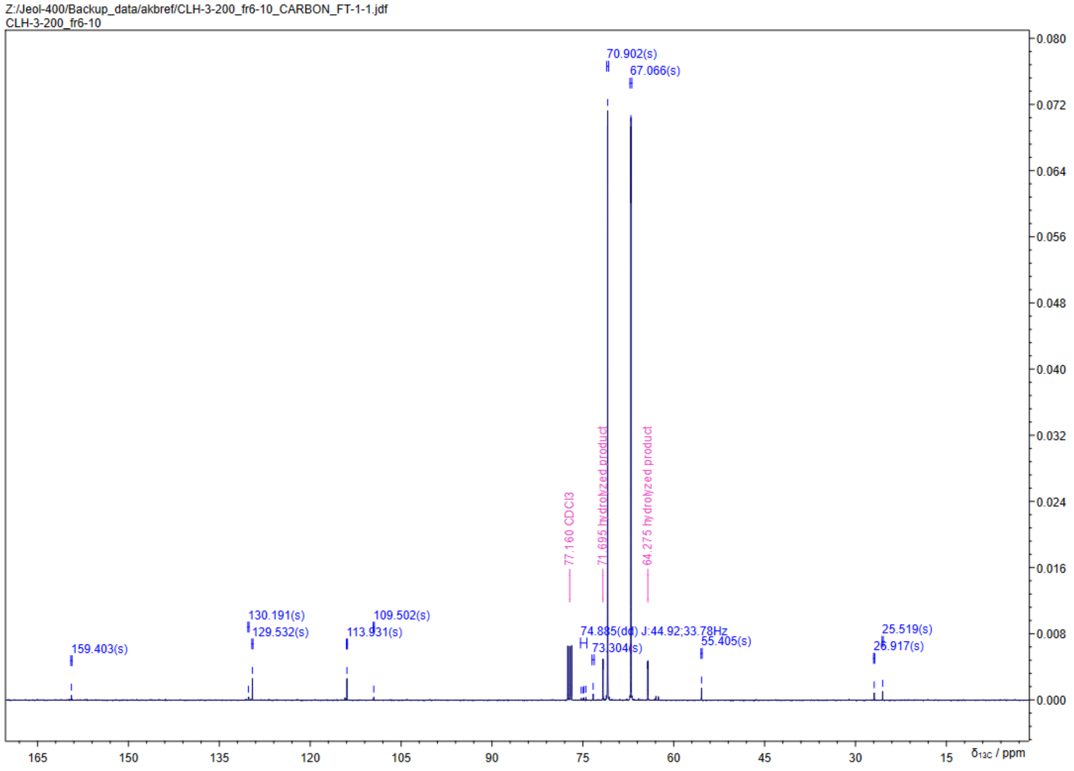
Z:/Jeol-400/Backup_data/akbref/CLH-3-200_fr6-10_CARBON_FT-1-1.jdf
CLH-3-200_fr6-10
70.902(s)
67.066(s)
77.160 CDCl3
71.695 hydrolyzed product
64.275 hydrolyzed product
130.191(s)
129.532(s)
109.502(s)
113.931(s)
159.403(s)
74.885(dd) J:44.92;33.78Hz
73.304(s)
55.405(s)
25.519(s)
26.917(s)
δ13C / ppm

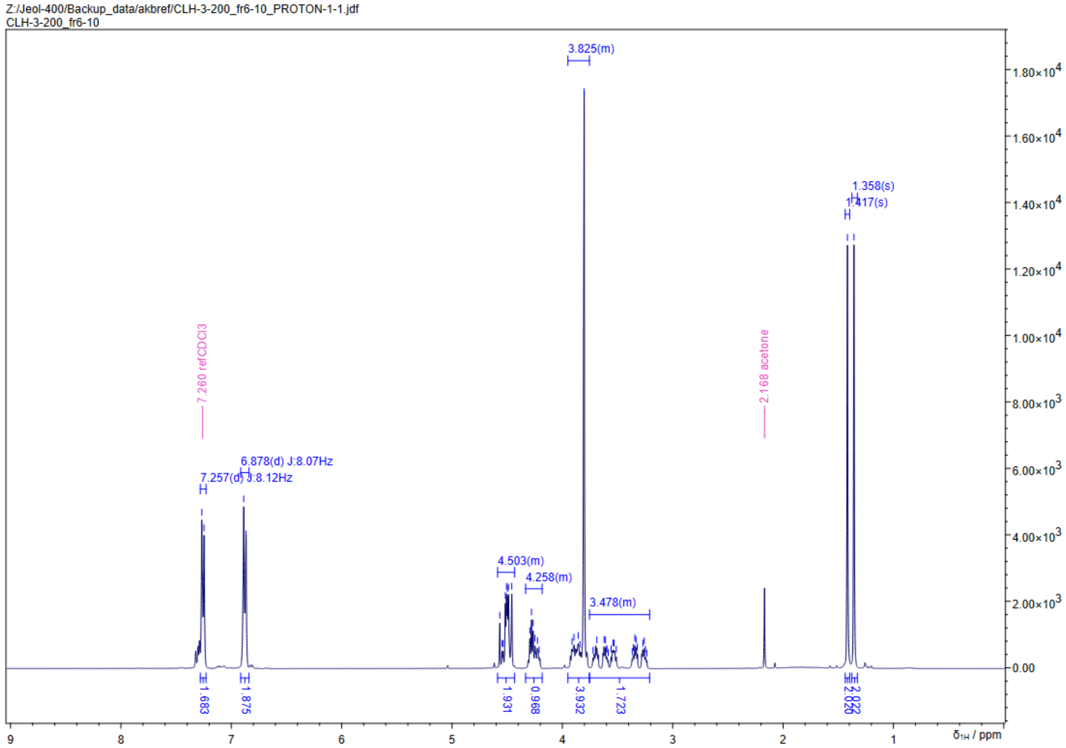
Z:/Jeol-400/Backup_data/akbref/CLH-3-200_fr6-10_PROTON-1-1.jdf
CLH-3-200_fr6-10
3.825(m)
1.358(s)
1.417(s)
7.260 ref CDCl3
2.168 acetone
6.878(d) J:8.07Hz
7.257(d) J:8.12Hz
4.503(m)
4.258(m)
3.478(m)
1.683
1.875
1.931
0.968
3.932
1.723
δ1H / ppm

**4-(((4-Methoxybenzyl)oxy)methyl-$^{13}$C)-2,2-dimethyl-1,3-dioxolane-4,5-$^{13}C_2$ (10)**

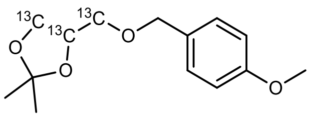

In a 50 mL round-bottom flask 406 mg (4.27 mmol, 1.0 eq.) $^{13}C_3$-glycerol were suspended in 8 mL dry acetone (6.31 g, 108.7 mmol, 25.5 eq.), followed by the addition of 2.7 µL (4.1 mg, 0.024 mmol, 0.01 eq.) methanesulfonic acid. The cloudy mixture was stirred at RT for 90 min yielding a clear colorless solution, which was subsequently neutralized by the addition of solid $K_2CO_3$. After stirring for further 30 min at RT the solids were filtered off and the solvent was removed under reduced pressure to obtain 526 mg (3.88 mmol, 91%) crude $^{13}C_3$-solketal as a clear colorless oil, which was immediately used in the next step. The 50 mL round bottom flask containing the intermediate was equipped with a Schlenk adapter and evacuated and then backfilled with Ar (3x). Then the intermediate was dissolved in a mixture of 10 mL THF and 20 mL $Et_2O$. The reaction mixture was cooled to 0 °C in an ice-bath, followed by addition of 225 mg (60% in mineral oil, 5.55 mmol, 1.3 eq.) NaH. A bubbler was connected to the Schlenk adapter and the suspension was stirred at 0 °C for 15 min until $H_2$-evolution stopped. Subsequently, 700 µL (805 mg, 5.14 mmol, 1.2 eq.) PMB-Cl were added over the course of 10 min and the ice-bath was removed. The reaction mixture was stirred for 4 d at RT until full conversion of $^{13}C_3$-solketal was indicated via GC-MS and excess NaH was quenched by slow addition of 30 mL $H_2O$. The mixture was extracted with 2 x 30 mL EtOAc and the combined organic phases were dried over $Na_2SO_4$. The solvents were removed under reduced pressure to yield a yellow oil, which was further purified via column chromatography (100 mL $SiO_2$, CH/EE = 10:1, CAM, Rf = 0.17) yielding 564 mg (51% yield) of a pale yellow oil.

| | |
|---|---|
| **$^{13}C_3C_9H_{20}O_4$** | [255.3 g/mol] |
| **Yield** | 564 mg (2.20 mmol, 51%), pale yellow oil |
| **TLC** | $R_f$ = 0.17 (CH/EtOAc = 10:1, UV and CAM) |
| **GC-MS** | $t_R$ = 6.505 min; 255 (3), 197 (12), 166 (12), 121 (100) |

**$^1$H-NMR** (400 MHz, $CDCl_3$) δ = 7.26 (d, *J* = 8.7, 2H), 6.88 (d, *J* = 8.6, 2H), 4.42 - 4.58 (m, 3H), 4.19 - 4.27 (m, 0H), 4.09 ([qi], *J* = 6.1, 0H), 3.83 - 3.94 (m, 1H), 3.80 (s, 3H), 3.58 - 3.74 (m, 1H), 3.50 - 3.57 (m, 1H), 3.22 - 3.39 (m, 1H), 1.42 (s, 3H), 1.36 (s, 3H).
**$^{13}$C-NMR** (101 MHz, $CDCl_3$) δ = 159.4 (s), 130.2 (s), 129.5 (s), 113.9 (s), 109.5 (s), 74.9 (dd, *J* = 33.9, 44.8, **$^{13}$C**), 73.3 (s), 70.9 (dd, *J* = 1.4, 44.8, **$^{13}$C**), 67.1 (dd, *J* = 1.4, 33.9, **$^{13}$C**), 55.4 (s), 26.9 (s), 25.5 (s).

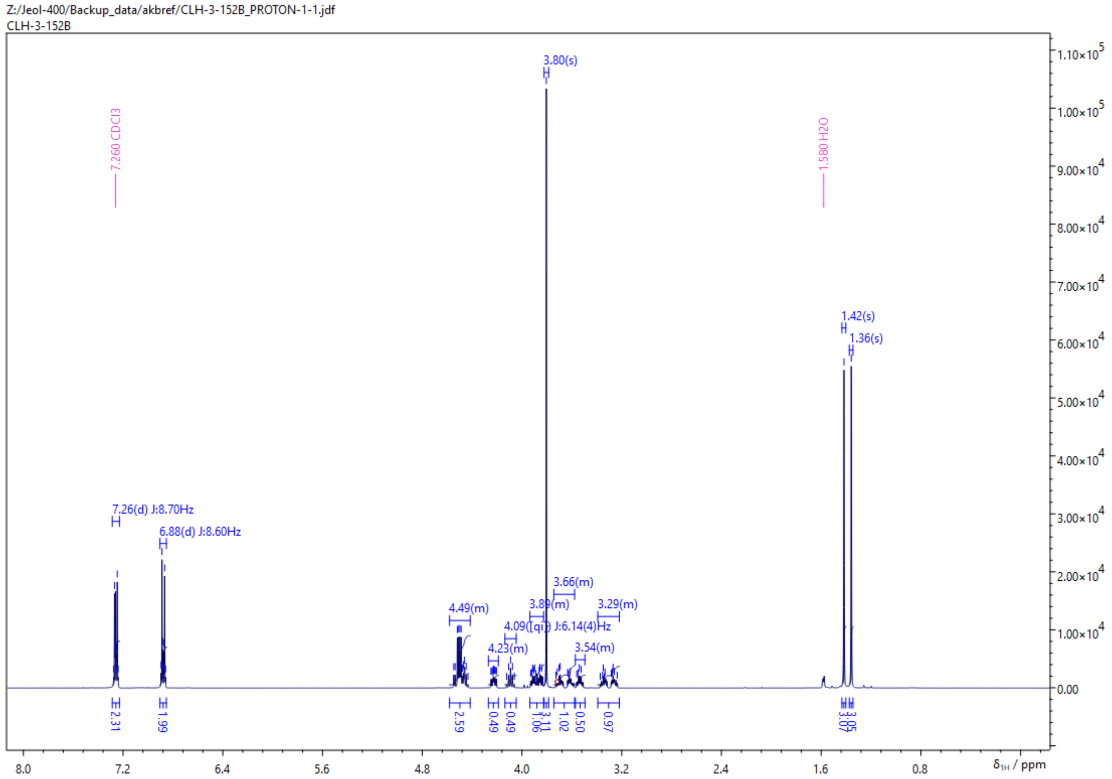

Z:/Jeol-400/Backup_data/akbref/CLH-3-152B_PROTON-1-1.jdf
CLH-3-152B
7.260 CDCl3
1.580 H2O
3.80(s)
1.42(s)
1.36(s)
7.26(d) J:8.70Hz
6.88(d) J:8.60Hz
4.49(m)
4.23(m)
4.09(qi) J:6.14(4)Hz
3.89(m)
3.66(m)
3.54(m)
3.29(m)
δ1H / ppm


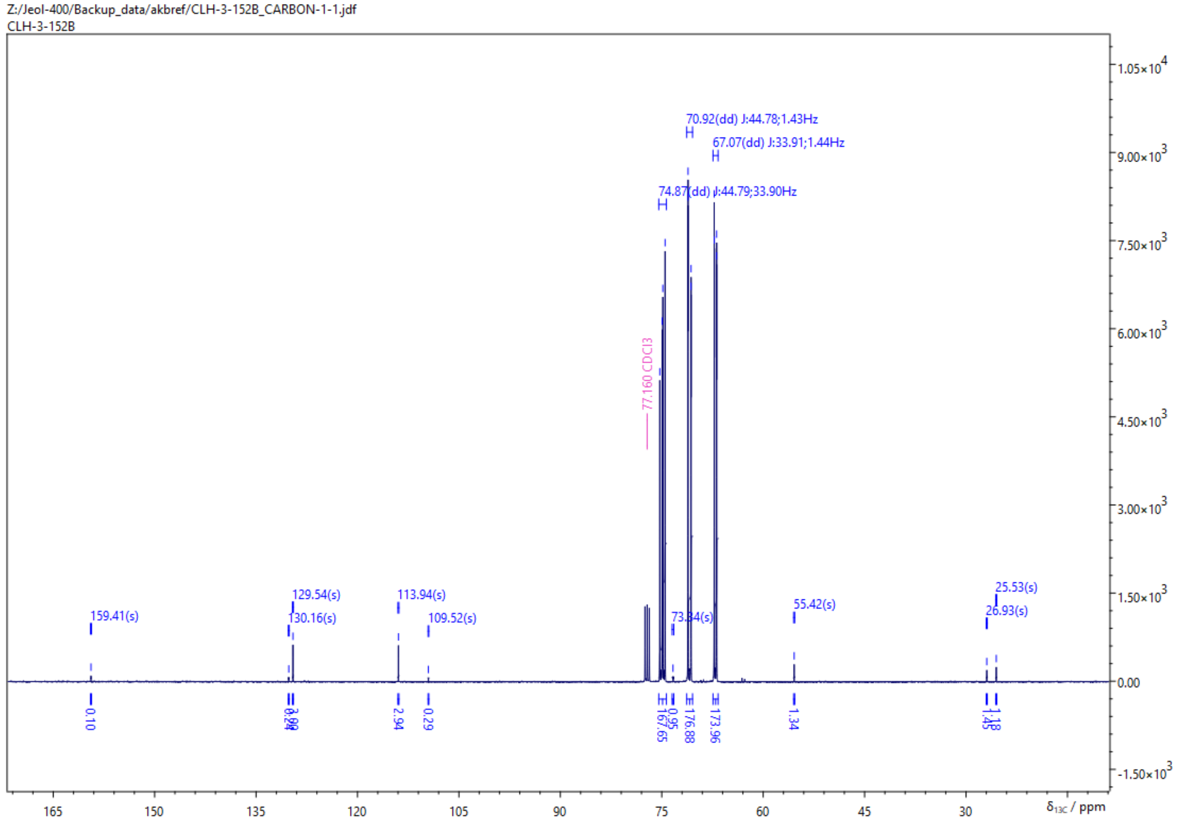

Z:/Jeol-400/Backup_data/akbref/CLH-3-152B_CARBON-1-1.jdf
CLH-3-152B
70.92(dd) J:44.78;1.43Hz
67.07(dd) J:33.91;1.44Hz
74.87(dd) J:44.79;33.90Hz
77.160 CDCl3
159.41(s)
129.54(s)
130.16(s)
113.94(s)
109.52(s)
73.84(s)
55.42(s)
25.53(s)
26.93(s)
δ13C / ppm

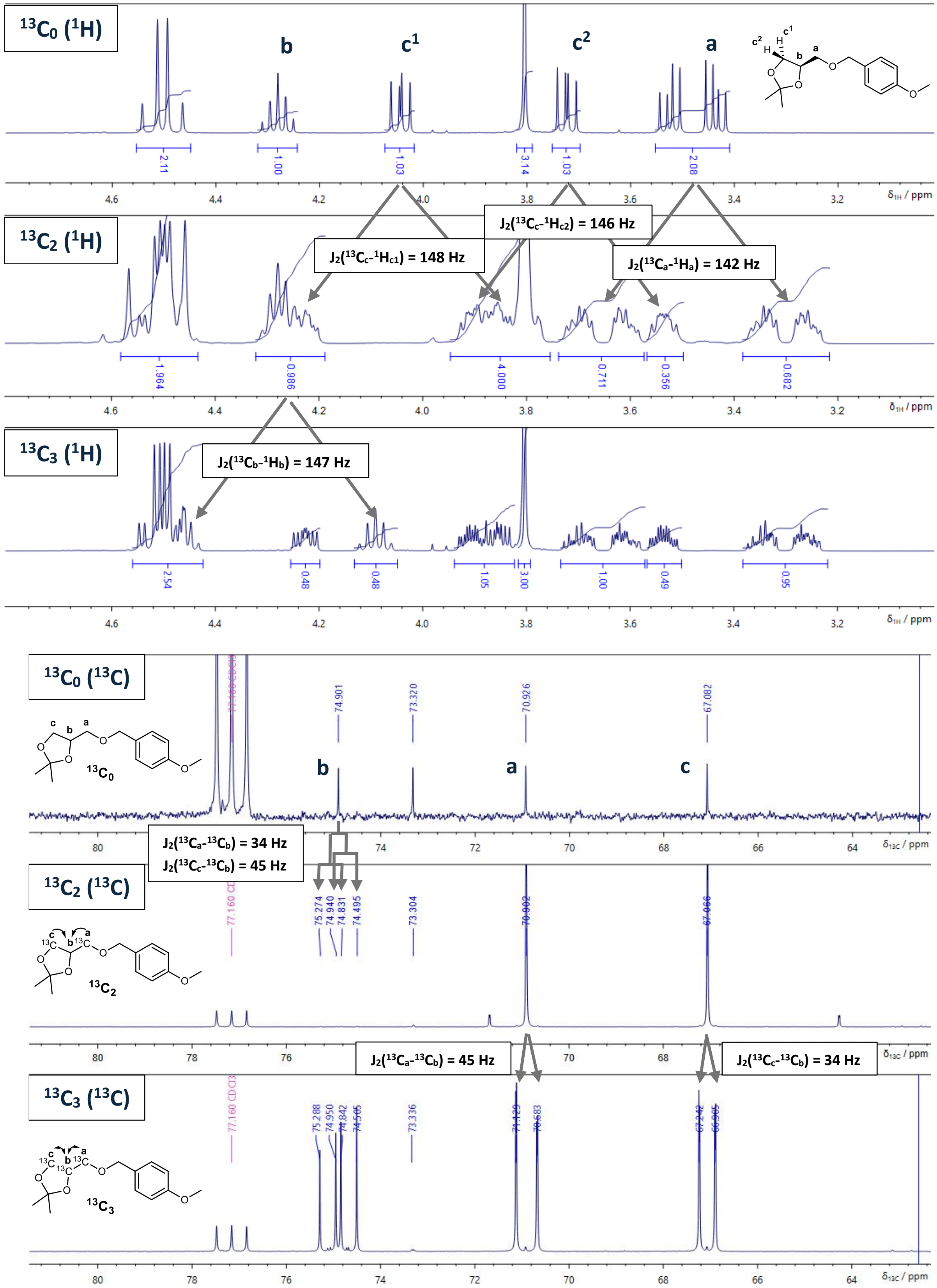

13C0 (1H)
b
c1
c2
a
13C2 (1H)
J2(13Cc-1Hc1) = 148 Hz
J2(13Cc-1Hc2) = 146 Hz
J2(13Ca-1Ha) = 142 Hz
13C3 (1H)
J2(13Cb-1Hb) = 147 Hz
δ1H / ppm
13C0 (13C)
13C0
74.901
73.320
70.926
67.082
b
a
c
J2(13Ca-13Cb) = 34 Hz
J2(13Cc-13Cb) = 45 Hz
13C2 (13C)
13C2
77.160 CDCl3
75.274
74.940
74.831
74.495
73.304
J2(13Ca-13Cb) = 45 Hz
J2(13Cc-13Cb) = 34 Hz
13C3 (13C)
13C3
77.160 CDCl3
75.288
74.950
74.842
74.505
73.336
70.683
δ13C / ppm

**(((3-((4-Methoxybenzyl)oxy)propane-1,2-diyl)bis(oxy))bis(methylene))dibenzene (11)**

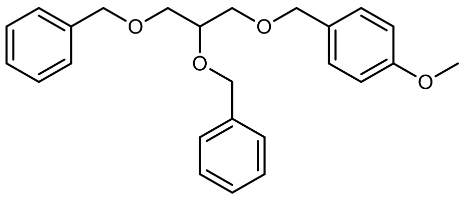

In a 100 mL round-bottom flask 2.34 g (9.31 mmol, 1.0 eq.) **8** were dissolved in 10 mL 90% acetic acid. The reaction was stirred at 40 °C until full conversion was indicated by HPLC (3 h). The volatiles were removed under reduced pressure to yield a pale yellow oil, which was used immediately in the next step without further purification. The 100 mL round bottom flask containing the intermediate was equipped with a Schlenk adapter and evacuated and then backfilled with Ar (3x). Then the intermediate was dissolved in a mixture of 40 mL THF and 20 mL $Et_2O$. The reaction mixture was cooled to 0 °C in an ice-bath, followed by addition of 969 mg (60% in mineral oil, 24.2 mmol, 2.6 eq.) NaH. A bubbler was connected to the Schlenk adapter and the reaction mixture was stirred for 15 min at 0 °C before 2.6 mL (3.74 g, 21.6 mmol, 2.4 eq.) benzyl bromide were added to the mixture. After removal of the ice bath the reaction mixture was stirred for 18 h. Excess NaH was subsequently quenched by the addition of 100 mL $H_2O$. The reaction mixture was extracted with EtOAc (2 x 50 mL). The combined organic phases were washed with brine (50 mL) and dried over $Na_2SO_4$. The solvents were removed under reduced pressure to yield the crude product as a dark yellow oil. Purification via column chromatography (250 mL $SiO_2$, CH/EE = 15:1, CAM, $R_f$ = 0.13) yielded 1.90 g (52% yield) of a yellowish oil.

**$C_{25}H_{28}O_4$** [392.50 g/mol]

**Yield** 1.90 g (4.89 mmol, 52%), yellowish oil

**TLC** $R_f$ = 0.13 (CH/EtOAc = 15:1, UV and CAM)

**HPLC-MS** $t_R$ = 5.815 min; m/z (%): [$M+H^+$] = 393 (19), [$M+NH_4^+$] = 410 (37), [$M+Na^+$]= 415 (100)

**$^1$H-NMR** (400 MHz, $CDCl_3$) δ = δ = 7.13 - 7.30 (m, 12H), 6.79 (dd, *J* = 2.0, 6.7, 2H), 4.62 (s, 2H), 4.46 (s, 2H), 4.39 (s, 2H), 3.69 - 3.76 (m, 4H), 3.48 - 3.59 (m, 4H).
**$^{13}$C-NMR** (101 MHz, $CDCl_3$) δ = 159.3 (s), 138.8 (s), 138.5 (s), 130.5 (s), 129.4 (s), 128.5 (s), 128.4 (s), 127.9 (s), 127.7 (s), 127.7 (s), 127.6 (s, 2C), 113.9 (s), 77.4 (s), 73.5 (s), 73.2 (s), 72.3 (s), 70.53 (s), 70.1 (s), 55.4 (s), 27.1 (s).

Z:/Jeol-400/Backup_data/akbref/CLH-3-145B_PROTON-1-1.jdf
CLH-3-145B

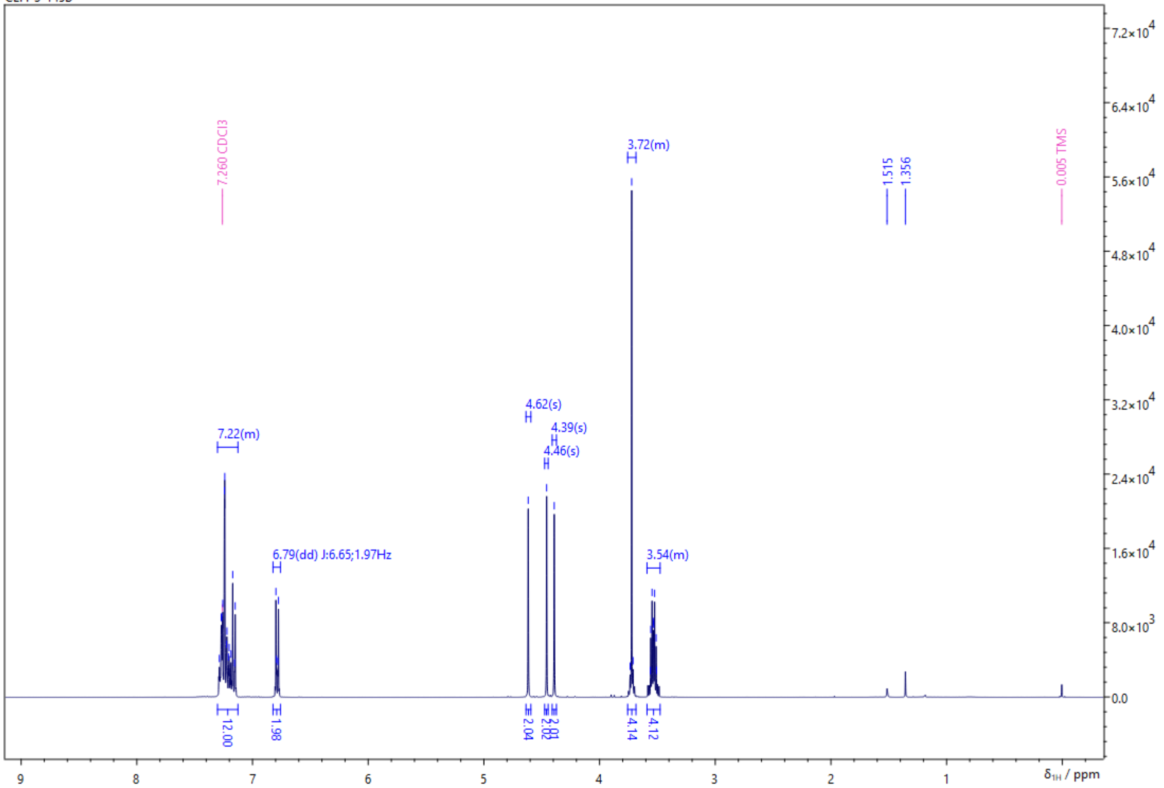


Z:/Jeol-400/Backup_data/akbref/CLH-3-145B_CARBON-1-1.jdf
CLH-3-145B

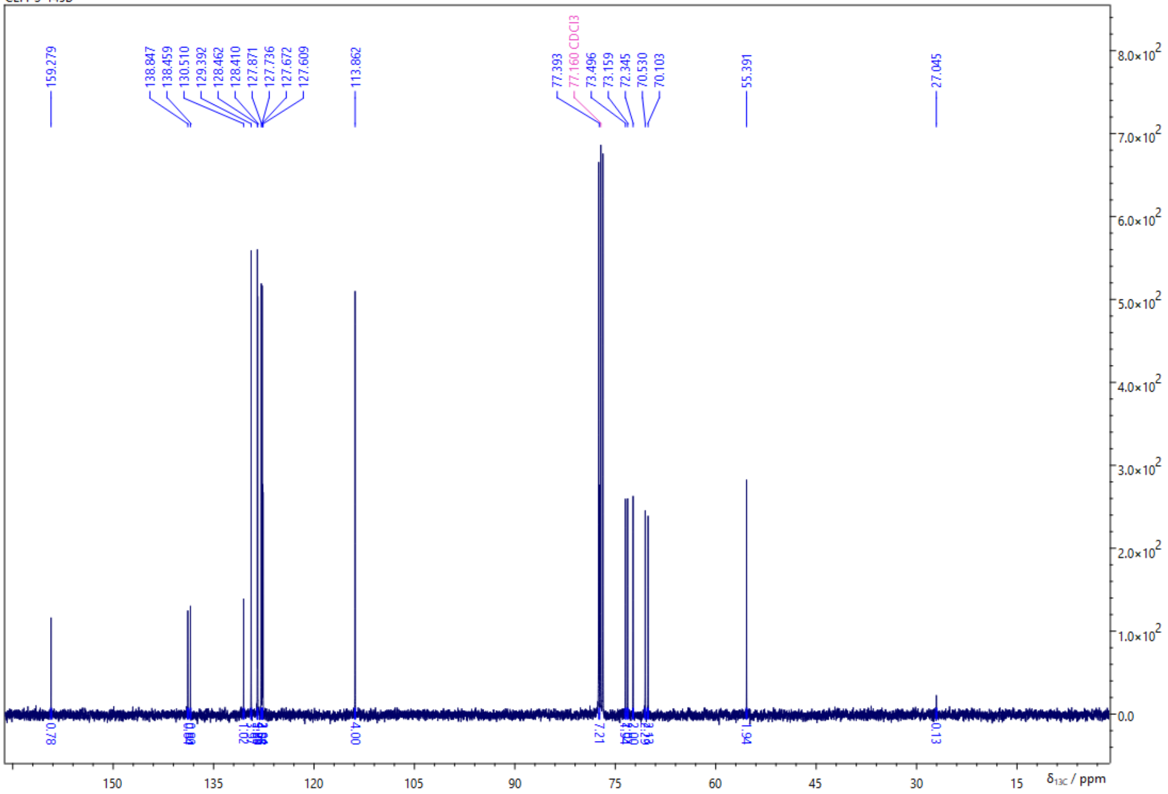

**(((3-((4-Methoxybenzyl)oxy)propane-1,2-diyl-1,3-$^{13}C_2$)bis(oxy))bis(methylene))dibenzene (12)**

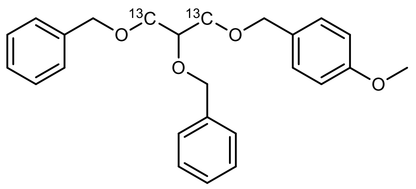

In a 100 mL round-bottom flask 207.3 mg (0.815 mmol, 1.0 eq.) **9** were dissolved in 5 mL 90% acetic acid. The reaction mixture was stirred at 40 °C until full conversion was indicated by HPLC (5 h). The solvent was removed under reduced pressure to yield a colorless oil, which was used immediately in the next step without further purification. The 100 mL round bottom flask containing the intermediate was equipped with a Schlenk adapter and evacuated and then backfilled with Ar (3x). Then the intermediate was dissolved in a mixture of 30 mL THF and 15 mL $Et_2O$. The reaction mixture was cooled to 0 °C in an ice-bath, followed by addition of 85 mg (60% in mineral oil, 3.52 mmol, 2.6 eq.) NaH. A bubbler was connected to the Schlenk adapter and the suspension was stirred at 0 °C for 10 min until $H_2$-evolution stopped, which was accompanied by strong thickening of the solution. Subsequently, a solution of 498 µL (498 mg, 3.52 mmol, 2.4 eq.) PMB-Cl in 12.5 mL $Et_2O$ was added over the course of 10 min and the ice-bath was removed. The reaction was stirred for 4 d at RT until full conversion of $^{13}C_2$-glycerol was indicated via GC-MS and excess NaH was quenched by slow addition of 150 mL $H_2O$. The mixture was extracted with 2 x 100 mL EtOAc and the combined organic phases were washed with 50 mL brine and dried over $Na_2SO_4$. The volatiles were removed under reduced pressure to yield a yellow oil, which was further purified via column chromatography (50 mL $SiO_2$, CH/EE = 15:1, CAM, $R_f$ = 0.12) yielding 209 mg (30% yield) of a colorless oil.

| | |
|---|---|
| **$^{13}C_2C_{23}H_{28}O_4$** | [394.50 g/mol] |
| **Yield** | 209 mg (0.529 mmol, 65%), colorless oil |
| **TLC** | $R_f$ = 0.12 (CH/EtOAc =15:1 , UV and CAM) |
| **HPLC-MS** | $t_R$ = 5.812 min; m/z (%): [M+H$^+$] = 395 (13), [M+NH$_4^+$] = 412 (36), [M+Na$^+$] = 417 (100) |

**$^1$H-NMR** (400 MHz, $CDCl_3$) δ = 7.12 - 7.31 (m, 12H), 6.78 (d, *J* = 8.2, 2H), 4.61 (s, 2H), 4.45 (d, *J* = 3.4, 2H), 4.39 (d, *J* = 3.3, 2H), 3.72 (d, *J* = 10.0, 6H), 3.28 - 3.42 (m, 2H).
**$^{13}$C-NMR** (101 MHz, $CDCl_3$) δ = 159.3 (s), 138.9 (s), 138.5 (s), 129.4 (s), 128.5 (d, *J* = 5.1), 127.5 - 128.1 (m), 113.9 (s), 77.4 (s), 73.2 (s), 72.4 (s, **$^{13}$C**), 70.5 (s, **$^{13}$C**), 70.1 (s, **$^{13}$C**), 55.4 (s).

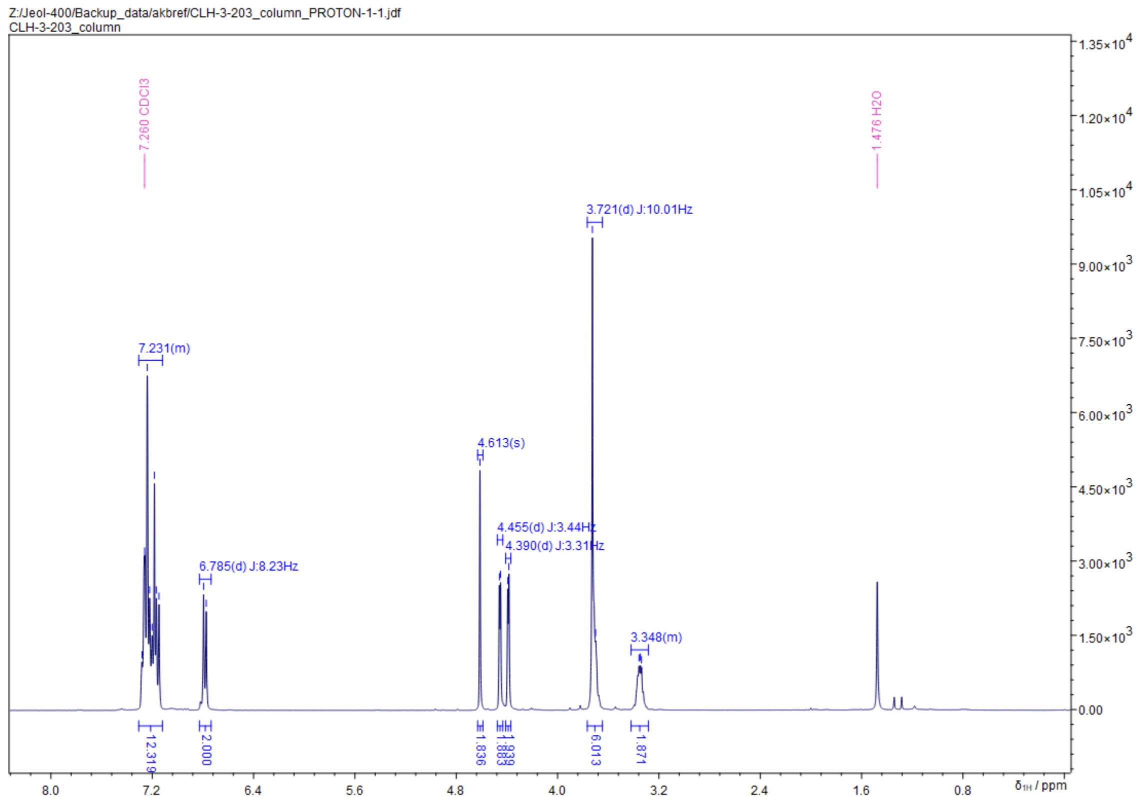

Z:/Jeol-400/Backup_data/akbref/CLH-3-203_column_PROTON-1-1.jdf
CLH-3-203_column
7.260 CDCl3
1.476 H2O
3.721(d) J:10.01Hz
7.231(m)
4.613(s)
4.455(d) J:3.44Hz
4.390(d) J:3.31Hz
6.785(d) J:8.23Hz
3.348(m)
12.319
2.000
1.836
1.883
1.939
6.013
1.871
δ1H / ppm


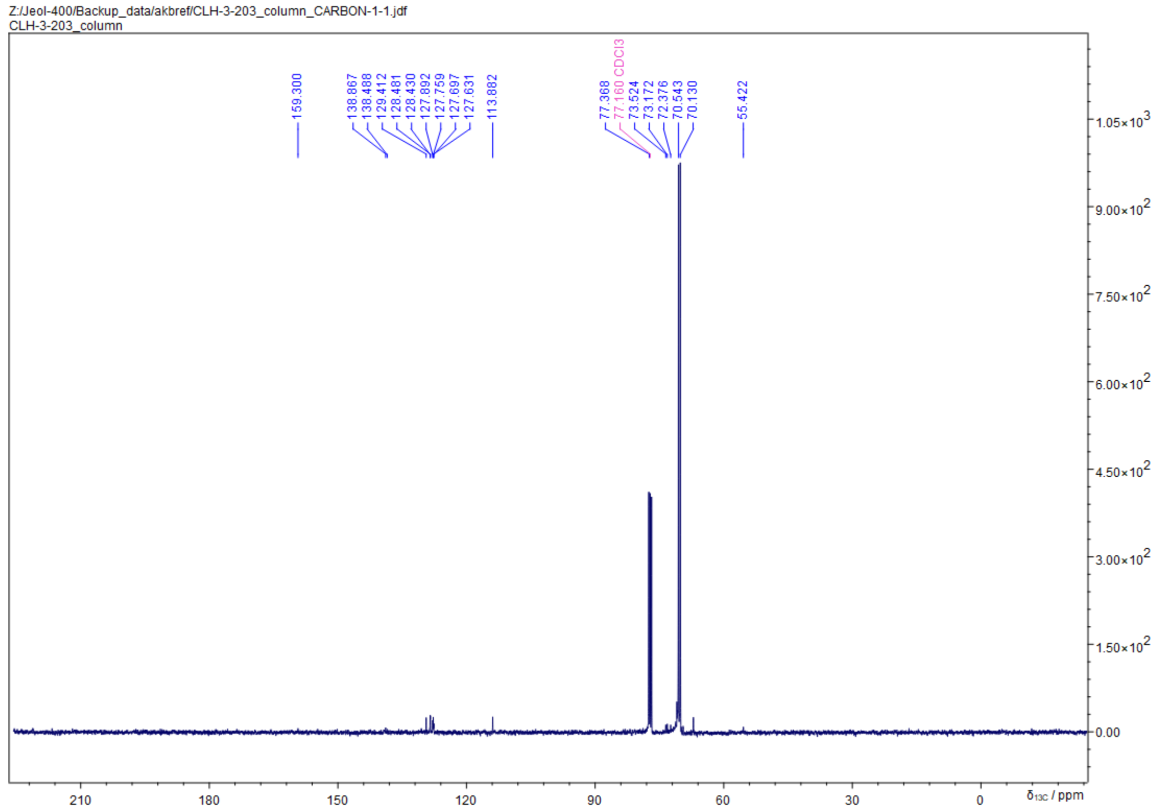

Z:/Jeol-400/Backup_data/akbref/CLH-3-203_column_CARBON-1-1.jdf
CLH-3-203_column
159.300
138.867
138.488
129.412
128.481
128.430
127.892
127.759
127.697
127.631
113.882
77.368
77.160 CDCl3
73.524
73.172
72.376
70.543
70.130
55.422
δ13C / ppm

**(((3-((4-Methoxybenzyl)oxy)propane-1,2-diyl-1,2,3-$^{13}C_3$)bis(oxy))bis(methylene))dibenzene (13)**

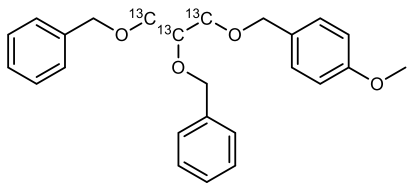


In a 100 mL round-bottom flask 547 mg (2.14 mmol, 1.0 eq.) **10** were dissolved in 10 mL 90% acetic acid. The reaction was stirred at 40 °C until full conversion was indicated by HPLC (3 h). The solvent was removed under reduced pressure to yield a yellow oil, which was used immediately in the next step without further purification. The 100 mL round bottom flask containing the intermediate was equipped with a Schlenk adapter and evacuated and then backfilled with Ar (3x). Then the intermediate was dissolved in a mixture of 40 mL THF and 20 mL $Et_2O$. The reaction mixture was cooled to 0 °C in an ice-bath, followed by addition of 222 mg (60% in mineral oil, 5.57 mmol, 2.6 eq.) NaH. A bubbler was connected to the Schlenk adapter and the suspension was stirred at 0 °C for 20 min until $H_2$-evolution stopped, which was accompanied by strong thickening of the solution. Subsequently, 610 µL (880 mg, 5.14 mmol, 2.4 eq.) BnBr were added and the ice-bath was removed. The reaction was stirred for 3 d at RT until full conversion was indicated via HPLC-MS and excess NaH was quenched by slow addition of 100 mL $H_2O$. The reaction mixture was extracted with 2 x 50 mL EtOAc and the combined organic phases were dried over $Na_2SO_4$. The volatiles were removed under reduced pressure to yield the crude product, which was further purified via column chromatography (50 mL $SiO_2$, CH/EE = 15:1, CAM, $R_f$ = 0.12) yielding 565 mg (66% yield) of a colorless oil.

| | |
|---|---|
| **$^{13}C_3C_{22}H_{28}O_4$** | [395.47 g/mol] |
| **Yield** | 564 mg (2.20 mmol, 66%), colorless oil |
| **TLC** | $R_f$ = 0.12 (CH/EtOAc = 15:1, UV and CAM) |
| **HPLC-MS** | $t_R$ = 5.828; m/z (%): [$MH^+$] = 396 (13), [$M(NH_4^+)$] = 413 (36), [$MNa^+$]= 418 (100) |

**$^1$H-NMR** (400 MHz, $CDCl_3$) δ = 7.13 - 7.31 (m, 12H), 6.76 - 6.83 (m, 2H), 4.62 (d, *J* = 3.5, 2H), 4.46 (d, *J* = 4.0, 2H), 4.39 (d, *J* = 4.0, 2H), 3.85 - 3.93 (m, 0H), 3.65 - 3.77 (m, 5H), 3.50 - 3.58 (m, 1H), 3.28 - 3.42 (m, 2H).

**$^{13}$C-NMR** (101 MHz, $CDCl_3$) δ = 159.3 (s), 138.9 (s), 138.5 (s), 130.5 (s), 129.4 (s), 128.5 (d, *J* = 5.3), 127.9 (s), 127.5 (s), 113.9 (s), 77.8 (s), 77.4 (s,), 77.0 (s), 73.5 (s), 73.2 (s), 72.4 (s), 70.8 (s), 70.3 (s), 69.9 (s), 55.4 (s)

Z:/Jeol-400/Backup_data/akbref/CLH-3-155B_PROTON-1-1.jdf
CLH-3-155B

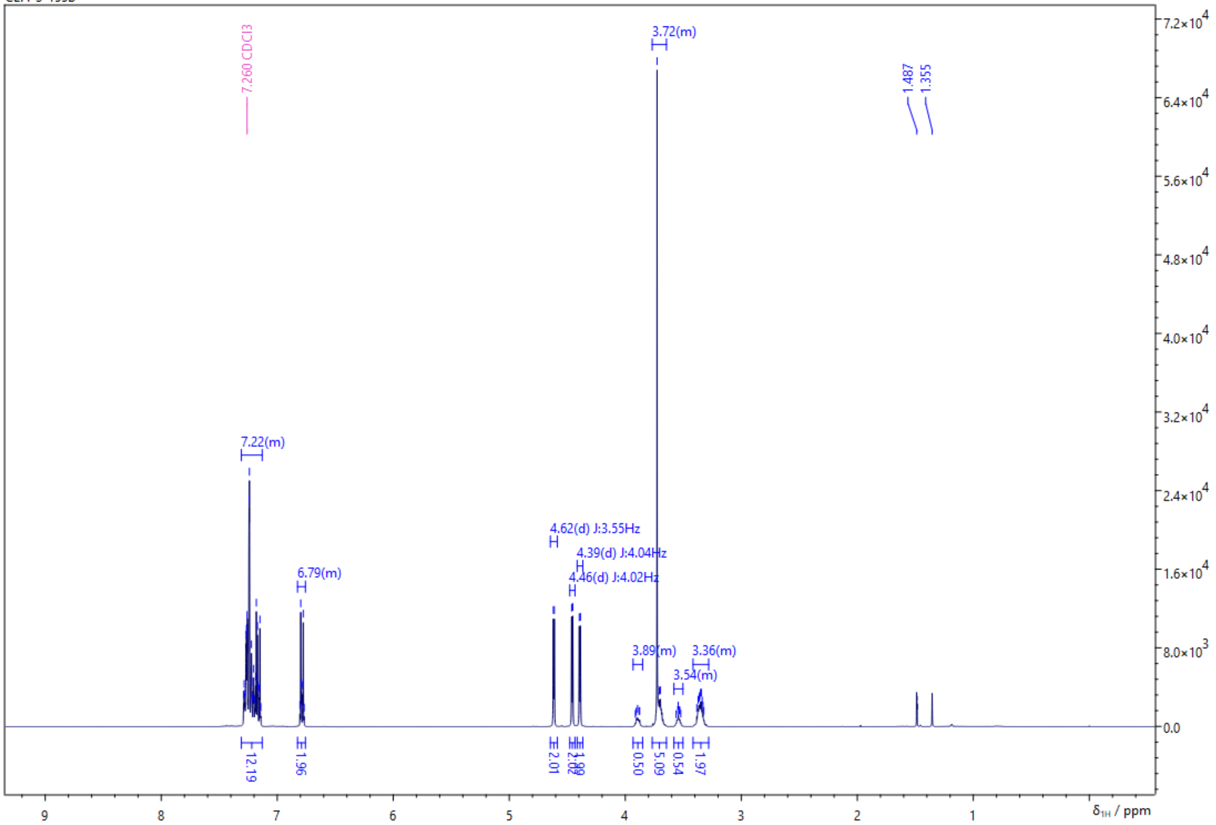


Z:/Jeol-400/Backup_data/akbref/CLH-3-155B_CARBON-1-1.jdf
CLH-3-155B

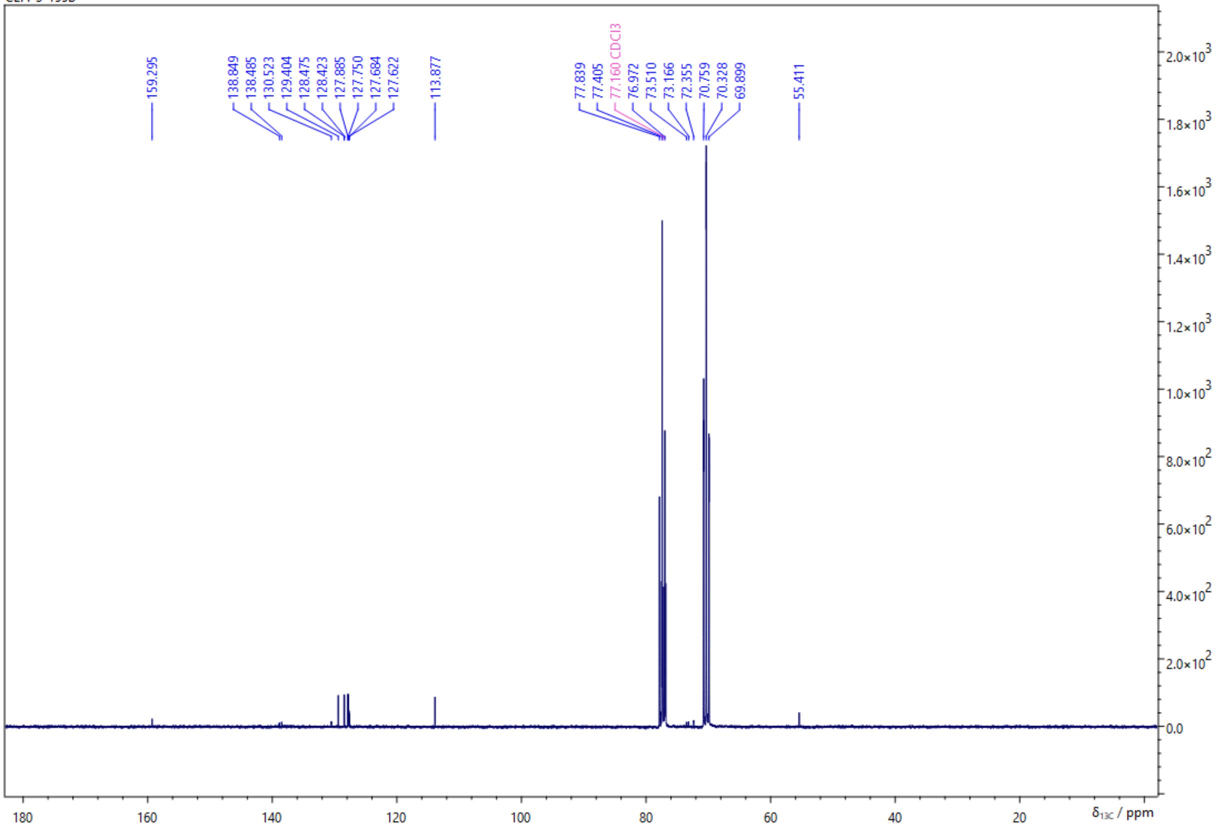

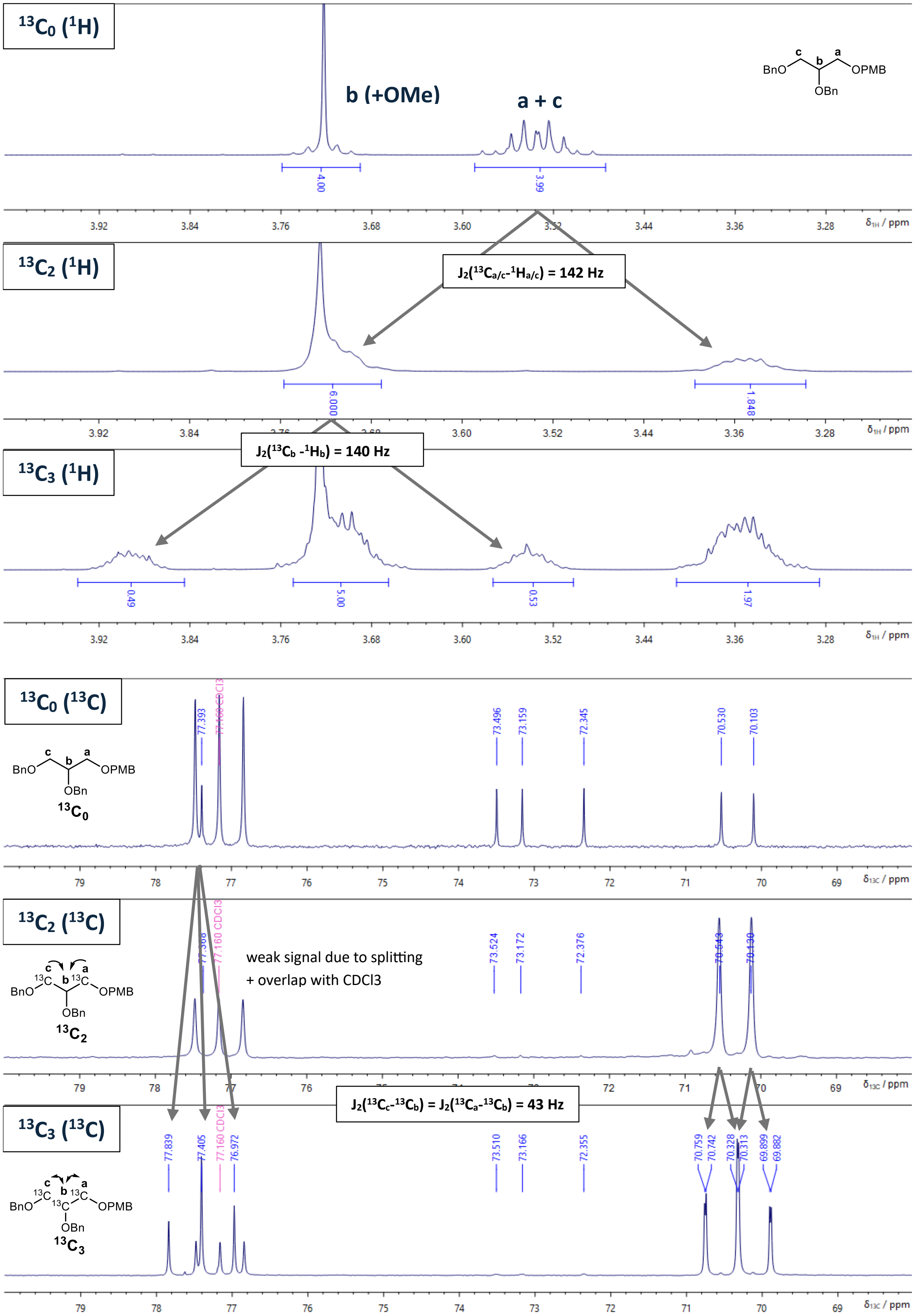

13C0 (1H)
b (+OMe)
a + c
BnO OPMB OBn
4.00
3.99
δ1H / ppm
13C2 (1H)
J2(13Ca/c-1Ha/c) = 142 Hz
6.000
1.848
13C3 (1H)
J2(13Cb -1Hb) = 140 Hz
0.49
5.00
0.53
1.97
13C0 (13C)
77.393
77.160 CDCl3
73.496
73.159
72.345
70.530
70.103
13C0
δ13C / ppm
13C2 (13C)
weak signal due to splitting
+ overlap with CDCl3
77.160 CDCl3
73.524
73.172
72.376
13C2
J2(13Cc-13Cb) = J2(13Ca-13Cb) = 43 Hz
13C3 (13C)
77.839
77.405
77.160 CDCl3
76.972
73.510
73.166
72.355
70.759
70.742
70.328
70.313
69.899
69.882
13C3

**2,3-Bis(benzyloxy)propan-1-ol (14)**

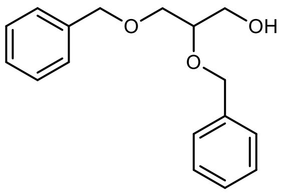


In an argon flushed Schlenk flask 505 mg (1.29 mmol, 1.0 eq.) **11** were dissolved in 5 mL degassed DCM and 250 µL $H_2O$. The reaction mixture was cooled to 0 °C in an ice bath, followed by the addition of 322 mg (1.42 mmol, 1.1 eq.) DDQ. The dark green emulsion was stirred at 0 °C and the progress of the reaction was accompanied by the formation of a light pink precipitate. After 4 h full conversion was indicated via HPLC-MS and the reaction was quenched by the addition of 5 mL sat. $NaHCO_3$, further diluted with 20 mL $H_2O$ and extracted with DCM (3 x 15 mL). The combined organic phases were dried over $Na_2SO_4$. The volatiles were removed under reduced pressure and the resulting yellow oil was purified via column chromatography (25 mL $SiO_2$, CH/EE = 4:1, CAM, $R_f$ = 0.14) to yield 272 mg (77%) of a slightly yellow oil.

| | |
|---|---|
| **$^{13}C_0C_{17}H_{20}O_3$** | [272.34 g/mol] |
| **Yield** | 272 mg (1.000 mmol, 77%), slightly yellow oil |
| **TLC** | $R_f$ = 0.14 (CH/EtOAc =4:1, UV and CAM) |
| **HPLC-MS** | $t_R$ = 4.297 min; m/z (%): [M+H$^+$] = 273 (22), [M+Na$^+$] = 295 (73), [M-Bn$^+$] = 181 (100) |

**$^1$H-NMR** (400 MHz, $CDCl_3$) δ = 7.27 - 7.40 (m, 10H), 4.72 (d, *J* = 11.7, 1H), 4.63 (d, *J* = 11.8, 1H), 4.55 (t, *J* = 12.8, 2H), 3.57 - 3.82 (m, 5H), 1.95 (bs, 1H).

**$^{13}$C-NMR** (101 MHz, $CDCl_3$) δ = 138.4 (s), 138.1 (s), 128.6 (s), 128.6 (s), 127.9 (s), 127.9 (s, 3C), 127.8 (s), 78.2 (s), 73.7 (s), 72.3 (s), 70.4 (s), 63.0 (s).

Z:/Jeol-400/Backup_data/akbref/CLH-3-146-2_PROTON-1-1.jdf
CLH-3-146-2

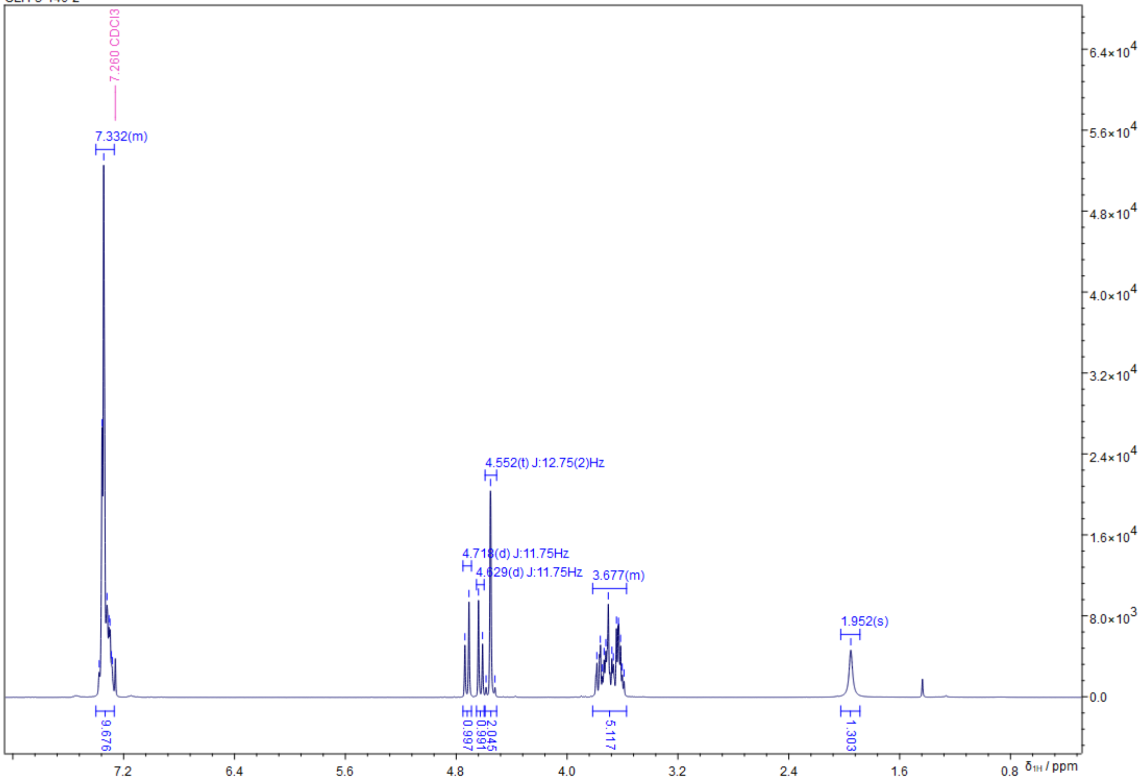


Z:/Jeol-400/Backup_data/akbref/CLH-3-146-2_CARBON-1-1.jdf
CLH-3-146-2

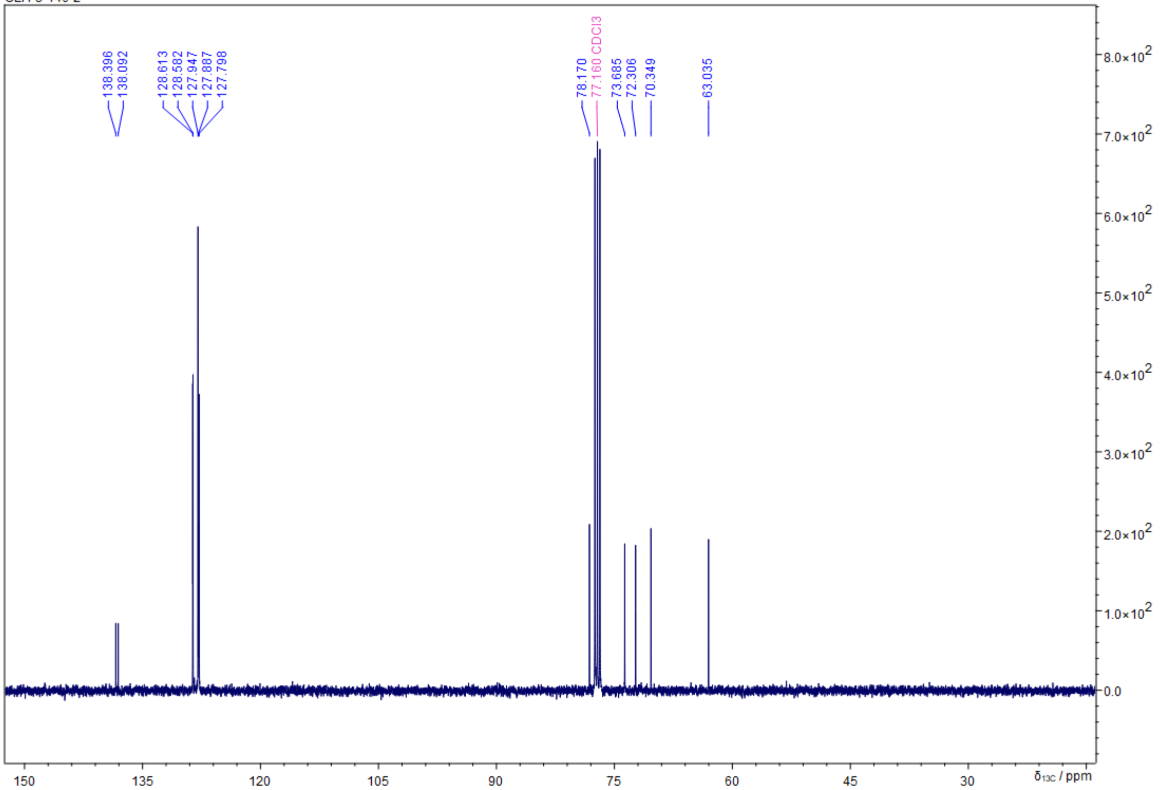

**2,3-Bis(benzyloxy)propan-1-ol-1,3-$^{13}C_2$ (15)**

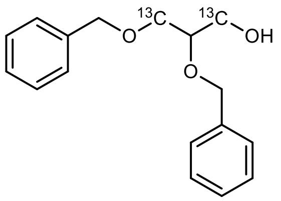


In an argon flushed Schlenk flask 189 mg (0.479 mmol, 1.0 eq.) **12** were dissolved in 5 mL degassed DCM and 250 µL $H_2O$. The reaction mixture was cooled to 0 °C in an ice bath, followed by the addition of 119 mg (0.527 mmol, 1.1 eq.) DDQ. The dark green emulsion was stirred at 0 °C and the progress of the reaction was accompanied by the formation of a light pink precipitate. After 4 h full conversion was indicated via HPLC-MS and the reaction was quenched by the addition of 5 mL sat. $NaHCO_3$, further diluted with 20 mL $H_2O$ and extracted with DCM (3 x 15 mL). The combined organic phases were dried over $Na_2SO_4$. The volatiles were removed under reduced pressure and the resulting yellow oil was purified via column chromatography (30 mL $SiO_2$, CH/EE = 4:1, CAM, Rf = 0.14) to yield 100 mg (76%) of a colorless oil.

| | |
|---|---|
| **$^{13}C_2C_{15}H_{20}O_3$** | [274.15 g/mol] |
| **Yield** | 100 mg (0.365 mmol, 76%), colorless oil |
| **TLC** | $R_f$ = 0.16 (CH/EtOAc =4:1, UV and CAM) |
| **HPLC-MS** | $t_R$ = 4.317 min; m/z (%): [M+H$^+$] = 275 (12), [M+Na$^+$] = 297 (68), [M-Bn$^+$] = 184 (100) |

**$^1$H-NMR** (400 MHz, $CDCl_3$) δ = 7.27 - 7.39 (m, 10H), 4.72 (d, *J* = 11.7, 1H), 4.63 (d, *J* = 11.7, 1H), 4.50 - 4.59 (m, 2H), 3.95 (d, *J* = 11.5, 0H), 3.75 - 3.90 (m, 2H), 3.72 (s, 1H), 3.59 (d, *J* = 11.6, 1H), 3.38 - 3.54 (m, 2H), 1.88 (s, 1H).
**$^{13}$C-NMR** (101 MHz, $CDCl_3$) δ = 138.4 (s), 138.1 (s), 128.6 (s), 128.6 (s), 127.9 (s), 127.9 (s), 127.8 (s), 73.7 (s), 72.3 (s), 71.4 (s), 70.4 (s), 63.1 (s).

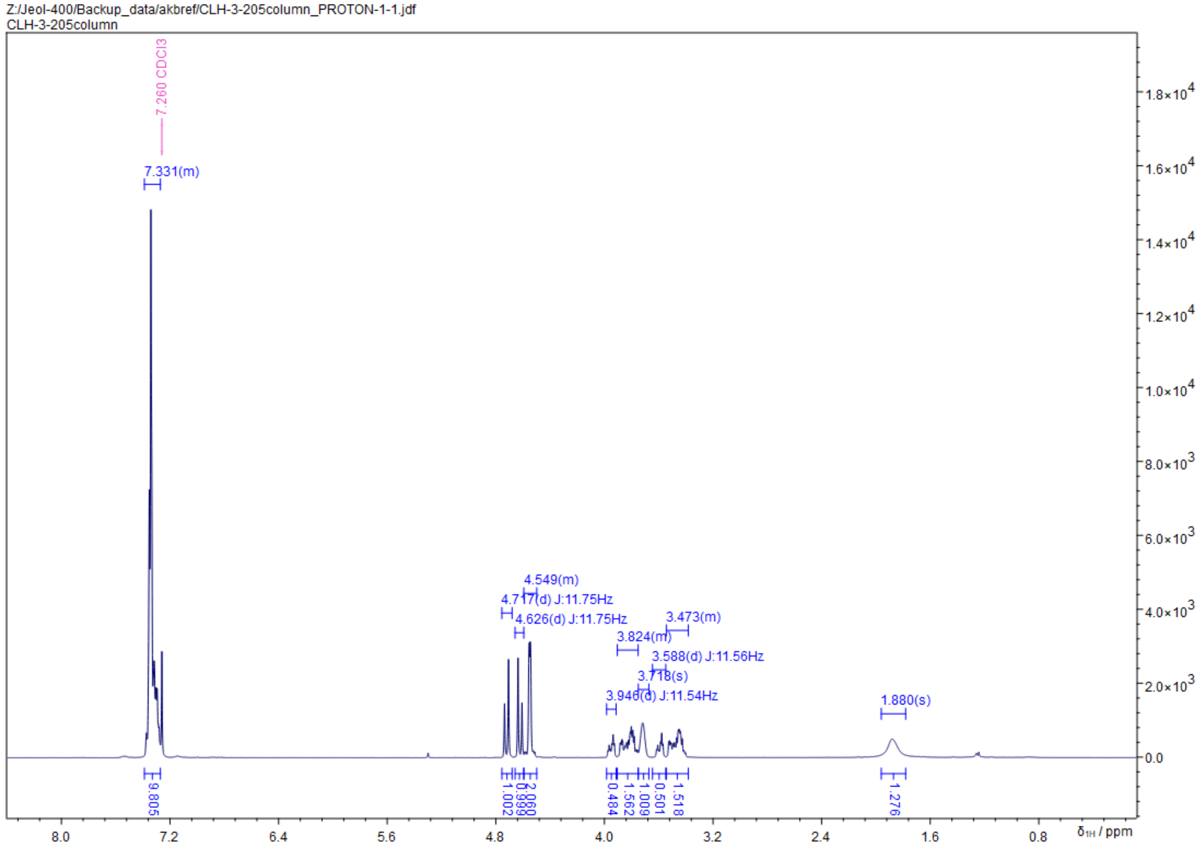

Z:/Jeol-400/Backup_data/akbref/CLH-3-205column_PROTON-1-1.jdf
CLH-3-205column
7.260 CDCl3
7.331(m)
4.549(m)
4.717(d) J:11.75Hz
4.626(d) J:11.75Hz
3.473(m)
3.824(m)
3.588(d) J:11.56Hz
3.718(s)
3.946(d) J:11.54Hz
1.880(s)
9.805
1.002
0.999
2.060
0.484
1.562
1.009
0.501
1.518
1.276
δ1H / ppm


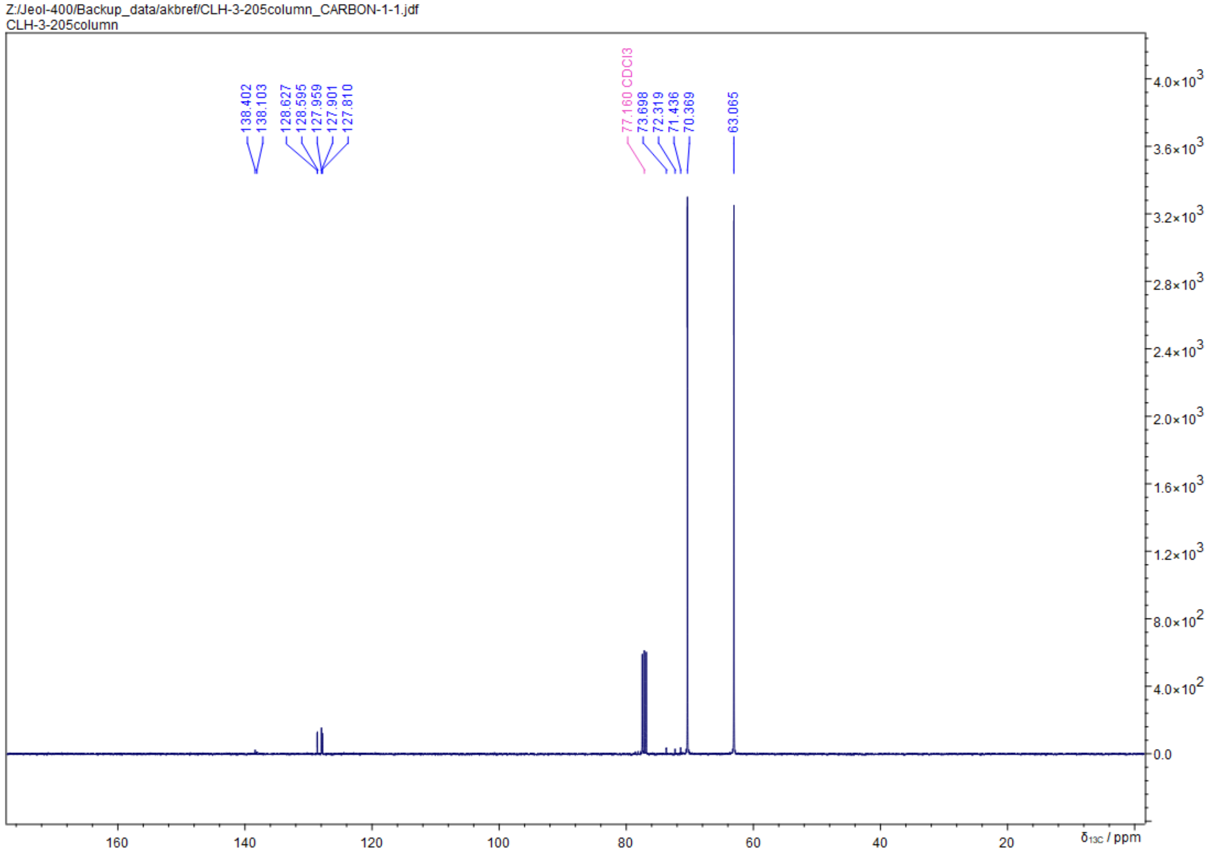

Z:/Jeol-400/Backup_data/akbref/CLH-3-205column_CARBON-1-1.jdf
CLH-3-205column
138.402
138.103
128.627
128.595
127.959
127.901
127.810
77.160 CDCl3
73.698
72.319
71.436
70.369
63.065
δ13C / ppm

**2,3-Bis(benzyloxy)propan-1-ol-1,2,3-13C$_3$ (16)**

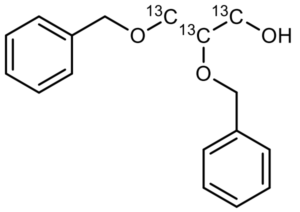


In an argon flushed Schlenk flask 410 mg (1.04 mmol, 1.0 eq.) **13** were dissolved in 10 mL degassed DCM and 500 µL $H_2O$. The reaction was cooled to 0 °C in an ice bath, followed by the addition of 260 mg (1.15 mmol, 1.1 eq.) DDQ. The dark green emulsion was stirred at 0 °C and the progress of the reaction was accompanied by the formation of a light pink precipitate. After 4 h full conversion was indicated via HPLC-MS and the reaction was quenched by the addition of 10 mL sat. $NaHCO_3$, further diluted with 40 mL $H_2O$ and extracted with DCM (3 x 30 mL). The combined organic phases were dried over $Na_2SO_4$. The volatiles were removed under reduced pressure and the resulting yellow oil was purified via column chromatography (30 mL $SiO_2$, CH/EE = 4:1, CAM, Rf = 0.14) to yield 221 mg (77%) of a colorless oil.

| | |
|---|---|
| **$^{13}C_3C_{14}H_{20}O_3$** | [275.15 g/mol] |
| **Yield** | 221 mg (0.806 mmol, 77%), colorless oil |
| **TLC** | $R_f$ = 0.14 (CH/EtOAc =4:1, UV and CAM) |
| **HPLC-MS** | $t_R$ = 4.299 min; m/z (%): [M+H$^+$] = 276 (15), [M+Na$^+$] = 298 (84), [M-Bn$^+$] = 185 (100) |

**$^1$H-NMR** (400 MHz, $CDCl_3$) δ = 7.27 - 7.39 (m, 10H), 4.72 (dd, *J* = 2.8, 11.7, 1H), 4.63 (dd, *J* = 2.7, 11.7, 1H), 4.50 - 4.59 (m, 2H), 3.678 (d, *J* = 141Hz, 5H), 1.93 (s, 1H).
**$^{13}$C-NMR** (101 MHz, $CDCl_3$) δ = 138.4 (s), 138.1 (s), 128.6 (s), 128.6 (s), 128.0 (s), 127.9 (s), 127.8 (s), 78.1 (dd, *J* = 40.8, 43.5, **$^{13}$C**), 72.3 (s), 70.3 (dd, *J* = 1.6, 43.7, **$^{13}$C**), 63.1 (dd, *J* = 1.4, 40.6, **$^{13}$C**).

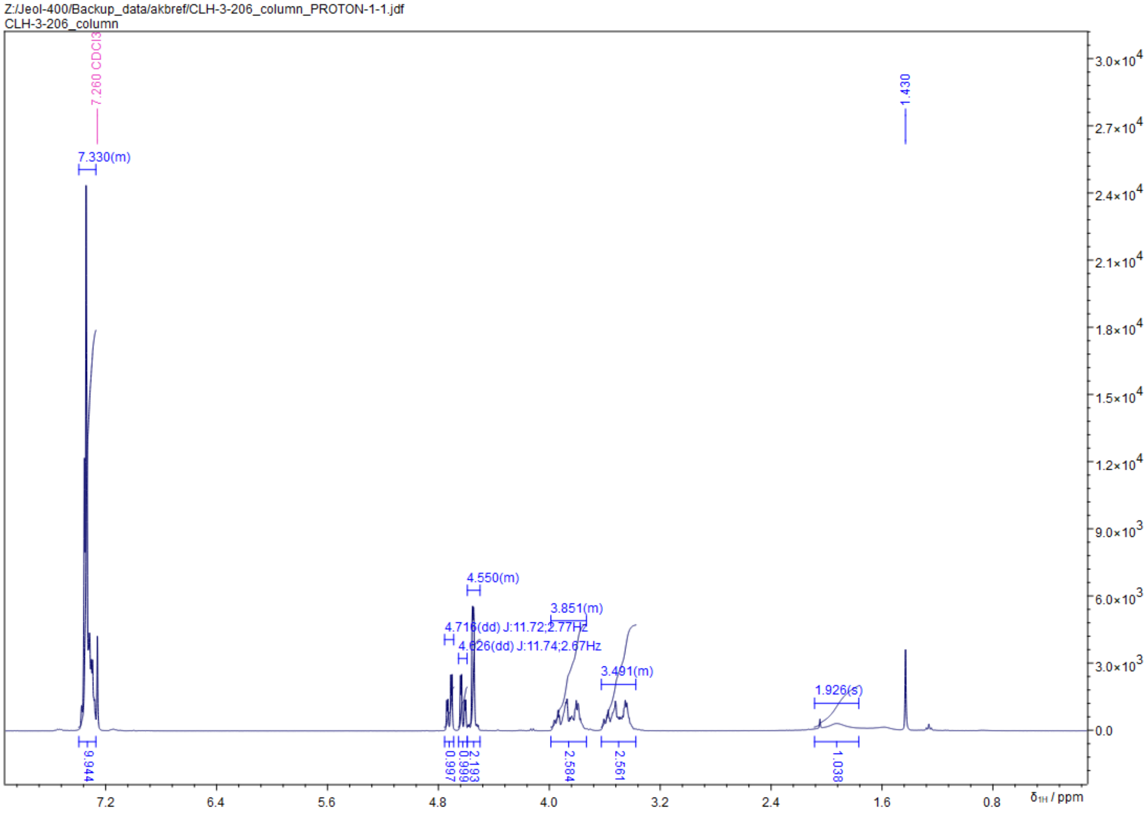

Z:/Jeol-400/Backup_data/akbref/CLH-3-206_column_PROTON-1-1.jdf
CLH-3-206_column
7.260 CDCl3
1.430
7.330(m)
4.550(m)
3.851(m)
4.716(dd) J:11.72;2.77Hz
4.626(dd) J:11.74;2.67Hz
3.491(m)
1.926(s)
9.944
0.997
0.999
2.193
2.584
2.561
1.038
δ1H / ppm


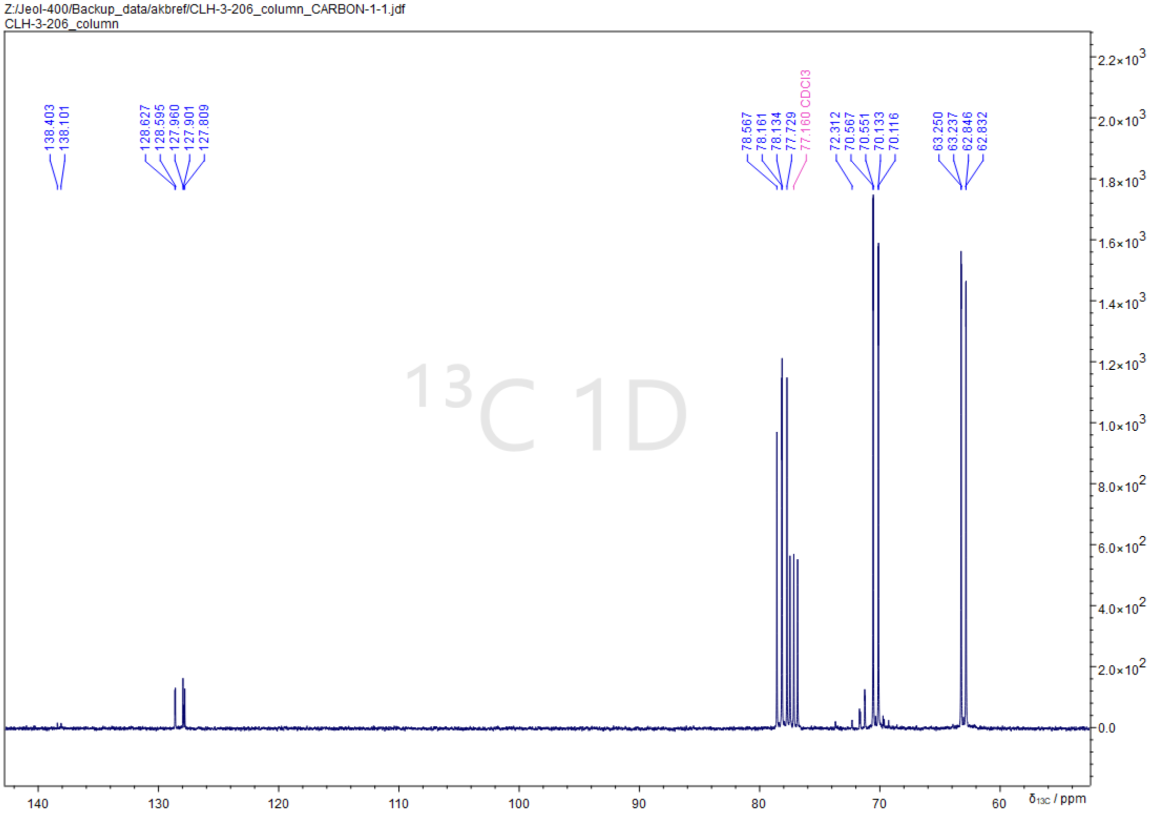

Z:/Jeol-400/Backup_data/akbref/CLH-3-206_column_CARBON-1-1.jdf
CLH-3-206_column
138.403
138.101
128.627
128.595
127.960
127.901
127.809
78.567
78.161
78.134
77.729
77.160 CDCl3
72.312
70.567
70.551
70.133
70.116
63.250
63.237
62.846
62.832
13C 1D
δ13C / ppm

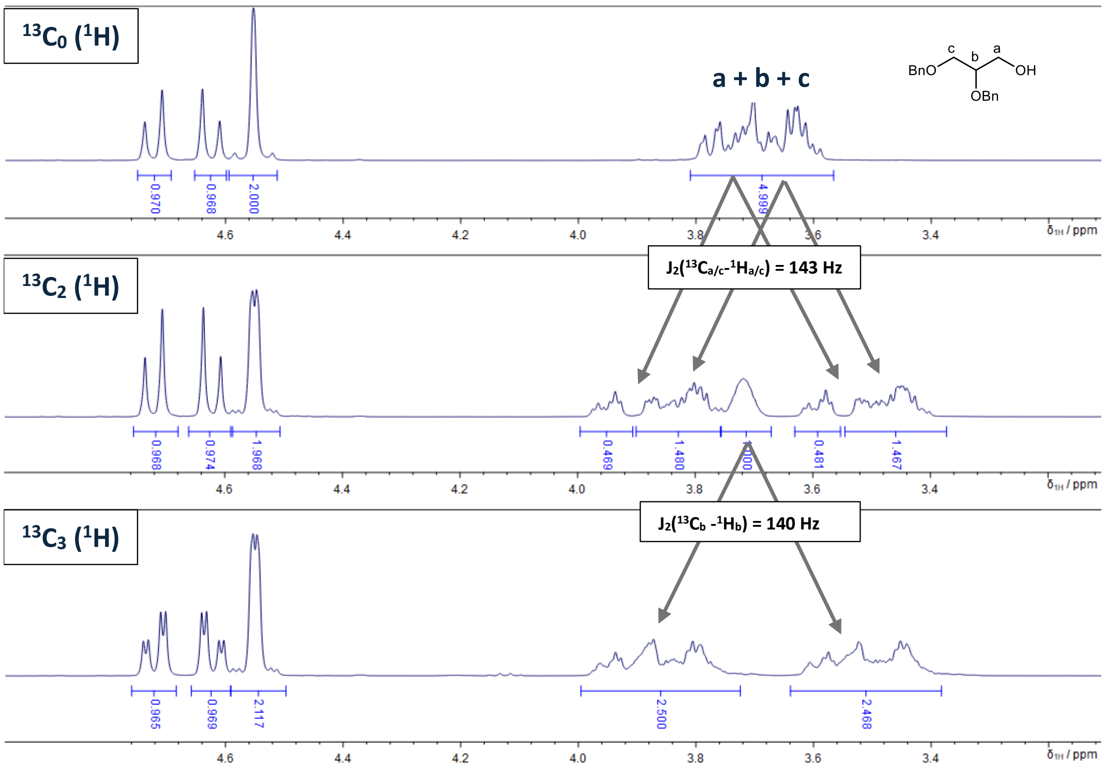

$^{13}C_0$ ($^1H$)
a + b + c
BnO c b a OH OBn
$J_2(^{13}C_{a/c}-^{1}H_{a/c})$ = 143 Hz
$^{13}C_2$ ($^1H$)
$J_2(^{13}C_b-^{1}H_b)$ = 140 Hz
$^{13}C_3$ ($^1H$)
δ1H / ppm


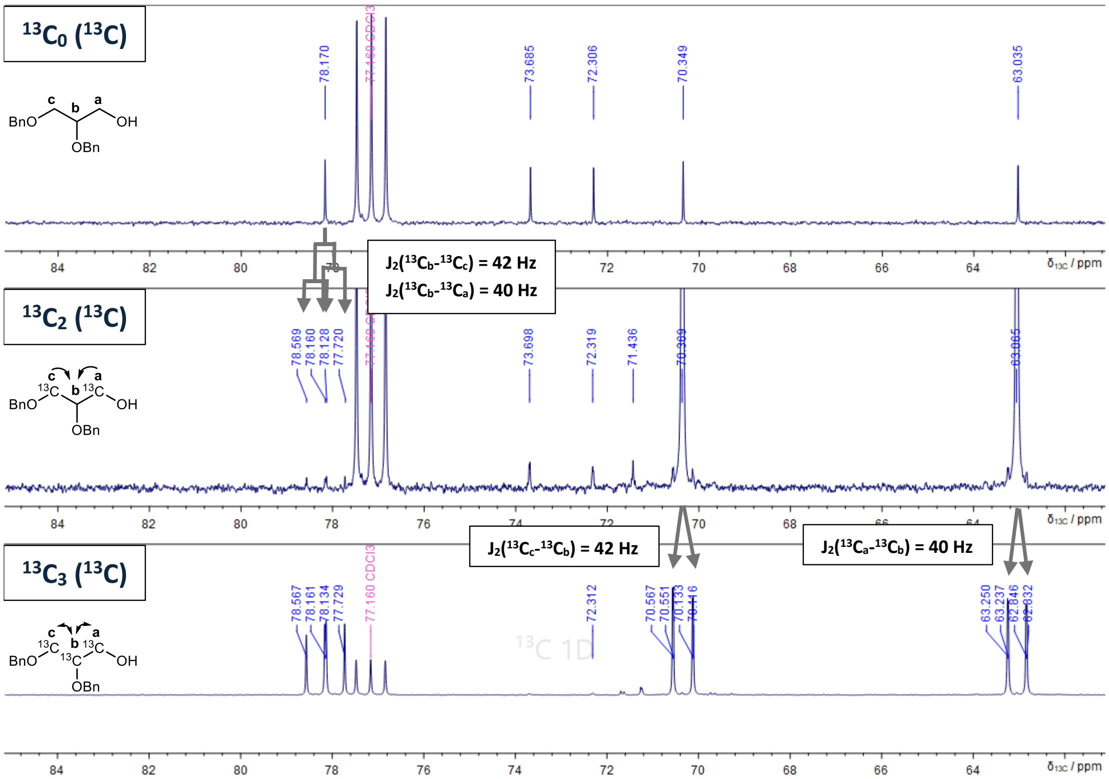

$^{13}C_0$ ($^{13}C$)
78.170
77.160 CDCl3
73.685
72.306
70.349
63.035
$J_2(^{13}C_b-^{13}C_c)$ = 42 Hz
$J_2(^{13}C_b-^{13}C_a)$ = 40 Hz
$^{13}C_2$ ($^{13}C$)
78.569
78.160
78.128
77.720
73.698
72.319
71.436
$J_2(^{13}C_c-^{13}C_b)$ = 42 Hz
$J_2(^{13}C_a-^{13}C_b)$ = 40 Hz
$^{13}C_3$ ($^{13}C$)
78.567
78.161
78.134
77.729
77.160 CDCl3
72.312
70.567
70.551
70.133
63.250
63.237
62.846
62.832
δ13C / ppm

**bBenzyl (2,3-bis(benzyloxy)propyl-1,2,3-$^{13}C_3$) diisopropylphosphoramidite (17)**

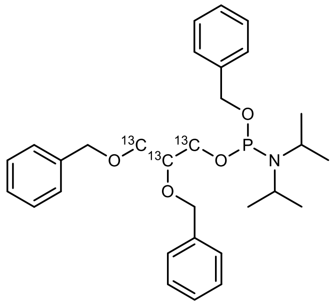


In an argon flushed Schlenk flask 553 mg (1.56 mg, 2.0 eq.) **15** and 215 mg (0.780 mmol, 1.0 eq.) 1-(benzyloxy)-N,N,N',N'-tetraisopropylphosphanediamine were azeotropically dried by the addition and evaporation of dry toluene. The dried reagents were dissolved in 6 mL dry DCM, followed by the addition of a solution of 185 mg (1.56 mmol, 2.0 eq.) 4,5-dicyanoimidazole in 2 mL dry ACN. The reaction was stirred for 15 h at RT and solvent was removed under reduced pressure to obtain a yellow oil, which was immediately purified by two fast (otherwise decomposition on silica gel) rounds of column chromatography (20 mL $SiO_2$, $Et_2O$/pentane = 1:10 + 5% $Et_3N$, CAM, Rf = 0.78) yielding 144 mg (36% yield) of a colorless oil.

| | |
|---|---|
| **$^{13}C_3C_{27}H_{40}NO_4P$** | [512.60 g/mol] |
| **Yield** | 144 mg (0.281 mmol, 36%), colorless oil |
| **TLC** | $R_f$ = 0.78 ($Et_2O$/pentane = 1:10 + 5% $Et_3N$, UV and CAM) |
| **HPLC-MS** | P-N(iPr)$_2$ hydrolysis; $t_R$ = 5.190 min (m/z (%): [M+H$^+$] = 430 (100), [M+Na$^+$] = 452 (31), [M+HCOOH$^+$] = 475 (34) |

**$^1$H-NMR** (400 MHz, $CDCl_3$) δ = 7.20 - 7.42 (m, 15H), 4.57 - 4.81 (m, 4H), 4.54 (d, *J* = 3.5, 2H), 3.24 - 4.14 (m, 7H), 1.17 (q, *J* = 5.65 Hz).
**$^{13}$C-NMR** (101 MHz, $CDCl_3$) δ = 139.7 (s), 138.9 (s), 128.5 (s), 128.4 (s), 128.4 (s), 127.9 (s), 127.8 (s), 127.7 (s), 127.7 (s), 127.6 (s), 127.6 (s), 127.4 (s), 127.3 (s), 127.1 (d, *J* = 1.9), 78.1 (tdd, *J* = 2.7, 7.6, 43.8, **$^{13}$C**), 70.5 (d, *J* = 43.4, **$^{13}$C**) 63.2 (dddd, *J* = 1.7, 7.3, 15.7, 44.0, **$^{13}$C**), 43.1 (d, *J* = 13.4), 24.8 (dd, *J* = 6.6, 4.5)
**$^{31}$P-NMR** (162 MHz, $CDCl_3$) δ = 147.67 - 148.57 (m)

Z:/Jeol-400/Backup_data/akbref/CLH-3-207column2_PROTON-1-1.jdf
CLH-3-207column2

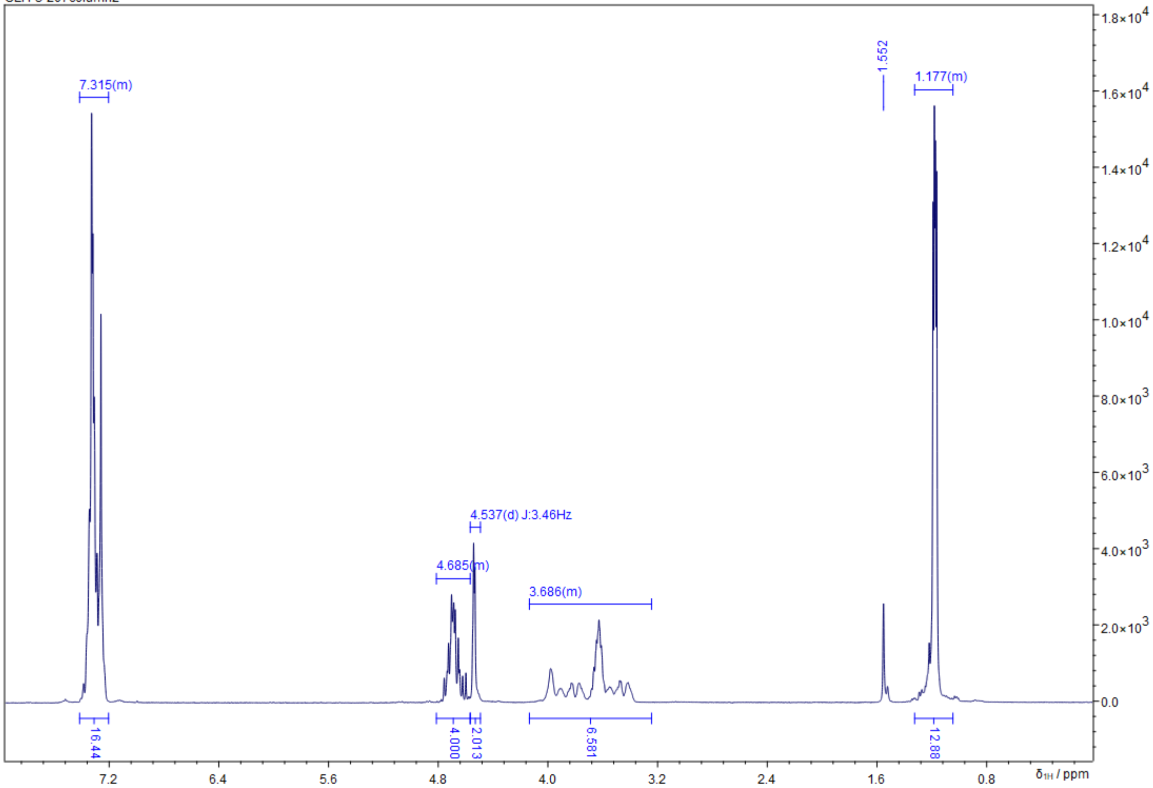


Z:/Jeol-400/Backup_data/akbref/CLH-3-207column2_CARBON-1-1.jdf
CLH-3-207column2

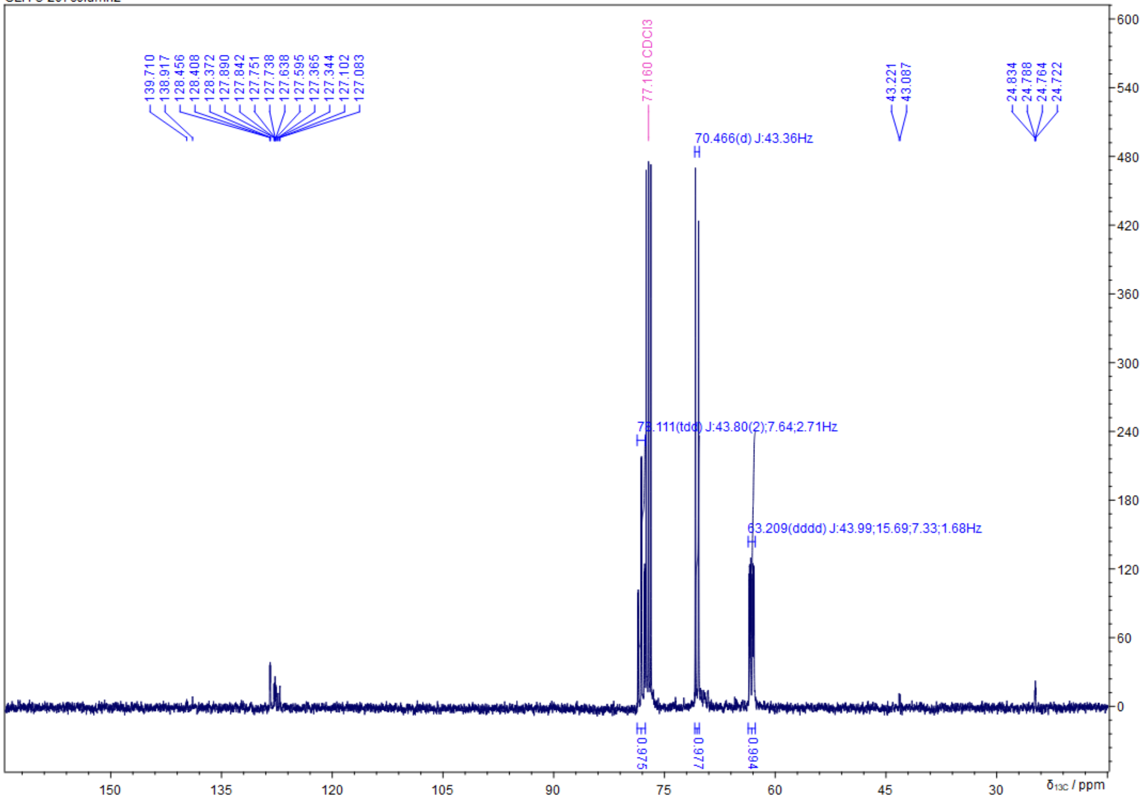

Z:/Jeol-400/Backup_data/akbref/CLH-3-207column2_PHOSPHORUS-1-1.jdf
CLH-3-207column2

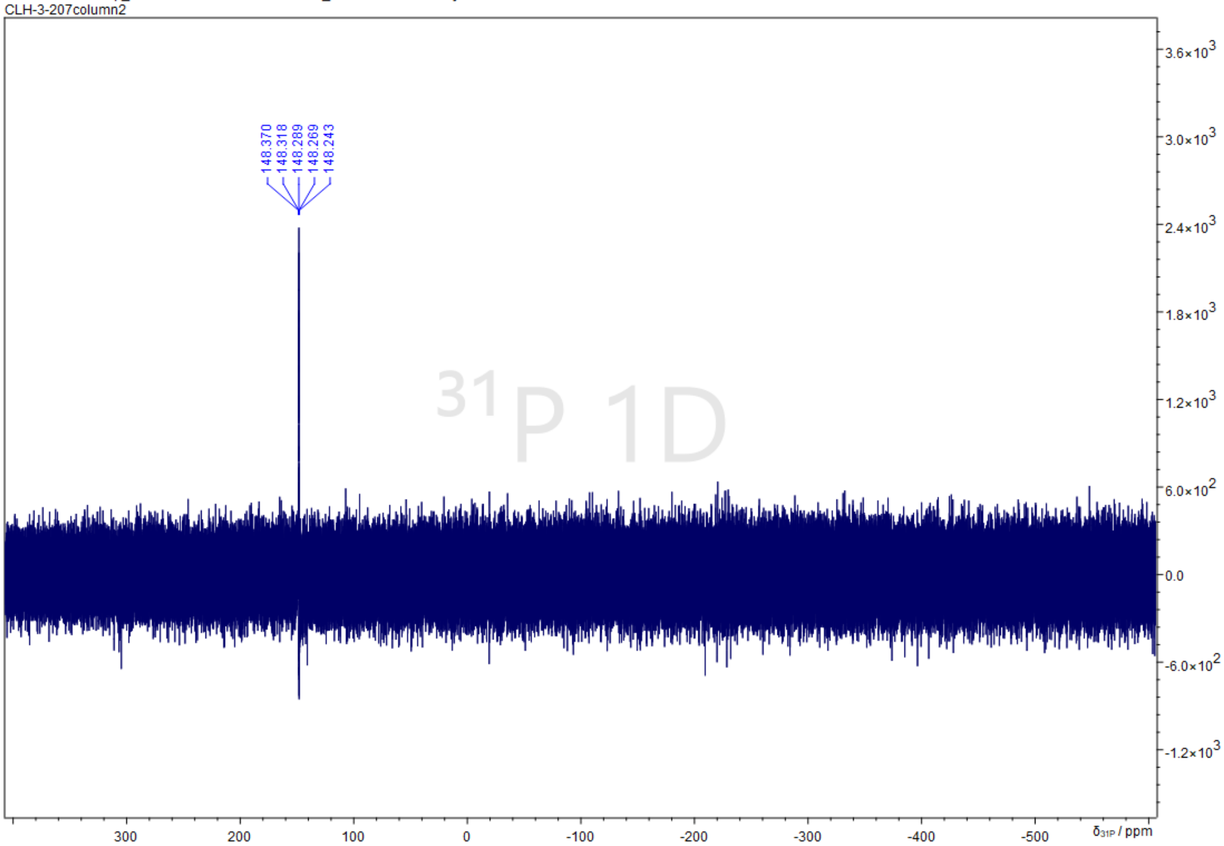

**Benzyl (2,3-bis(benzyloxy)propyl-1,2,3-13C3) (2,3-bis(benzyloxy)propyl-1,3-$^{13}C_2$) phosphate (18)**

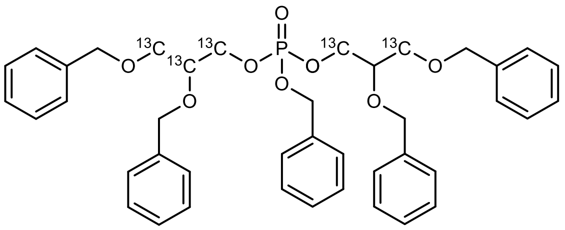

In an argon flushed Schlenk flask 136 mg (0.265 mmol, 1.0 eq.) **17** and 80 mg (0.291 mmol, 1.1 eq.) **16** were azeotropically dried by the addition and evaporation of dry toluene. The dried reagents were dissolved in 6 mL dry DCM, followed by the addition of a solution of 35 mg (0.291 mmol, 1.1 eq.) 4,5-dicyanoimidazole in 2 mL dry ACN. After stirring at RT for 2 d, the reaction mixture was cooled to -20 °C using a salt/ice-bath and 101 mg (0.400 mmol, 1.5 eq.) m-CPBA were added. Complete oxidation was indicated by HPLC-MS after 2 h of stirring at -20 °C and the reaction was quenched by the addition of 20 mL sat. $NaHCO_3$. After extraction with DCM (3 x 20 mL), the combined organic phases were dried over $Na_2SO_4$ and the volatiles removed under reduced pressure to obtain a yellow oil, which was purified via column chromatography (20 mL $SiO_2$, CH/EE = 2:1, CAM, Rf = 0.55) to yield 140 mg of the desired product, which was contaminated with 5% (16 mol%) **17.**

| | |
|---|---|
| **$^{13}C_5C_{36}H_{45}NO_8P$** | [701.74 g/mol] |
| **Yield** | 140 mg, 95% purity, (0.189 mmol, 71%), colorless oil |
| **TLC** | $R_f$ = 0.55 (CH/EE = 2:1, UV and CAM, Rf = 0.55) |
| **HPLC-MS** | P-N(iPr)$_2$ hydrolysis; $t_R$ = 5.190 min (m/z (%): [M+H$^+$] = 430 (100), [M+Na$^+$] = 452 (31), [M+HCOOH$^+$] = 475 (34) |

**$^1$H-NMR** (400 MHz, $CDCl_3$) δ = 7.21 - 7.40 (m, 30H), 5.01 (d, *J* = 7.8, 2H), 4.50 - 4.77 (m, 6H), 4.48 (s, 4H), 4.34 (d, *J* = 30.7, 2H), 3.96 (d, *J* = 30.9, 3H), 3.72 (d, 4H), 3.29 - 3.62 (m, 3H)
**$^{13}$C-NMR** (101 MHz, $CDCl_3$) δ = δ = 138.3 (s), 138.1 (s), 128.7 (s), 128.6 (s), 128.6 (s), 128.5 (s), 128.5 (s), 128.0 (s), 127.9 (s), 127.9 (s), 127.8 (s), 127.8 (s), 76.5 (dd, *J* = 6.9, 43.6, **1$^{13}$C**), 69.3 (t, *J* = 20.3, **2$^{13}$C**), 67.1 (dt, *J* = 5.8, 21.3, **2$^{13}$C**)
**$^{31}$P-NMR** (162 MHz, $CDCl_3$) δ = -0.23 (m).

Z:/Jeol-400/Backup_data/akbref/CLH-3-208_column_PROTON-1-1.jdf
CLH-3-208_column

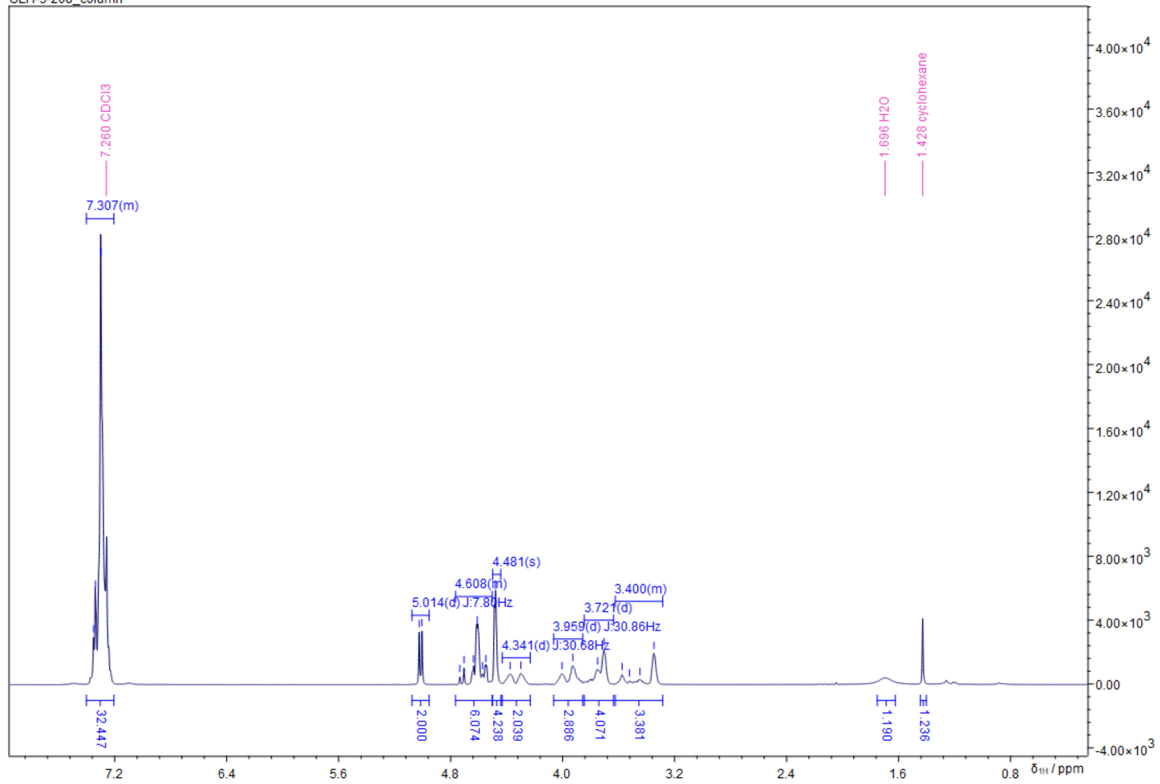


Z:/Jeol-400/Backup_data/akbref/CLH-3-208_column_CARBON-1-1.jdf
CLH-3-208_column

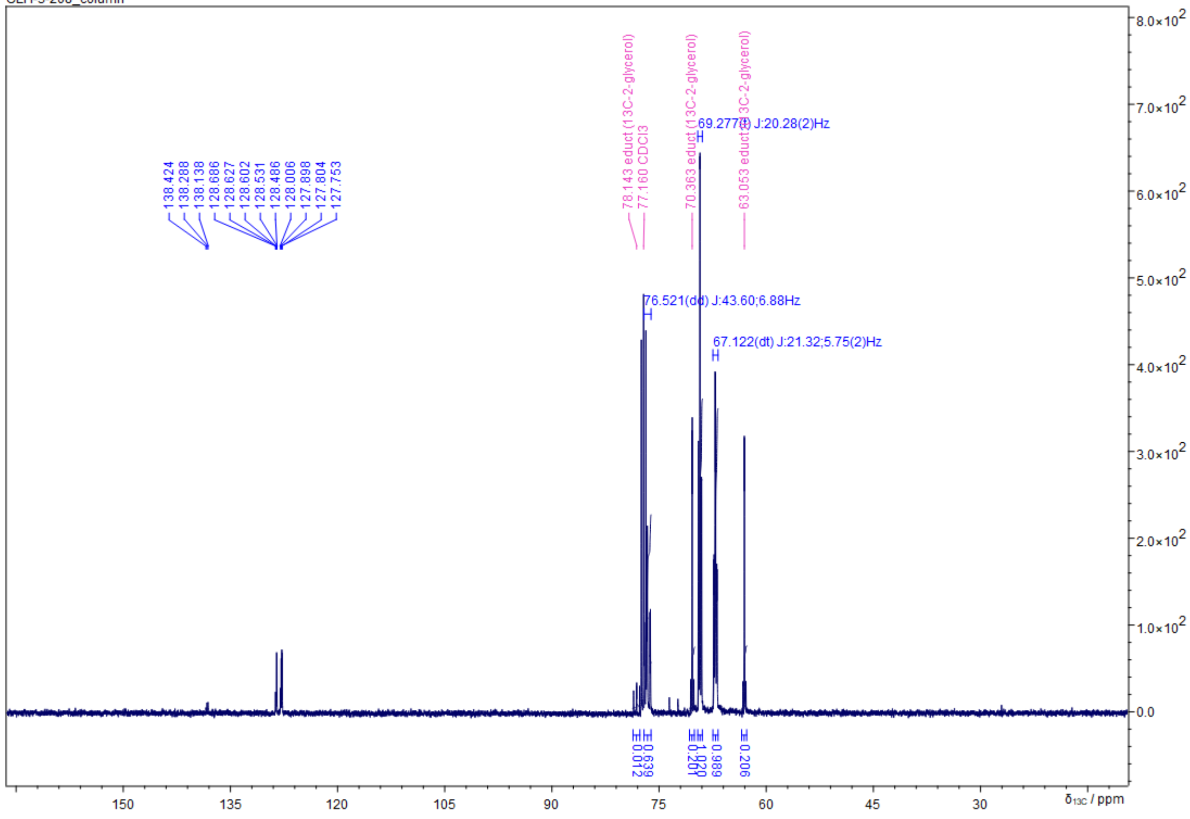

Z:/Jeol-400/Backup_data/akbref/CLH-3-208_column_PHOSPHORUS-1-1.jdf
CLH-3-208_column

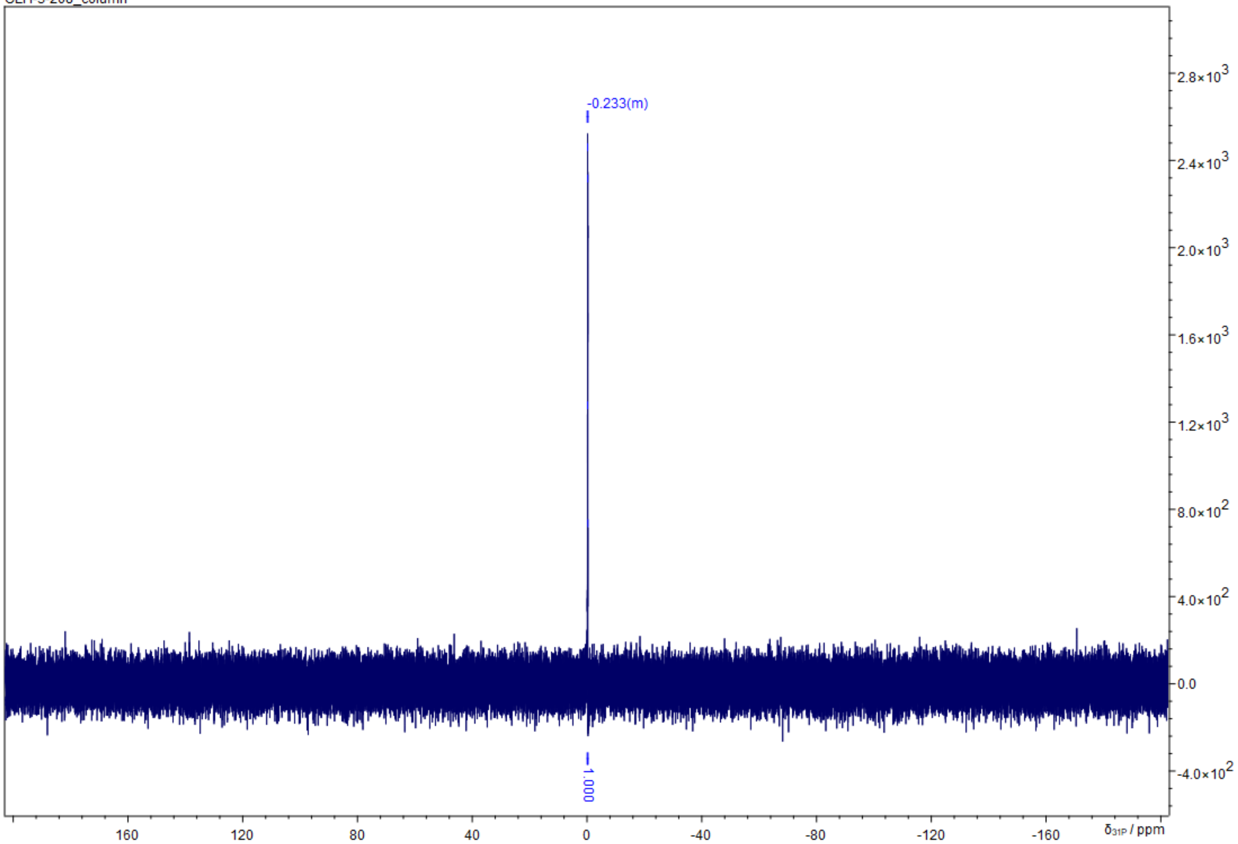

**Ammonium (*rac*-glyceryl-1,2,3-$^{13}C_3$) (*rac*-glyceryl -1,3-$^{13}C_2$) phosphate (GPG-$^{13}C_5$) (19)**

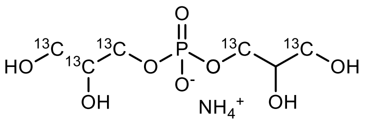


In an argon flushed 100 mL round-bottom flask with a Schlenk adapter 137 mg (0.195 mmol, 1.0 eq.) **18** were dissolved in 5 mL dry iPrOH, followed by the addition of 22 mg (10% Pd, 0.020 mmol, 0.1 eq.) Pd/C. The flask was carefully evacuated and flushed with $H_2$-gas (3 x), then placed in a 40 °C oil-bath and stirred for 3 d until HPLC-MS indicated full conversion of starting material. The reaction was subsequently diluted with 5 mL $H_2O$ and the Pd-catalyst removed by filtration under argon through a pad of celite (1 cm). The filter pad was rinsed using 20 mL $H_2O$. After lyophilization of the crude product, it was purified by reverse-phase column chromatography (18C-$SiO_2$, 20 mL, $H_2O$, CAM). As a result from contamination in the starting material however, the product was contaminated with unbound $^{13}C_2$-glycerol and required further purification with ion-exchange chromatography (Sephadex DEAE, 20 mL, conditioned to 0.1M $NH_4OAc$ pH = 5, washing with 100 mL $H_2O$, elution with 100 mL 0.5 M $NH_4OH$) resulting in 22 mg (44% yield) $^{13}C_5$-GPG ammonium salt as an amorphous solid.

**$^{13}C_5C_1H_{15}O_8P$** [251.11 g/mol]

**Yield** 22 mg, (0.087 mmol, 44%), amorphous solid

**HPLC-MS** $t_R$ = 0.460 min (m/z (%): [M+H$^+$] = 252 (48), [M+Na$^+$] = 274 (57), [M$_2$+Na$^+$] = 525 (100)

**$^1$H-NMR** (400 MHz, $CDCl_3$) δ = 3.97 - 4.17 (m, 2.5H), 3.81 - 3.97 (m, 2H), 3.60 - 3.81 (m, 3.5H), 3.34 - 3.60 (m, 2H).
**$^{13}$C-NMR** (101 MHz, $CDCl_3$) δ = 70.7 (td, *J* = 7.8, 41.8, **1$^{13}$C**, glycerol-$^{13}C_3$), 66.3 (ddd, *J* = 2.9, 5.4, 42.2, **1$^{13}$C**, glycerol-$^{13}C_3$ + dd, *J* = 2.9, 5.4, **1$^{13}$C**, glycerol-$^{13}C_2$), 62.1 (dd, *J* = 2.8, 41.2, **1$^{13}$C**, glycerol-$^{13}C_3$ + d, *J* = 2.6, **1$^{13}$C**, glycerol-$^{13}C_2$)
**$^{31}$P-NMR** (162 MHz, $CDCl_3$) δ = -0.23 (m).

Z:/Jeol-400/Backup_data/akbref/CLH-3-209_clean_PROTON-1-1.jdf
CLH-3-209_clean

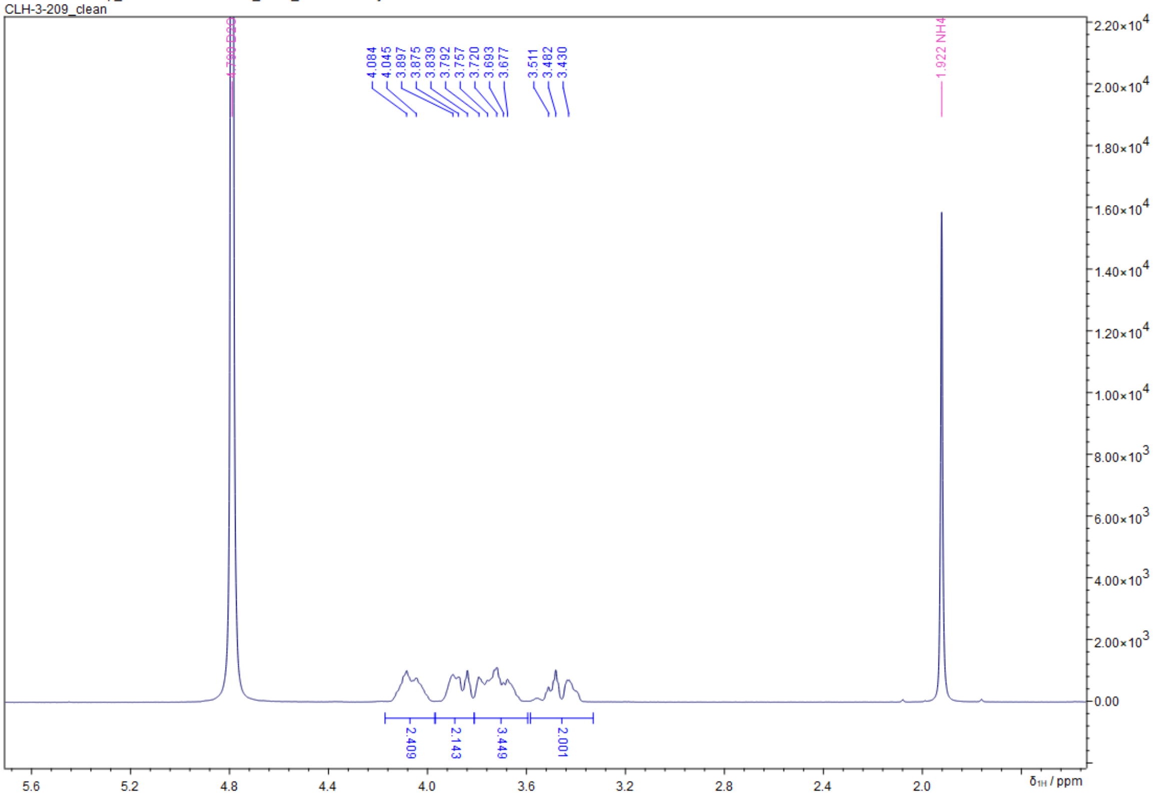


Z:/Jeol-400/Backup_data/akbref/CLH-3-209_clean_CARBON-1-1.jdf
CLH-3-209_clean

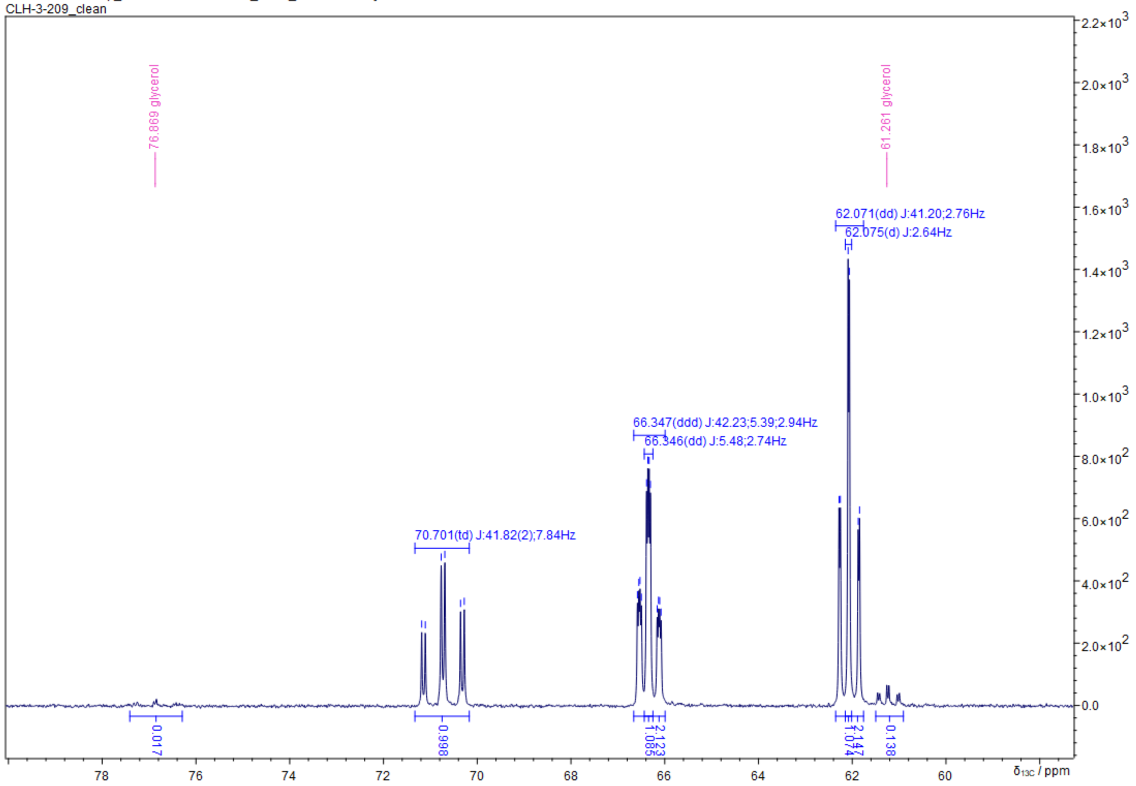

Z:/Jeol-400/Backup_data/akbref/CLH-3-209_clean_PHOSPHORUS-1-1.jdf
CLH-3-209_clean

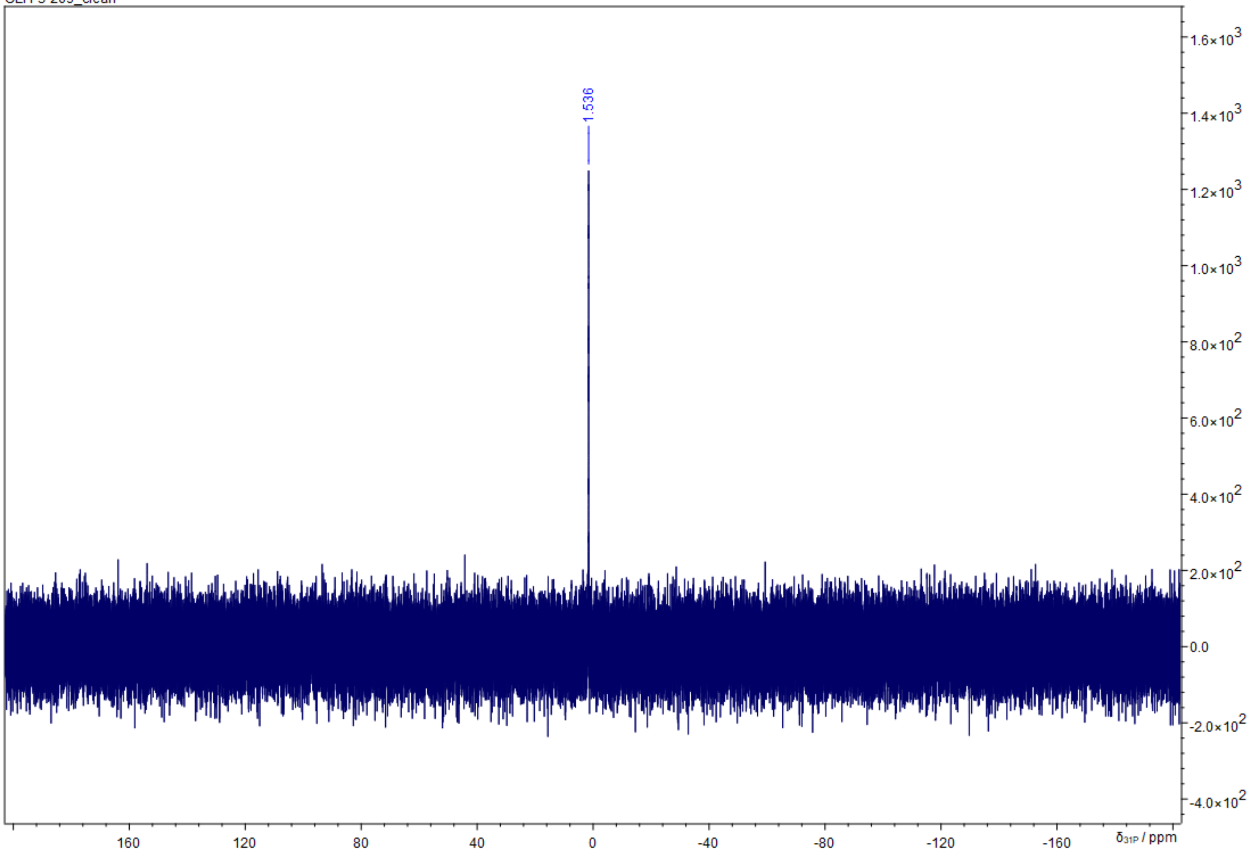
