## Supplementary Tables for "CLN8 enables a non-canonical phospholipid synthesis pathway"

### Supplementary table 1. Chemicals

| Chemical Name | Company | Catalog number |
|---|---|---|
| DMEM 4.5 g/L glucose | Gibco, Thermo Fisher Scientific, Waltham, USA | 11965092 |
| IMDM (Iscove's Modified Dulbecco's Medium) | Gibco, Thermo Fisher Scientific, Waltham, USA | 12440053 |
| Expi293T Expression Medium | Gibco, Thermo Fisher Scientific, Waltham, USA | A1435101 |
| 125 ml Vented Erlenmeyer flasks | Corning, NY, USA | 431143 |
| 50 ml CELLSTAR cell reactor tubes | Greiner Bio-One, Kremsmünster, Austria | 227245 |
| 35 mm glass bottom dishes | Ibidi, Graefelfing, Germany | 81218-200 |
| 8-well chamber slides | Sarstedt, Nümbrecht, Germany | 94.6170.802 |
| Lipofectamine 3000 transfection reagent | Invitrogen, Waltham, USA | L3000001 |
| PureCube 100 Ni-NTA Agarose beads | Cube Biotech GmbH, Manheim, Germany | 74103 |
| Cabbage phospholipase D (PLD) | Sigma-Aldrich, St. Louis, USA | P8398 |
| Poly-D-lysin | Sigma-Aldrich, St. Louis, USA | A-003-M |
| LysoTracker Red DND-99 | Invitrogen, Waltham, USA | L7528 |
| LysoSensor™ Green DND-189 | Invitrogen, Waltham, USA | L7535 |
| HCS LipidTOX™ Green phospholipidosis detection reagent | Invitrogen, Waltham, USA | H34350 |
| Hoechst 3342 | Abcam, Cambridge, United Kingdom | ab228551 |
| Nile red | Sigma-Aldrich, St. Louis, USA | 72485 |
| 17:1 LPG | Avanti Polar Lipids, Alabaster, USA | 858127 |
| 14:0-14:0 BMP | Avanti Polar Lipids, Alabaster, USA | 857131 |
| 14:0-14:0-14:0 hemi-BMP | Avanti Polar Lipids, Alabaster, USA | 857132 |
| 14:0-14:0 PG | Avanti Polar Lipids, Alabaster, USA | 840445 |
| 17:0-17:0 PE | Avanti Polar Lipids, Alabaster, USA | 830756 |
| 17:0-17:0 PS | Avanti Polar Lipids, Alabaster, USA | 840028 |
| 17:1 LPC | Avanti Polar Lipids, Alabaster, USA | 855677 |
| 17:1 LPE | Avanti Polar Lipids, Alabaster, USA | 856707 |
| 17:1 LPS | Avanti Polar Lipids, Alabaster, USA | 858141 |
| 17:0-17:0 PC | Larodan, Solana, Sweden | 37-170 |
| *sn*-1-oleoyl-R,*rac* LPG (18:1 LPG) | Avanti Polar Lipids, Alabaster, USA | 858125 |
| *sn*-3,3'-18:1-18:1-*S*,*S* BMP | Avanti Polar Lipids, Alabaster, USA | 857135 |
| 16:0-CoA | Avanti Polar Lipids, Alabaster, USA | 870716 |
| 18:1-CoA | Avanti Polar Lipids, Alabaster, USA | 870719 |
| 18:2-CoA | Avanti Polar Lipids, Alabaster, USA | 870736 |
| GPE | Cayman Chemical, Ann Arbor, USA | Cay34460 |
| GPC | Sigma-Aldrich, St. Louis, USA | G5291 |
| Bafilomycin A1 | MedChemExpress, Monmouth Junction, USA | HY-100558 |
| Triacsin C | Sigma-Aldrich, St. Louis, USA | T4540 |
| Lauryl maltose neopentyl glycol (LMNG) | Thermo Fisher Scientific, Waltham, USA | A50940 |
| Torin1 | Sigma-Aldrich, St. Louis, USA | 475991 |
| $^{13}$C-Glycerin | Sigma-Aldrich, St. Louis, USA | 489476 |
| $^{13}$C-UL-D-Glucose | Sigma-Aldrich, St. Louis, USA | 389374 |
| Color Prestained Protein Standard Broad Range | New England Biolabs, Ipswich, USA | P7719S |
| Q5 site-directed mutagenesis kit | New England Biolabs, Ipswich, USA | E0552S |
| Low protein binding tube | Thermo Fisher Scientific, Waltham, USA | 90410 |
| Pierce™ Anti-HA-magnetic beads | Thermo Fisher Scientific, Waltham, USA | 88837 |
| EasyEights™ EasySep™ magnet | STEMCELL Technologies, Vancouver, Canada | 18103 |

**Supplementary table 2. Plasmids**

| Plasmid | Addgene ID | Coding sequence cloned into the expression vector |
|---|---|---|
| pcDNA3.1(+)/myc-HisC-h*CLN8* | - | >NM_018941.4 Homo sapiens CLN8 transmembrane ER and ERGIC protein (CLN8), mRNA<br>ATGAATCCTGCGAGCGATGGGGGCACATCAGAGAGCATTTTTGACCTGGACTATGCATC<br>CTGGGGGATCCGCTCCACGCTGATGGTCGCTGGCTTTGTCTTCTACTTGGGCGTCTTTGT<br>GGTCTGCCACCAGCTGTCCTCTTCCCTGAATGCCACTTACCGTTCTTTGGTGGCCAGAGA<br>GAAGGTCTTCTGGGACCTGGCGGCCACGCGTGCAGTCTTTGGTGTTCAGAGCACAGCCG<br>CAGGCCTGTGGGCTCTGCTGGGGGACCCTGTGCTGCATGCCGACAAGGCGCGTGGCCAG<br>CAGAACTGGTGCTGGTTTCACATCACGACAGCAACGGGATTCTTTTGCTTTGAAAATGTT<br>GCAGTCCACCTGTCCAACTTGATCTTCCGGACATTTGACTTGTTTCTGGTTATCCACCATC<br>TCTTTGCCTTTCTTGGGTTTCTTGGCTGCTTGGTCAATCTCCAAGCTGGCCACTATCTAGC<br>TATGACCACGTTGCTCCTGGAGATGAGCACGCCCTTTACCTGCGTTTCCTGGATGCTCTT<br>AAAGGCGGGCTGGTCCGAGTCTCTGTTTTGGAAGCTCAACCAGTGGCTGATGATTCACA<br>TGTTTCACTGCCGCATGGTTCTAACCTACCACATGTGGTGGGTGTGTTTCTGGCACTGGG<br>ACGGCCTGGTCAGCAGCCTGTATCTGCCTCATTTGACACTGTTCCTTGTCGGACTGGCTC<br>TGCTTACGCTAATCATTAATCCATATTGGACCCATAAGAAGACTCAGCAGCTTCTCAATC<br>CGGTGGACTGGAACTTCGCACAGCCAGAAGCCAAGAGCAGGCCAGAAGGCAACGGGCA<br>GCTGCTGCGGAAGAAGAGGCCA |
| pcDNA3.1(+)/myc-HisC-h*CLN8*-R204C created by site directed mutagenesis Mutation is highlighted in **bold** | - | ATGAATCCTGCGAGCGATGGGGGCACATCAGAGAGCATTTTTGACCTGGACTATGCATC<br>CTGGGGGATCCGCTCCACGCTGATGGTCGCTGGCTTTGTCTTCTACTTGGGCGTCTTTGT<br>GGTCTGCCACCAGCTGTCCTCTTCCCTGAATGCCACTTACCGTTCTTTGGTGGCCAGAGA<br>GAAGGTCTTCTGGGACCTGGCGGCCACGCGTGCAGTCTTTGGTGTTCAGAGCACAGCCG<br>CAGGCCTGTGGGCTCTGCTGGGGGACCCTGTGCTGCATGCCGACAAGGCGCGTGGCCAG<br>CAGAACTGGTGCTGGTTTCACATCACGACAGCAACGGGATTCTTTTGCTTTGAAAATGTT<br>GCAGTCCACCTGTCCAACTTGATCTTCCGGACATTTGACTTGTTTCTGGTTATCCACCATC<br>TCTTTGCCTTTCTTGGGTTTCTTGGCTGCTTGGTCAATCTCCAAGCTGGCCACTATCTAGC<br>TATGACCACGTTGCTCCTGGAGATGAGCACGCCCTTTACCTGCGTTTCCTGGATGCTCTT<br>AAAGGCGGGCTGGTCCGAGTCTCTGTTTTGGAAGCTCAACCAGTGGCTGATGATTCACA<br>TGTTTCACTGC**T**GCATGGTTCTAACCTACCACATGTGGTGGGTGTGTTTCTGGCACTGGG<br>ACGGCCTGGTCAGCAGCCTGTATCTGCCTCATTTGACACTGTTCCTTGTCGGACTGGCTC<br>TGCTTACGCTAATCATTAATCCATATTGGACCCATAAGAAGACTCAGCAGCTTCTCAATC<br>CGGTGGACTGGAACTTCGCACAGCCAGAAGCCAAGAGCAGGCCAGAAGGCAACGGGCA<br>GCTGCTGCGGAAGAAGAGGCCA |
| pcDNA3.1(+)/myc-HisC-h*CLN5* | - | >NM_006493.4:19-1095 Homo sapiens CLN5 intracellular trafficking protein (CLN5), transcript variant 1, mRNA<br>ATGGCGCAGGAGGTAGACACGGCACAGGGCGCCGAGATGCGGCGGGGCGCGGGCGCG<br>GCTCGGGGACGCGCTTCCTGGTGCTGGGCCCTGGCGCTGCTTTGGCTCGCGGTGGTTCCG<br>GGCTGGTCCCGGGTCTCGGGCATCCCCTCCCGGCGCCACTGGCCGGTGCCCTACAAGCG<br>CTTTGACTTCCGTCCAAAACCTGATCCTTATTGTCAAGCTAAGTATACTTTCTGTCCAACT<br>GGCTCACCTATCCCAGTTATGGAGGGTGATGATGACATTGAAGTTTTTCGATTACAAGCC<br>CCAGTATGGGAATTTAAATATGGAGACCTCCTGGGACACTTGAAAATTATGCATGATGC<br>CATTGGATTCAGAAGTACATTAACTGGCAAGAACTACACAATGGAATGGTATGAACTTT<br>TCCAACTTGGCAACTGTACATTTCCCCATCTCCGACCTGAAATGGATGCCCCTTTCTGGT<br>GTAATCAAGGCGCTGCCTGCTTTTTTGAGGGAATTGATGATGTTCACTGGAAGGAAAAT<br>GGGACATTAGTTCAAGTAGCAACTATATCAGGAAACATGTTCAACCAAATGGCAAAGTG<br>GGTGAAACAGGACAATGAAACAGGAATTTATTATGAGACATGGAATGTAAAAGCCAGC<br>CCAGAAAAGGGGGCAGAGACATGGTTTGATTCCTACGACTGTTCCAAATTTGTGTTAAG<br>GACCTTTAACAAGTTGGCTGAATTTGGAGCAGAGTTCAAGAACATAGAAACCAACTATA<br>CAAGAATATTTCTTTACAGTGGAGAACCTACTTATCTGGGAAATGAAACATCTGTTTTTG<br>GGCCAACAGGAAACAAGACTCTTGGTTTAGCCATAAAAAGATTTTATTACCCCTTCAAA<br>CCACATTTGCCAACTAAAGAATTTCTGTTGAGTCTCTTGCAAATTTTTGATGCAGTGATT<br>GTGCACAAACAGTTCTATTTGTTTTATAATTTTGAATATTGGTTTTTACCTATGAAATTCC<br>CTTTTATTAAAATAACATATGAAGAAATCCCTTTACCTATCAGAAACAAAACACTCTCTG<br>GTTTA |
| pcDNA3.1(+)/myc-HisB-*hLPGAT1* | - | >NM_001375841.1:363-1475 Homo sapiens lysophosphatidylglycerol acyltransferase 1 (LPGAT1), transcript variant 3, mRNA<br>ATGGCTATAACTTTGGAAGAAGCTCCGTGGCTGGGCTGGCTCTTGGTGAAAGCACTGAT<br>GAGGTTTGCCTTCATGGTCGTCAACAACCTGGTTGCTATTCCATCCTACATCTGCTATGT<br>AATTATACTTCAGCCCCTTCGAGTGCTGGACAGTAAGCGGTTCTGGTATATCGAAGGAA<br>TCATGTATAAATGGCTTTTAGGAATGGTAGCTTCCTGGGGATGGTATGCTGGATATACA<br>GTGATGGAATGGGGAGAAGATATTAAAGCAGTTTCAAAAGATGAAGCAGTGATGTTGG<br>TGAATCATCAGGCAACAGGAGATGTGTGCACACTGATGATGTGCCTCCAGGACAAAGG<br>ACTGGTTGTTGCTCAGATGATGTGGTTGATGGATCATATTTTTAAGTACACAAACTTTGG<br>AATTGTTTCTCTAGTTCATGGAGACTTCTTTATAAGACAGGGAAGATCTTATCGTGACCA<br>ACAGCTGCTGCTTCTCAAGAAGCACTTAGAAAATAATTACAGGAGCAGAGATCGAAAAT<br>GGATTGTTTTGTTTCCAGAAGGGGGCTTCCTCAGGAAGAGGCGAGAAACAAGTCAGGCA<br>TTTGCCAAGAAAAATAACTTGCCATTTCTTACAAATGTTACTCTGCCAAGGTCTGGGGCA<br>ACAAAAATTATTTTGAATGCACTTGTAGCACAACAGAAAAATGGAAGTCCAGCAGGAG<br>GAGATGCTAAAGAATTAGACAGCAAATCAAAAGGCCTCCAGTGGATAATAGATACAAC<br>GATAGCTTATCCCAAAGCTGAACCTATAGATATTCAAACCTGGATCCTTGGATACAGGA<br>AACCAACAGTCACACATGTACATTACAGGATCTTTCCAATTAAAGATGTACCCCTGGAG<br>ACTGATGACCTTACCACTTGGCTCTATCAGCGGTTTGTTGAAAAAGAAGACCTCTTATCA<br>CATTTTTATGAAACAGGAGCTTTTCCACCTTCCAAGGGCCATAAGGAAGCTGTTTCCAGG<br>GAGATGACCCTCAGCAACTTGTGGATATTTCTCATACAGTCTTTTGCATTTTTGTCAGGC<br>TATATGTGGTACAACATCATTCAGTATTTTTACCATTGCCTGTTT |
| pcDNA4/His-MaxC-*LacZ* | - | |
| pLJC5-Tmem192-3xHA | gift from David Sabatini, Addgene plasmid #102930 | |
| psPAX2 | gift from Didier Trono, Addgene plasmids #12260 | |
| pMD2.G | gift from Didier Trono, Addgene plasmids #12259 | |
| pSpCas9(BB)-2A-Puro (PX459) V2.0 | gift from Feng Zhang, Addgene, plasmid #62988 | |

**Supplementary table 3. Antibodies**

| Antibodies | Company | Catalog number | Dilution* |
|---|---|---|---|
| Anti-6X His tag® antibody | Abcam, Cambridge, United Kingdom | ab18184 | 1:3,000 |
| Anti-HA antibody | Sigma-Aldrich, St. Louis, USA | H3663 | 1:2,000 |
| Rabbit Anti-GAPDH Monoclonal Antibody | Cell Signaling Technology, Danvers, USA | 2118 | 1:10,000 |
| PDI (C81H6) Rabbit mAb | Cell Signaling Technology, Danvers, USA | 3501S | 1:1,000 |
| LAMP1 (C54H11) Rabbit mAb | Cell Signaling Technology, Danvers, USA | 3243 | 1:1,000 |
| ß-Actin Antibody | Cell Signaling Technology, Danvers, USA | 4967 | 1:10,000 |
| Anti-SDHA antibody | Abcam, Cambridge, United Kingdom | ab14715 | 1:1,000 |
| Goat Anti-Rabbit IgG Antibody (H+L), Peroxidase | Vector Laboratories, Newark, USA | PI-1000-1 | 1:10,000 |
| Sheep Anti-Mouse IgG ECL Antibody, HRP Conjugated | Cytiva, Marlborough, USA | NA9310-1ml | 1:10,000 |
| | | *(in tris-buffered saline with tween20 supplemented with 5% blotting grade milk powder | |

**Supplementary table 4.  Targeted MS Precursor-to-Product Ion Transitions**

| Compound name | Precursor (m/z) | Product (m/z) | Fragmentor (V) | CAV (V) | CE (V) | Polarity |
|---|---|---|---|---|---|---|
| IS BMP 28:0 (14:0-14:0) | 665.4 | 227.2 | 250 V | 5 V | 40 V | Neagtive |
| BMP 32:0 (16:0-16:0) | 721.5 | 255.2 | 250 V | 5 V | 40 V | Neagtive |
| BMP 32:0 (16:0-16:0) +2 | 723.5 | 255.2 | 250 V | 5 V | 40 V | Neagtive |
| BMP 32:0 (16:0-16:0) +3 | 724.5 | 255.2 | 250 V | 5 V | 40 V | Neagtive |
| BMP 32:0 (16:0-16:0) +5 | 726.5 | 255.2 | 250 V | 5 V | 40 V | Neagtive |
| BMP 32:1 (16:0-16:1) | 719.5 | 253.2 | 250 V | 5 V | 40 V | Neagtive |
| BMP 34:0 (16:0-18:0) | 749.5 | 283.2 | 250 V | 5 V | 40 V | Neagtive |
| BMP 34:1 (16:0-18:1) | 747.5 | 281.2 | 250 V | 5 V | 40 V | Neagtive |
| BMP 34:1 (16:0-18:1) +2 | 749.5 | 281.2 | 250 V | 5 V | 40 V | Neagtive |
| BMP 34:1 (16:0-18:1) +3 | 750.5 | 281.2 | 250 V | 5 V | 40 V | Neagtive |
| BMP 34:1 (16:0-18:1) +5 | 752.5 | 281.2 | 250 V | 5 V | 40 V | Neagtive |
| BMP 34:2 (16:0-18:2) | 745.5 | 279.2 | 250 V | 5 V | 40 V | Neagtive |
| BMP 36:1 (18:0-18:1) | 775.5 | 283.2 | 250 V | 5 V | 40 V | Neagtive |
| BMP 36:2 (18:1-18:1) | 773.5 | 281.2 | 250 V | 5 V | 40 V | Neagtive |
| BMP 36:2 (18:1-18:1) +2 | 775.5 | 281.2 | 250 V | 5 V | 40 V | Neagtive |
| BMP 36:2 (18:1-18:1) +3 | 776.5 | 281.2 | 250 V | 5 V | 40 V | Neagtive |
| BMP 36:2 (18:1-18:1) +5 | 778.5 | 281.2 | 250 V | 5 V | 40 V | Neagtive |
| BMP 36:3 (18:1-18:2) | 771.5 | 279.2 | 250 V | 5 V | 40 V | Neagtive |
| BMP 36:3 (18:1-18:2) +2 | 773.5 | 279.2 | 250 V | 4 V | 35 V | Neagtive |
| BMP 36:3 (18:1-18:2) +3 | 774.5 | 279.2 | 250 V | 4 V | 35 V | Neagtive |
| BMP 36:3 (18:1-18:2) +5 | 776.5 | 279.2 | 250 V | 4 V | 35 V | Neagtive |
| BMP 36:4 (18:2-18:2) | 769.5 | 279.2 | 250 V | 5 V | 40 V | Neagtive |
| BMP 36:4 (18:2-18:2) +2 | 771.5 | 279.2 | 250 V | 5 V | 40 V | Neagtive |
| BMP 36:4 (18:2-18:2) +3 | 772.5 | 279.2 | 250 V | 5 V | 40 V | Neagtive |
| BMP 36:4 (18:2-18:2) +5 | 774.5 | 279.2 | 250 V | 5 V | 40 V | Neagtive |
| BMP 36:5 (18:2-18:3) | 767.5 | 277.2 | 250 V | 5 V | 40 V | Neagtive |
| BMP 38:4 (18:0-20:4) | 797.5 | 303.2 | 250 V | 5 V | 40 V | Neagtive |
| BMP 38:5 (18:1-20:4) | 795.5 | 303.2 | 250 V | 5 V | 40 V | Neagtive |
| BMP 38:5 (18:1-20:4) +2 | 797.5 | 303.2 | 250 V | 5 V | 40 V | Neagtive |
| BMP 38:5 (18:1-20:4) +3 | 798.5 | 303.2 | 250 V | 5 V | 40 V | Neagtive |
| BMP 38:5 (18:1-20:4) +5 | 800.5 | 303.2 | 250 V | 5 V | 40 V | Neagtive |
| BMP 38:5 (18:2-20:3) | 795.5 | 305.2 | 250 V | 5 V | 40 V | Neagtive |
| BMP 38:6 (16:0-22:6) | 793.5 | 255.2 | 250 V | 5 V | 40 V | Neagtive |
| BMP 38:6 (18:2-20:4) | 793.5 | 303.23 | 250 V | 5 V | 40 V | Neagtive |
| BMP 38:7 (16.1-22:6) | 791.48 | 327.23 | 250 V | 5 V | 40 V | Neagtive |
| BMP 40:5 (18:1-22:4) | 823.5 | 331.3 | 250 V | 5 V | 40 V | Neagtive |
| BMP 40:6 (18:1-22:5) | 821.5 | 329.3 | 250 V | 5 V | 40 V | Neagtive |
| BMP 40:6 (18:2-22:4) | 821.5 | 331.3 | 250 V | 5 V | 40 V | Neagtive |
| BMP 40:7 (18:1-22:6) | 819.5 | 327.2 | 250 V | 5 V | 40 V | Neagtive |
| BMP 40:7 (18:1-22:6) +2 | 821.5 | 327.2 | 250 V | 5 V | 40 V | Neagtive |
| BMP 40:7 (18:1-22:6) +3 | 822.5 | 327.2 | 250 V | 5 V | 40 V | Neagtive |
| BMP 40:7 (18:1-22:6) +5 | 824.5 | 327.2 | 250 V | 5 V | 40 V | Neagtive |
| BMP 40:7 (18:2-22.5) | 819.5 | 329.2 | 250 V | 5 V | 40 V | Neagtive |
| BMP 40:8 (20:4-20:4) | 817.5 | 327.2 | 250 V | 5 V | 40 V | Neagtive |
| BMP 40:8 (20:4-20:4) +2 | 819.5 | 327.2 | 250 V | 5 V | 40 V | Neagtive |
| BMP 40:8 (20:4-20:4) +3 | 820.5 | 327.2 | 250 V | 5 V | 40 V | Neagtive |
| BMP 40:8 (20:4-20:4) +5 | 822.5 | 327.2 | 250 V | 5 V | 40 V | Neagtive |
| BMP 40:9 (18:3-22:6) | 815.48 | 277.3 | 250 V | 5 V | 40 V | Neagtive |
| BMP 42:10 (20:4-22:6) | 841.5 | 327.23 | 250 V | 5 V | 40 V | Neagtive |

| | | | | | | |
|---|---|---|---|---|---|---|
| BMP 42:9 (20:3-22:6) | 843.5 | 327.2 | 250 V | 5 V | 40 V | Neagtive |
| BMP 44:10 (22:4-22:6) | 869.5 | 331.2 | 250 V | 5 V | 40 V | Neagtive |
| BMP 44:11 (22:5-22:6) | 867.5 | 329.2 | 250 V | 5 V | 40 V | Neagtive |
| BMP 44:12 (22:6-22:6) | 865.5 | 327.2 | 250 V | 5 V | 40 V | Neagtive |
| BMP 44:12 (22:6-22:6) +2 | 867.5 | 327.2 | 250 V | 5 V | 40 V | Neagtive |
| BMP 44:12 (22:6-22:6) +3 | 868.5 | 327.2 | 250 V | 5 V | 40 V | Neagtive |
| BMP 44:12 (22:6-22:6) +5 | 870.5 | 327.2 | 250 V | 5 V | 40 V | Neagtive |
| CL 70:3 | 1430 | 1430 | 110 V | 5 V | 10 V | Neagtive |
| CL 70:3 +2 | 1432 | 1432 | 110 V | 5 V | 10 V | Neagtive |
| CL 70:3 +3 | 1433 | 1433 | 110 V | 5 V | 10 V | Neagtive |
| CL 70:3 +4 | 1434 | 1434 | 110 V | 5 V | 10 V | Neagtive |
| CL 70:3 +5 | 1435 | 1435 | 110 V | 5 V | 10 V | Neagtive |
| CL 70:3 +6 | 1436 | 1436 | 110 V | 5 V | 10 V | Neagtive |
| CL 70:3 +7 | 1437 | 1437 | 110 V | 5 V | 10 V | Neagtive |
| CL 70:3 +8 | 1438 | 1438 | 110 V | 5 V | 10 V | Neagtive |
| CL 70:3 +9 | 1439 | 1439 | 110 V | 5 V | 10 V | Neagtive |
| CL 70:4 | 1427.9 | 1427.9 | 110 V | 5 V | 10 V | Neagtive |
| CL 70:4 +2 | 1429.9 | 1429.9 | 110 V | 5 V | 10 V | Neagtive |
| CL 70:4 +3 | 1430.9 | 1430.9 | 110 V | 5 V | 10 V | Neagtive |
| CL 70:4 +4 | 1431.9 | 1431.9 | 110 V | 5 V | 10 V | Neagtive |
| CL 70:4 +5 | 1432.9 | 1432.9 | 110 V | 5 V | 10 V | Neagtive |
| CL 70:4 +6 | 1433.9 | 1433.9 | 110 V | 5 V | 10 V | Neagtive |
| CL 70:4 +7 | 1434.9 | 1434.9 | 110 V | 5 V | 10 V | Neagtive |
| CL 70:4 +8 | 1435.9 | 1435.9 | 110 V | 5 V | 10 V | Neagtive |
| CL 70:4 +9 | 1436.9 | 1436.9 | 110 V | 5 V | 10 V | Neagtive |
| CL 70:5 | 1425.9 | 1425.9 | 110 V | 5 V | 10 V | Neagtive |
| CL 70:5 +2 | 1427.9 | 1427.9 | 110 V | 5 V | 10 V | Neagtive |
| CL 70:5 +3 | 1428.9 | 1428.9 | 110 V | 5 V | 10 V | Neagtive |
| CL 70:5 +4 | 1429.9 | 1429.9 | 110 V | 5 V | 10 V | Neagtive |
| CL 70:5 +5 | 1430.9 | 1431.9 | 110 V | 5 V | 10 V | Neagtive |
| CL 70:5 +6 | 1431.9 | 1430.9 | 110 V | 5 V | 10 V | Neagtive |
| CL 70:5 +7 | 1432.9 | 1432.9 | 110 V | 5 V | 10 V | Neagtive |
| CL 70:5 +8 | 1433.9 | 1433.9 | 110 V | 5 V | 10 V | Neagtive |
| CL 70:5 +9 | 1434.9 | 1434.9 | 110 V | 5 V | 10 V | Neagtive |
| CL 72:3 | 1458 | 1458 | 110 V | 5 V | 10 V | Neagtive |
| CL 72:3 +2 | 1460 | 1460 | 110 V | 5 V | 10 V | Neagtive |
| CL 72:3 +3 | 1461 | 1461 | 110 V | 5 V | 10 V | Neagtive |
| CL 72:3 +4 | 1462 | 1462 | 110 V | 5 V | 10 V | Neagtive |
| CL 72:3 +5 | 1463 | 1463 | 110 V | 5 V | 10 V | Neagtive |
| CL 72:3 +6 | 1464 | 1464 | 110 V | 5 V | 10 V | Neagtive |
| CL 72:3 +7 | 1465 | 1465 | 110 V | 5 V | 10 V | Neagtive |
| CL 72:3 +8 | 1466 | 1466 | 110 V | 5 V | 10 V | Neagtive |
| CL 72:3 +9 | 1467 | 1467 | 110 V | 5 V | 10 V | Neagtive |
| CL 72:4 | 1456 | 1456 | 110 V | 5 V | 10 V | Neagtive |
| CL 72:4 +2 | 1458 | 1458 | 110 V | 5 V | 10 V | Neagtive |
| CL 72:4 +3 | 1459 | 1459 | 110 V | 5 V | 10 V | Neagtive |
| CL 72:4 +4 | 1460 | 1460 | 110 V | 5 V | 10 V | Neagtive |
| CL 72:4 +5 | 1461 | 1461 | 110 V | 5 V | 10 V | Neagtive |
| CL 72:4 +6 | 1462 | 1462 | 110 V | 5 V | 10 V | Neagtive |
| CL 72:4 +7 | 1463 | 1463 | 110 V | 5 V | 10 V | Neagtive |
| CL 72:4 +8 | 1464 | 1464 | 110 V | 5 V | 10 V | Neagtive |
| CL 72:4 +9 | 1465 | 1465 | 110 V | 5 V | 10 V | Neagtive |
| CL 72:5 | 1454 | 1454 | 110 V | 5 V | 10 V | Neagtive |
| CL 72:5 +2 | 1456 | 1456 | 110 V | 5 V | 10 V | Neagtive |
| CL 72:5 +3 | 1457 | 1457 | 110 V | 5 V | 10 V | Neagtive |
| CL 72:5 +4 | 1458 | 1458 | 110 V | 5 V | 10 V | Neagtive |
| CL 72:5 +5 | 1459 | 1459 | 110 V | 5 V | 10 V | Neagtive |
| CL 72:5 +6 | 1460 | 1460 | 110 V | 5 V | 10 V | Neagtive |
| CL 72:5 +7 | 1461 | 1461 | 110 V | 5 V | 10 V | Neagtive |
| CL 72:5 +8 | 1462 | 1462 | 110 V | 5 V | 10 V | Neagtive |
| CL 72:5 +9 | 1463 | 1463 | 110 V | 5 V | 10 V | Neagtive |
| IS Hemi BMP 42:0 (14:0-14:0-14:0) | 875.6 | 227.2 | 250 V | 4 V | 44 V | Neagtive |
| Hemi-BMP 52:2 (16:0-18:1-18:1) | 1011.8 | 281.2 | 250 V | 4 V | 44 V | Neagtive |
| Hemi-BMP 52:2 (16:0-18:1-18:1) +2 | 1013.8 | 281.2 | 250 V | 4 V | 44 V | Neagtive |
| Hemi-BMP 52:2 (16:0-18:1-18:1) +3 | 1014.8 | 281.2 | 250 V | 4 V | 44 V | Neagtive |
| Hemi-BMP 52:2 (16:0-18:1-18:1) +5 | 1016.8 | 281.2 | 250 V | 4 V | 44 V | Neagtive |
| Hemi-BMP 54:2 (18:0-18:1-18:1) | 1039.8 | 281.2 | 250 V | 4 V | 44 V | Neagtive |
| Hemi-BMP 54:2 (18:0-18:1-18:1) +2 | 1041.8 | 281.2 | 250 V | 4 V | 44 V | Neagtive |
| Hemi-BMP 54:2 (18:0-18:1-18:1) +3 | 1042.8 | 281.2 | 250 V | 4 V | 44 V | Neagtive |
| Hemi-BMP 54:2 (18:0-18:1-18:1) +5 | 1044.8 | 281.2 | 250 V | 4 V | 44 V | Neagtive |
| Hemi-BMP 54:3 (18:1-18:1-18:1) | 1037.7 | 281.2 | 250 V | 4 V | 44 V | Neagtive |
| Hemi-BMP 54:3 (18:1-18:1-18:1) +2 | 1039.7 | 281.2 | 250 V | 4 V | 44 V | Neagtive |
| Hemi-BMP 54:3 (18:1-18:1-18:1) +3 | 1040.7 | 281.2 | 250 V | 4 V | 44 V | Neagtive |

| | | | | | | |
|---|---|---|---|---|---|---|
| Hemi-BMP 54:3 (18:1-18:1-18:1) +5 | 1042.7 | 281.2 | 250 V | 4 V | 44 V | Neagtive |
| Hemi-BMP 54:4 (18:2-18:1-18:1) | 1035.76 | 281.2 | 250 V | 4 V | 44 V | Neagtive |
| LPG 16:0 | 483.3 | 255.2 | 180 V | 4 V | 28 V | Neagtive |
| LPG 16:0 +2 | 485.3 | 255.2 | 180 V | 4 V | 28 V | Neagtive |
| LPG 16:0 +3 | 486.3 | 255.2 | 180 V | 4 V | 28 V | Neagtive |
| LPG 16:0 +5 | 488.3 | 255.2 | 180 V | 4 V | 28 V | Neagtive |
| LPG 16:1 | 481.2 | 253.2 | 180 V | 4 V | 28 V | Neagtive |
| LPG 16:1 +2 | 483.2 | 253.2 | 180 V | 4 V | 28 V | Neagtive |
| LPG 16:1 +3 | 484.2 | 253.2 | 180 V | 4 V | 28 V | Neagtive |
| LPG 16:1 +5 | 486.2 | 253.2 | 180 V | 4 V | 28 V | Neagtive |
| IS LPG 17:1 | 495.2 | 267.2 | 180 V | 4 V | 28 V | Neagtive |
| LPG 18:0 | 511.3 | 283.2 | 180 V | 4 V | 28 V | Neagtive |
| LPG 18:1 | 509.3 | 281.2 | 180 V | 4 V | 28 V | Neagtive |
| LPG 18:1 +2 | 511.3 | 281.2 | 180 V | 4 V | 28 V | Neagtive |
| LPG 18:1 +3 | 512.3 | 281.2 | 180 V | 4 V | 28 V | Neagtive |
| LPG 18:1 +5 | 514.3 | 281.2 | 180 V | 4 V | 28 V | Neagtive |
| LPG 18:1 +3 (MS2 MAG +0) | 514.3 | 339.3 | 130 V | 4 V | 16 V | Positive |
| LPG 18:1 +3 (MS2 MAG +3) | 514.3 | 342.3 | 130 V | 4 V | 16 V | Positive |
| LPG 18:2 | 507.3 | 279.2 | 180 V | 4 V | 28 V | Neagtive |
| LPG 18:2 +2 | 509.3 | 279.2 | 180 V | 4 V | 28 V | Neagtive |
| LPG 18:2 +3 | 510.3 | 279.2 | 180 V | 4 V | 28 V | Neagtive |
| LPG 18:2 +5 | 512.3 | 279.2 | 180 V | 4 V | 28 V | Neagtive |
| LPG 20:4 | 531.3 | 303.2 | 180 V | 4 V | 28 V | Neagtive |
| LPG 20:4 +2 | 533.3 | 303.2 | 180 V | 4 V | 28 V | Neagtive |
| LPG 20:4 +3 | 534.3 | 303.2 | 180 V | 4 V | 28 V | Neagtive |
| LPG 20:4 +5 | 536.3 | 303.2 | 180 V | 4 V | 28 V | Neagtive |
| LPG 22:6 | 555.3 | 327.2 | 180 V | 4 V | 28 V | Neagtive |
| LPG 22:6 +2 | 557.3 | 327.2 | 180 V | 4 V | 28 V | Neagtive |
| LPG 22:6 +3 | 558.3 | 327.2 | 180 V | 4 V | 28 V | Neagtive |
| LPG 22:6 +5 | 560.3 | 327.2 | 180 V | 4 V | 28 V | Neagtive |
| IS PG 28:0 (14:0-14:0) | 665.4 | 227.2 | 250 V | 4 V | 45 V | Neagtive |
| PG 32:0 (16:0-16:0) | 721.5 | 255.2 | 250 V | 4 V | 45 V | Neagtive |
| PG 32:0 (16:0-16:0) +2 | 723.5 | 255.2 | 250 V | 4 V | 45 V | Neagtive |
| PG 32:0 (16:0-16:0) +3 | 724.5 | 255.2 | 250 V | 4 V | 45 V | Neagtive |
| PG 32:0 (16:0-16:0) +5 | 726.5 | 255.2 | 250 V | 4 V | 45 V | Neagtive |
| PG 32:1 (16:0-16:1) | 719.48 | 253.2 | 250 V | 4 V | 45 V | Neagtive |
| IS PG 34:0 (17:0-17:0) | 749.5 | 269.2 | 250 V | 4 V | 45 V | Neagtive |
| PG 34:1 (16:0-18:1) | 747.5 | 255.2 | 250 V | 4 V | 45 V | Neagtive |
| PG 34:1 (16:0-18:1) +2 | 749.5 | 255.2 | 250 V | 4 V | 45 V | Neagtive |
| PG 34:1 (16:0-18:1) +3 | 750.5 | 255.2 | 250 V | 4 V | 45 V | Neagtive |
| PG 34:1 (16:0-18:1) +5 | 752.5 | 255.2 | 250 V | 4 V | 45 V | Neagtive |
| PG 34:2 (16:0-18:2) | 745.5 | 255.2 | 250 V | 4 V | 45 V | Neagtive |
| PG 34:2 (16:1-18:1) | 745.5 | 281.2 | 250 V | 4 V | 45 V | Neagtive |
| PG 34:3 (16:1-18:2) | 743.48 | 253.2 | 250 V | 4 V | 45 V | Neagtive |
| PG 36:1(18:0-18:1) | 775.5 | 281.2 | 250 V | 4 V | 45 V | Neagtive |
| PG 36:1(18:0-18:1) +2 | 777.5 | 281.2 | 250 V | 4 V | 45 V | Neagtive |
| PG 36:1(18:0-18:1) +3 | 778.5 | 281.2 | 250 V | 4 V | 45 V | Neagtive |
| PG 36:1(18:0-18:1) +5 | 780.5 | 281.2 | 250 V | 4 V | 45 V | Neagtive |
| PG 36:2 (18:1-18:1) | 773.5 | 281.3 | 250 V | 4 V | 45 V | Neagtive |
| PG 36:2 (18:1-18:1) +2 | 775.5 | 281.3 | 250 V | 4 V | 45 V | Neagtive |
| PG 36:2 (18:1-18:1) +3 | 776.5 | 281.3 | 250 V | 4 V | 45 V | Neagtive |
| PG 36:2 (18:1-18:1) +5 | 778.5 | 281.3 | 250 V | 4 V | 45 V | Neagtive |
| PG 36:3 (18:1-18:2) | 771.5 | 281.28 | 250 V | 4 V | 45 V | Neagtive |
| PG 36:4 (16:0-20:4) | 769.5 | 255.23 | 250 V | 4 V | 45 V | Neagtive |
| PG 38:5 (18:1-20:4) | 795.5 | 281.2 | 250 V | 4 V | 45 V | Neagtive |
| PG 38:5 (18:1-20:4) +2 | 797.5 | 281.2 | 250 V | 4 V | 45 V | Neagtive |
| PG 38:5 (18:1-20:4) +3 | 798.5 | 281.2 | 250 V | 4 V | 45 V | Neagtive |
| PG 38:5 (18:1-20:4) +5 | 800.5 | 281.2 | 250 V | 4 V | 45 V | Neagtive |
| DAG 34:1-18:1 | 612.6 | 339 | 110 V | 5 V | 16 V | Positive |
| DAG 34:1-18:1 + 2 | 614.6 | 341 | 110 V | 5 V | 16 V | Positive |
| DAG 34:1-18:1 + 3 | 615.6 | 342 | 110 V | 5 V | 16 V | Positive |
| DAG 36:1-18:1 | 640.6 | 339 | 110 V | 5 V | 16 V | Positive |
| DAG 36:1-18:1 + 2 | 642.6 | 341 | 110 V | 5 V | 16 V | Positive |
| DAG 36:1-18:1 + 3 | 643.6 | 342 | 110 V | 5 V | 16 V | Positive |
| DAG 36:2-18:1 | 638.6 | 339 | 110 V | 5 V | 16 V | Positive |
| DAG 36:2-18:1 +2 | 640.6 | 340 | 110 V | 5 V | 16 V | Positive |
| DAG 36:2-18:1 +3 | 641.6 | 341 | 110 V | 5 V | 16 V | Positive |
| DAG 36:2-18:2 | 638.6 | 337 | 110 V | 5 V | 16 V | Positive |
| DAG 36:2-18:2 +2 | 640.6 | 339 | 110 V | 5 V | 16 V | Positive |
| DAG 36:2-18:2 +3 | 641.6 | 400 | 110 V | 5 V | 16 V | Positive |
| DAG 38:4-20:4 | 662.6 | 361 | 110 V | 5 V | 16 V | Positive |
| DAG 38:4-20:4 +2 | 664.6 | 363 | 110 V | 5 V | 16 V | Positive |
| DAG 38:4-20:4 +3 | 665.6 | 364 | 110 V | 5 V | 16 V | Positive |

| | | | | | | |
|---|---|---|---|---|---|---|
| IS LPE 17:1 | 479.3 | 325.2 | 111 V | 5 V | 16 V | Positive |
| IS DG 34:0-17:0 | 614.6 | 327.3 | 137 V | 5 V | 20 V | Positive |
| IS LPC 17:1 | 508.3 | 184.1 | 79 V | 5 V | 28 V | Positive |
| IS LPS 17:1 | 510.3 | 325.2 | 121 V | 5 V | 20 V | Positive |
| IS PC 28:0 | 678.5 | 184.1 | 164 V | 5 V | 28 V | Positive |
| IS PC 34:0 | 762.6 | 184.1 | 204 V | 5 V | 32 V | Positive |
| IS PE 34:0 | 720.6 | 579.5 | 131 V | 5 V | 20 V | Positive |
| IS PS 34:0 | 764.6 | 579.5 | 131 V | 5 V | 20 V | Positive |
| LPC 16:0 | 496.3 | 184.1 | 179 V | 5 V | 28 V | Positive |
| LPC 16:0 +2 | 498.3 | 184.1 | 179 V | 5 V | 28 V | Positive |
| LPC 16:0 +3 | 499.3 | 184.1 | 179 V | 5 V | 28 V | Positive |
| LPC 16:1 | 494.3 | 184.1 | 179 V | 5 V | 28 V | Positive |
| LPC 16:1 +2 | 496.3 | 184.1 | 179 V | 5 V | 28 V | Positive |
| LPC 16:1 +3 | 497.3 | 184.1 | 179 V | 5 V | 28 V | Positive |
| LPC 18:0 | 524.3 | 184.1 | 179 V | 5 V | 28 V | Positive |
| LPC 18:0 +2 | 526.3 | 184.1 | 179 V | 5 V | 28 V | Positive |
| LPC 18:0 +3 | 527.3 | 184.1 | 179 V | 5 V | 28 V | Positive |
| LPC 18:1 | 522.3 | 184.1 | 179 V | 5 V | 28 V | Positive |
| LPC 18:1 +2 | 524.3 | 184.1 | 179 V | 5 V | 28 V | Positive |
| LPC 18:1 +3 | 525.3 | 184.1 | 179 V | 5 V | 28 V | Positive |
| LPC 20:4 | 544.3 | 184.1 | 179 V | 5 V | 28 V | Positive |
| LPC 20:4 +2 | 546.3 | 184.1 | 179 V | 5 V | 28 V | Positive |
| LPC 20:4 +3 | 547.3 | 184.1 | 179 V | 5 V | 28 V | Positive |
| LPE 16:0 | 454.3 | 313.3 | 111 V | 5 V | 16 V | Positive |
| LPE 16:0 +2 | 456.3 | 315.3 | 111 V | 5 V | 16 V | Positive |
| LPE 16:0 +3 | 457.3 | 316.3 | 111 V | 5 V | 16 V | Positive |
| LPE 16:1 | 452.3 | 311.3 | 111 V | 5 V | 16 V | Positive |
| LPE 16:1 +2 | 454.3 | 313.3 | 111 V | 5 V | 16 V | Positive |
| LPE 16:1 +3 | 455.3 | 314.3 | 111 V | 5 V | 16 V | Positive |
| LPE 18:0 | 482.3 | 341.3 | 111 V | 5 V | 16 V | Positive |
| LPE 18:0 +2 | 484.3 | 343.3 | 111 V | 5 V | 16 V | Positive |
| LPE 18:0 +3 | 485.3 | 344.3 | 111 V | 5 V | 16 V | Positive |
| LPE 18:1 | 480.3 | 339.3 | 111 V | 5 V | 16 V | Positive |
| LPE 18:1 +2 | 482.3 | 341.3 | 111 V | 5 V | 16 V | Positive |
| LPE 18:1 +3 | 483.3 | 342.3 | 111 V | 5 V | 16 V | Positive |
| LPE 20:4 | 502.3 | 361.3 | 111 V | 5 V | 16 V | Positive |
| LPE 20:4 +2 | 504.3 | 363.3 | 111 V | 5 V | 16 V | Positive |
| LPE 20:4 +3 | 505.3 | 364.3 | 111 V | 5 V | 16 V | Positive |
| LPS 16:0 | 498.3 | 313.3 | 121 V | 5 V | 20 V | Positive |
| LPS 16:0 +2 | 500.3 | 315.3 | 121 V | 5 V | 20 V | Positive |
| LPS 16:0 +3 | 501.3 | 316.3 | 121 V | 5 V | 20 V | Positive |
| LPS 16:1 | 496.3 | 311.3 | 121 V | 5 V | 20 V | Positive |
| LPS 16:1 +2 | 498.3 | 313.3 | 121 V | 5 V | 20 V | Positive |
| LPS 16:1 +3 | 499.3 | 314.3 | 121 V | 5 V | 20 V | Positive |
| LPS 18:0 | 526.3 | 341.3 | 121 V | 5 V | 20 V | Positive |
| LPS 18:0 +2 | 528.3 | 343.3 | 121 V | 5 V | 20 V | Positive |
| LPS 18:0 +3 | 529.3 | 344.3 | 121 V | 5 V | 20 V | Positive |
| LPS 18:1 | 524.3 | 339.3 | 121 V | 5 V | 20 V | Positive |
| LPS 18:1 +2 | 526.3 | 341.3 | 121 V | 5 V | 20 V | Positive |
| LPS 18:1 +3 | 527.3 | 342.3 | 121 V | 5 V | 20 V | Positive |
| PC 32:1 | 732.6 | 184.1 | 180 V | 5 V | 30 V | Positive |
| PC 32:1 +2 | 734.6 | 184.1 | 180 V | 5 V | 30 V | Positive |
| PC 32:1 +3 | 735.6 | 184.1 | 180 V | 5 V | 30 V | Positive |
| PC 34:1 | 760.6 | 184.1 | 180 V | 5 V | 30 V | Positive |
| PC 34:1 +2 | 762.6 | 184.1 | 180 V | 5 V | 30 V | Positive |
| PC 34:1 +3 | 763.6 | 184.1 | 180 V | 5 V | 30 V | Positive |
| PC 34:2 | 758.6 | 184.1 | 180 V | 5 V | 30 V | Positive |
| PC 34:2 +2 | 760.6 | 184.1 | 180 V | 5 V | 30 V | Positive |
| PC 34:2 +3 | 761.6 | 184.1 | 180 V | 5 V | 30 V | Positive |
| PC 36:1 | 788.6 | 184.1 | 180 V | 5 V | 30 V | Positive |
| PC 36:1 +2 | 790.6 | 184.1 | 180 V | 5 V | 30 V | Positive |
| PC 36:1 +3 | 791.6 | 184.1 | 180 V | 5 V | 30 V | Positive |
| PC 36:2 | 786.6 | 184.1 | 180 V | 5 V | 30 V | Positive |
| PC 36:2 +2 | 788.6 | 184.1 | 180 V | 5 V | 30 V | Positive |
| PC 36:2 +3 | 789.6 | 184.1 | 180 V | 5 V | 30 V | Positive |
| PE 34:1 | 718.5 | 577.5 | 131 V | 5 V | 20 V | Positive |
| PE 34:1 +2 | 720.5 | 579.5 | 131 V | 5 V | 20 V | Positive |
| PE 34:1 +3 | 721.5 | 580.5 | 131 V | 5 V | 20 V | Positive |
| PE 36:1 | 746.6 | 605.6 | 131 V | 5 V | 20 V | Positive |
| PE 36:1 +2 | 748.6 | 607.6 | 131 V | 5 V | 20 V | Positive |
| PE 36:1 +3 | 749.6 | 608.6 | 131 V | 5 V | 20 V | Positive |
| PE 36:2 | 744.6 | 603.6 | 131 V | 5 V | 20 V | Positive |
| PE 36:2 +2 | 746.6 | 605.6 | 131 V | 5 V | 20 V | Positive |

| | | | | | | |
|---|---|---|---|---|---|---|
| PE 36:2 +3 | 747.6 | 606.6 | 131 V | 5 V | 20 V | Positive |
| PE 38:4 | 768.6 | 627.6 | 131 V | 5 V | 20 V | Positive |
| PE 38:4 +2 | 770.6 | 629.6 | 131 V | 5 V | 20 V | Positive |
| PE 38:4 +3 | 771.6 | 630.6 | 131 V | 5 V | 20 V | Positive |
| PE 38:5 | 766.6 | 625.6 | 131 V | 5 V | 20 V | Positive |
| PE 38:5 +2 | 768.6 | 627.6 | 131 V | 5 V | 20 V | Positive |
| PE 38:5 +3 | 769.6 | 628.6 | 131 V | 5 V | 20 V | Positive |
| PI 34:1 | 835.5 | 241 | 131 V | 5 V | 50 V | Neagtive |
| PI 34:1 +2 | 837.5 | 241 | 131 V | 5 V | 50 V | Neagtive |
| PI 34:1 +3 | 838.5 | 241 | 131 V | 5 V | 50 V | Neagtive |
| PI 36:2 | 861.5 | 241 | 131 V | 5 V | 50 V | Neagtive |
| PI 36:2 +2 | 863.5 | 241 | 131 V | 5 V | 50 V | Neagtive |
| PI 36:2 +3 | 864.5 | 241 | 131 V | 5 V | 50 V | Neagtive |
| PI 36:4 | 857.5 | 241 | 131 V | 5 V | 50 V | Neagtive |
| PI 36:4 +2 | 859.5 | 241 | 131 V | 5 V | 50 V | Neagtive |
| PI 36:4 +3 | 860.5 | 241 | 131 V | 5 V | 50 V | Neagtive |
| PI 38:4 | 885.5 | 241 | 131 V | 5 V | 50 V | Neagtive |
| PI 38:4 +2 | 887.5 | 241 | 131 V | 5 V | 50 V | Neagtive |
| PI 38:4 +3 | 888.5 | 241 | 131 V | 5 V | 50 V | Neagtive |
| PS 34:1 | 762.5 | 577.5 | 131 V | 5 V | 20 V | Positive |
| PS 34:1 +2 | 764.5 | 579.5 | 131 V | 5 V | 20 V | Positive |
| PS 34:1 +3 | 765.5 | 580.5 | 131 V | 5 V | 20 V | Positive |
| PS 36:1 | 790.6 | 605.6 | 131 V | 5 V | 20 V | Positive |
| PS 36:1 +2 | 792.6 | 607.6 | 131 V | 5 V | 20 V | Positive |
| PS 36:1 +3 | 793.6 | 608.6 | 131 V | 5 V | 20 V | Positive |
| PS 36:2 | 788.6 | 603.6 | 131 V | 5 V | 20 V | Positive |
| PS 36:2 +2 | 790.6 | 605.6 | 131 V | 5 V | 20 V | Positive |
| PS 36:2 +3 | 791.6 | 606.6 | 131 V | 5 V | 20 V | Positive |
| PS 38:1 | 818.8 | 633.8 | 131 V | 5 V | 20 V | Positive |
| PS 38:1 +2 | 820.8 | 635.8 | 131 V | 5 V | 20 V | Positive |
| PS 38:1 +3 | 821.8 | 636.8 | 131 V | 5 V | 20 V | Positive |
| IS GPC +3 | 261.1 | 86.1 | 130 V | 5 V | 37 V | Positive |
| GPG | 245.1 | 153.1 | 140 V | 5 V | 13 V | Neagtive |
| GPG | 245.1 | 79 | 140 V | 5 V | 45 V | Neagtive |
| GPG +3 | 248.1 | 79 | 140 V | 5 V | 45 V | Neagtive |
| GPG +5 | 250.1 | 79 | 140 V | 5 V | 45 V | Neagtive |
